# Development of Integrin α5β1-targeted PET/NIR imaging probes for glioblastoma intraoperative navigation and intracavity targeted radionuclide therapy

**DOI:** 10.64898/2026.01.09.698741

**Authors:** Siqi Zhang, Jieting Shen, Yiming Li, Xuekai Song, Wenhao Liu, Hongyi Huang, Jiang Wu, Can Liu, Mengjie Yang, Liyuan Xu, Dengfu Wu, Zhong Zhang, Feng, Yingzi Zhang, Rui Wang, Kuan Hu

**Affiliations:** State Key Laboratory of Bioactive Substance and Function of Natural Medicines, Institute of Materia Medica, Chinese Academy of Medical Sciences and Peking Union Medical College, No.1 Xian Nong Tan Street, Beijing, 100050, China; Department of Nuclear Medicine, Nanjing First Hospital, Nanjing Medical University, 210006, Nanjing, China; Department of Neurosurgery, Beijing Tiantan Hospital, Capital Medical University, 100070, Beijing, China; Department of Orthopaedics, the Second Affiliated Hospital of Soochow University, Suzhou 215031, China

**Keywords:** Glioblastoma, Integrin α5β1, PET/CT, Intraoperative navigation, Targeted radionuclide therapy

## Abstract

Incomplete extent of tumor resection is the major factor of glioblastoma recurrence, leading to its poor prognosis. Developing therapies that enable the minimal tumor cell remnant during surgical resection is a major challenge to combat this aggressive cancer. This study develops a glioblastoma integrin α5β1-selective peptide binder through series strategy screening and labeling it with ^68^Ga, indocyanine green (ICG), or ^177^Lu for efficacious multimodal treatment. This strategy combines positron emission tomography (PET) imaging for preoperative diagnosis, intraoperative NIR-II fluorescence-guided surgery for maximal tumor resection, and followed by intracavity targeted radionuclide therapy for elimination of residual tumor cells. We identified GR, with five arginine mutation, exhibiting superior integrin α5β1 binding affinity and brain penetration. [^68^Ga]GR demonstrated substantial tumor uptake and prolonged retention time in both mouse models and tissues from glioblastoma patients. The guidance of ICG-GR achieved accurately intraoperative tumor imaging and maximal tumor resection in orthotopic mouse glioblastoma models. Notably, combining intracavity administration of [^177^Lu]GR following ICG-GR guiding resection markedly inhibited tumor growth and reduced possibility of tumor recurrence compared with surgery alone (unguided or ICG-GR–guided) or unguided surgery followed by [^177^Lu]GR. Therefore, we have reported the integrin α5β1-targeted radionuclide and optical agents for maximal elimination of tumor cells during surgical resection in glioblastoma patients.

**HIGHLIGHTS:**

- GR is a glioblastoma-specific peptide with both favorable binding affinity superior brain penetration.
- ICG-GR is a glioblastoma-targeted NIR probe enabling maximal surgical resection.
- ICG-GR-guided surgery with [^177^Lu]GR-based adjuvant therapy allows robust elimination of tumor cells.

## INTRODUCTION

Glioblastoma is an aggressive and common malignant primary brain tumor. The overall prognosis of this aggressive cancer remains poor. Maximal safe surgical resection stands the main challenge during glioblastoma treatment due to the necessary to preserve essential brain functions, resulting in rapid recurrence ^1^. In relapse or recurrent setting, treatment options that prolong overall survival are less well established ^2^. Although multimodality therapies that combined surgery with chemotherapy and radiotherapy have been widely implemented in glioblastoma treatment, their clinical benefits are restricted by insufficient cell-killing, adaptive therapeutic resistance, and severe side effects ^3,4^. Therefore, efficacious therapies allowing maximal tumor removal and minimized tumor cell residue heralds promising to combat glioblastoma and reduce its mortality.

Incorporating surgery with adjuvant therapies that enable powerful tumor cell elimination is critical in improving the prognosis of glioblastoma ^5^. Targeted radionuclide therapy (TRT) enables the precise delivery of radionuclides to tumor cells. The concentrated dose of radiation on cancer cells induces direct cell death without reliance on signaling pathways, making TRT a powerful and efficient cancer therapeutic option ^6^. Therefore, TRT has garnered attention as promising approach for inducing selective radiation-induced cell killing and for treatment of refractory malignant cancers that harvest rare benefit from other therapies ^7–10^. Notably, glioblastoma-targeted TRT potentially serves as efficacious modality for complete eliminating residual tumor cells postoperatively. Moreover, the integration of positron emission tomography (PET) imaging into TRT results in radiotheranostics, allowing precise diagnosis and precise glioblastoma treatment ^11^.

Intraoperative fluorescent imaging has emerging as a valuable tool for precise identification of tumor margins, which have been clinically approved for imaging-guided tumor resection surgery in various cancers ^12–14^. Due to the demand of glioblastoma for maximal safe surgical resection, various fluorescent agents have been studied in neurosurgery for intraoperative navigation. Indocyanine green (ICG) is a FDA-approved fluorescence probe, its efficacy has been reported in glioma and high-grade gliomas ^15^. Alternative probes including 5-ALA is also increasingly used and exhibited satisfied outcomes in clinical trials ^16^. Despite the substantial progress of fluorescence-guided glioblastoma surgery, their clinical utility is limited by poor selectivity and specificity ^15,17–20^. To address this, glioblastoma specific biomarker-targeted fluorescence probe is necessary to improve the accuracy and efficiency of intraoperative tumor imaging and tumor resection ^21^.

Integrin α5β1 is overexpressed in various cancers, including glioblastoma ^22^. This heterodimeric cell surface receptor, composed of α5 and β1 subunits, interacts with extracellular matrix molecules including fibronectin, thereby promoting cell adhesion, which is crucial for tumor progression, angiogenesis, metastasis, and invasion ^23–25^. Herein, inspired by the strengths of radiotheranositcs and intraoperative navigation in oncology, we demonstrate a more efficient glioblastoma therapeutic paradigm that incorporating complementary integrin α5β1-targeted PET/CT/NIR imaging probes followed by intracavity TRT with same peptide binder-based therapeutic counterpart. This approach enables precise preoperative diagnosis, maximal safe intraoperative resection, powerful elimination of residual tumor cells, potentially reducing the possibility of recurrence. Developing of targeting ligands with high binding affinity and satisfied specificity to integrin α5β1 is essential. Peptides exhibited favorable molecular weight and biomarker recognize ability, making them suitable binders for precise brain delivery. Integrin α5β1-targeted peptide binders have demonstrated potential in PET/CT imaging, including the ^68^Ga-labeled peptidomimetics and the ^68^Ga-labelling N-methylated isoDGR peptides, both of which demonstrate high and specific uptake in α5β1-overexpressed tumors ^26–29^. Despite these, the α5β1-targeted peptide PR_b includes the primary and synergy binding site of for α5 and β1 subunit, which is connected by a linker sequence (GS)_5_ to mimic the distance between two binding site in fibronectin, showing potential in specific and robust binding capacity ^30–32^. The ^18^F-labeling PR_b has presented selective tumor uptake, but the rapid degradation and short tumor retention time hamper its clinical therapeutic potential, particularly for TRT^33,34^. Moreover, PR_b demonstrates poor blood-brain barrier (BBB) penetration, making it unsuitable for imaging of glioblastoma. To overcome these limitations, we modified PR_b based on multiple chemical strategies, including cyclization, PEGylated multivalency, amino acid mutation, and covalency, to enhance the peptide’s capacity on targeting efficacy and BBB penetration.

In this study, we screen the optimized integrin α5β1-targeted peptide binder with superior binding affinity and brain tumor uptake through multi-generation modification. The candidate was labeled with ^68^Ga for preoperative glioblastoma PET/CT imaging, with ICG for fluorescence-guided resection, which was followed by the intracavity TRT using ^177^Lu-labeled counterpart. We determined the theranostic efficacy of the integrin α5β1-targeted combined therapy in subcutaneous and orthotopic glioblastoma mouse models. Importantly, the PET/CT/NIR probe were evaluated in tumor tissues from glioblastoma patients, allowing to indicate its clinical values. Therefore, this study provides a proof-of-concept for a new therapeutic paradigm incorporating glioblastoma-targeted PET/CT imaging, intraoperative navigation, and TRT, enabling the maximal safe surgical resection and efficient residual cell elimination, which is critical for improving the prognosis of glioblastoma.

## RESULT

### Design of series integrin α5β1-targeted radioligands and their *in vitro* evaluation

The integrin α5β1-targeted peptide PR_b (defined as GS) was designed by mimicking the binding epitope of fibronectin to α5β1 ^30,31^, which is a favorable ligand for α5β1-selective radioligand. To further enhance the *in vivo* performance of GS for α5β1-targeted and binding ability, four strategies was introduced (**Figure 1A**). Cyclization possesses evident advantages regarding improved stability and binding ability. Covalently linking of the sidechains of lysine and glutamic acid induce α-helix, potential to enhance the cell membrane permeability and binding affinity ^35^. Subsequently, varying number and order of lysine and glutamic acid was added to GS for generating different cyclic peptides (Cyc-1, Cyc-2, Cyc-3, and Cyc-4). PEGylated multivalency serves as critical approach to enhance the protein binding. Therefore, PEGylated monomer (PEG-1), dimer (PEG-2), and trimer (PEG-3) radioligands were synthesized through multi-arm PEG linker, which may contribute to the improved *in vivo* stability and prolonged circulation time. Specific warheads could covalently link to specific amino acids on proteins, enhancing the binding of peptides to targeted protein. Three covalent warheads, Acrylamide (Alk), Chloroacetamide (CL), and Sulfonyl Fluoride (SF), were also added and covalent peptides were designed. Moreover, the spacer sequence (GS_5_) was designed to mimic distance between the binding sites of α5 and β1 subunits and fibronectin. We assume that alteration of the side chain of amino acids in spacer sequence do not influence their binding site distance. Additionally, replacement of various amino acid contributes to the modification of physical and chemical properties, which in turn improve their pharmacokinetics and pharmacodynamics. Therefore, five serine in linker sequence was mutated to aspartic acid (GD), glutamic acid (GE), or arginine (GR). For further evaluate the effect of arginine, we synthesis peptide with one arginine mutation, which named as R (**Figure S1-S33**).

**Figure 1.**
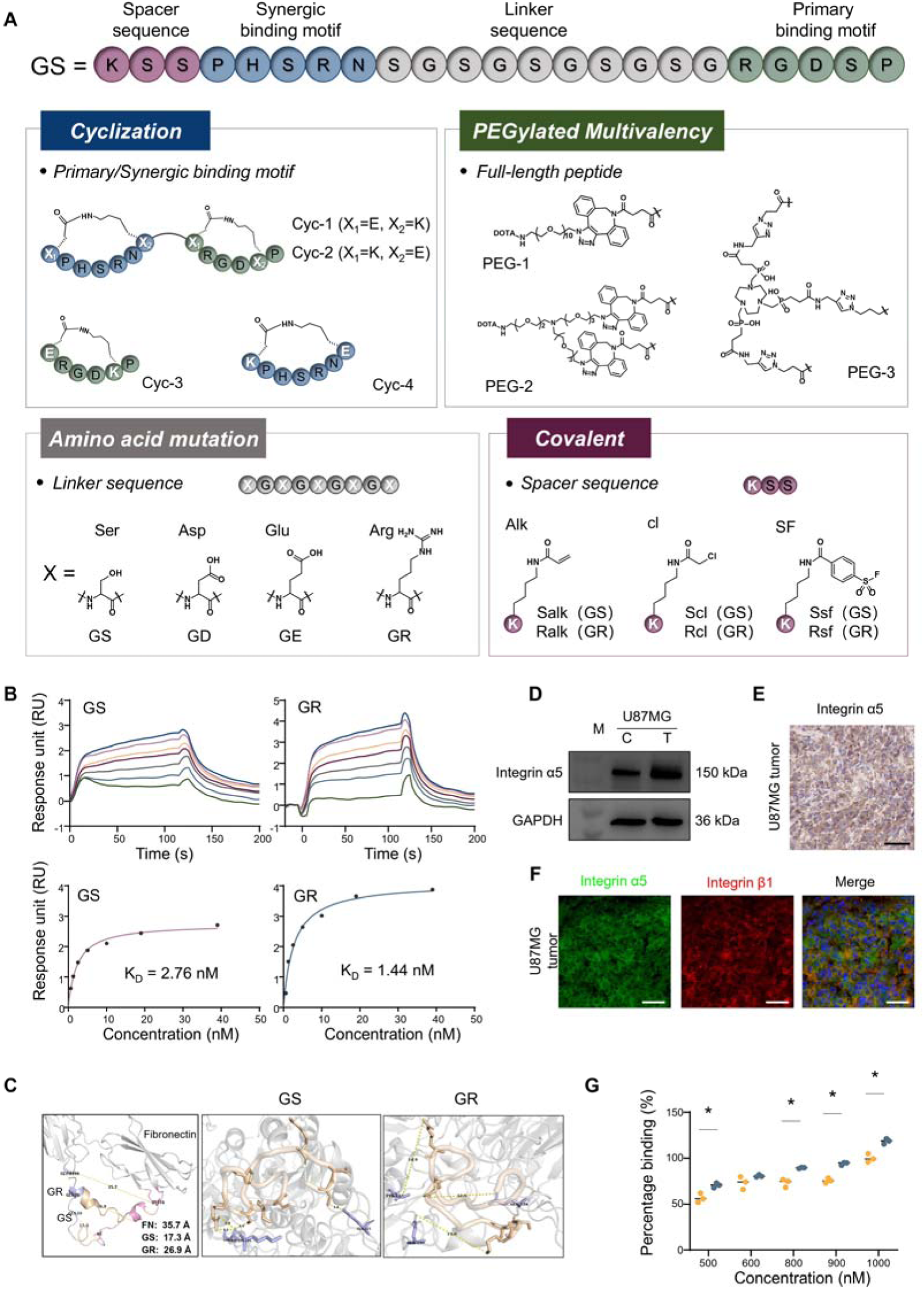
Design scheme and *in vitro* analysis of series integrin α5β1-targeted peptides. **(A)** Strategies for modification of GS, including amino acid mutation, covalent modification, cyclization, and multivalency. **(B)** SPR sensorgrams demonstrating the binding affinity of GS and GR for human integrin α5β1 in a concentration-dependent manner. The equilibrium dissociation constant (K_D_) of each peptide was calculated based on SPR measurements. The K_D_ values of each precursor are shown. **(C)** Molecular docking of GR and GS binding integrin a5b1 protein (grey; PDB: 7NWL) showing the selected possible ligation residues. **(D-F)** Analysis of integrin α5β1 expression in U87MG cells and tumor tissues by western blot (**D**), and immunohistochemistry (**E-F**) analysis. The band for integrin α5 was approximately 150 kDa. M, marker. C, cell. T, tumor. For immunofluorescence images, green is for integrin α5, red for integrin β1, and blue for nucleus. Scale bar, 50□μm (**E-F**). **(G)** *In vitro* cellular uptake of [^68^Ga]GS and [^68^Ga]GR in U87MG cell lines. All results are expressed as means ± SEM, as indicated in at least three independent experiments. “*” represents differences compared with the [^68^Ga]GS. * *p* < 0.05.

We next measured the integrin α5β1 affinity of each peptide through SPR method. Peptides involving cyclization, PEGylated multivalency, and covalent showed reduced binding affinity compared to GS, especially for PEGylated multivalency series, the K_D_ of PEG-1, PEG-2, and PEG-3 were higher than 1 μM. The results demonstrated that all 5 peptides regarding amino acid mutation exhibited high integrin α5β1 affinities, suggesting that the spacer modification did not influence the binding capacity of peptides to integrin α5β1 (**Table S1**). Furthermore, GR exhibited improved binding affinity compared to GS, which is K_D_ = 1.44 nM vs. 2.76 nM (**Figure 1B**). To investigate the binding pattern of amino acid mutation peptides to integrin α5β1, subsequently, we employed AlphaFold3 to predict their three-dimensional structures (**Figure 1C**). Peptides GS exhibited disordered coil configurations, while certain sequences of peptides GR presented alpha-helical structures. We further predicted the binding modes of GS and GR with integrin α5β1 proteins through molecular docking (**Figure 1C**). The analysis revealed that GR shares structural similarities with the natural ligand of integrin α5β1, fibronectin. Compared to the structurally flexible of GS, the two alpha-helical segments of GR confer greater structural stability, facilitating its binding to the active sites of integrin α5β1. Notably, the length of linker sequence for GR (25.9 Å) was closer to the nature distance of two recognized sites on fibronectin (35.7 Å) than GS (17.3 Å).

The enhanced binding affinity of GR inspired us to investigate their cellular uptake when labeling with ^68^Ga, which achieved satisfied radiochemical yield (**Table S1**). Human glioblastoma cell lines U87MG are used to determine the cellular uptake and *in vivo* performance of each radioligands. We confirmed the high expression of integrin α5β1 in U87MG cells and tumors and validated their elevated expression in animal tumor tissues (**Figure 1D-1F**). Incubation of [^68^Ga]GS and [^68^Ga]GR of varying concentration with U87MG cells showed concentration-dependent pattern. Importantly, [^68^Ga]GR showed elevated cell uptake compared to [^68^Ga]GS, indicating its higher binding affinity (**Figure 1G**).

We next investigate the membrane permeability of peptides. The flow cytometry and confocal analysis are performed using Cy5-labeled peptides (**Figure S34**). Due to the low binding affinity of PEGylated multivalency series peptides, peptide regarding cyclization, covalent and amino acid mutation are rolling to permeability determination. Covalent modification of GS or GR neither improve their cellular binding. Notably, the percentage of positive cells for Cy5-Cyc-4 and Cy5-GR treated cells was significantly increased compared with Cy5-GS. They exhibited the strongest cellular uptake capacity, reaching a Cy5-positive rate of 97% after a 4-h incubation (**Figure S34A**). Additionally, the mean fluorescence intensity of Cy5-positive cells showed distinct tendency, with lower Cy5-Cyc-4 cellular uptake than Cy5-GR (**Figure S34B**). Confocal imaging was therefore performed to further visualize their binding patterns. Cy5-GR and Cy5-Cyc-4 treated U87MG cells exhibited greater intracellular signals than Cy5-GS (**Figure S34C**). Furthermore, although comparable positive signals were observed in Cy5-GR and Cy5-Cyc-4 treated cells within 12 h post-injection, Cy5-Cyc-4 was decreased at 18 h and 24 h post-injection, which was consistent with the flow cytometry results (**Figure S34C**). We also found evident cell binding of Cy5-Scl and Cy5-Rcl, while the signals mainly located on the cell membrane (**Figure S34C**). Covalent binding of integrin α5β1 may hamper the cell permeability of peptides, thereby unsuitable for BBB penetration. Because of five arginine in the spacer sequence, Cy5-GR is more lipophobic and shows improved membrane permeability, which endow it prolonged retention time inside the U87MG cells. Therefore, among these peptides, GR exhibits superior binding affinity and cell penetration, which may allow the favorable *in vivo* performance.

### [^68^Ga]GR exhibits highest tumor uptake and prolonged retention time *in vivo*

We further investigate the *in vivo* performance of each peptide through PET/CT imaging. Peptides are labeled with ^68^Ga and satisfied RCY are achieved **(Table S1, Figure S35-S38**). Following intravenous injection of 3.7 MBq of ^68^Ga-labeled peptides per mouse, PET/CT imaging was performed at 0.5 h, 1 h, 2 h, and 4 h post-injection. No significant increasing was observed within 240 min post-injection among radiotracers involving PEGylated multivalency, cyclization, and covalent strategy (**Figure 2A-2E**). These data are consistent with the binding affinity results, further suggesting that these strategies may not suitable for promoting both binding capacity and tumor uptake of [^68^Ga]GS. We next investigate the *in vivo* performance of radioligands regarding amino acid mutation, [^68^Ga]GS, [^68^Ga]GD, [^68^Ga]GE, [^68^Ga]R, and [^68^Ga]GR (**Figure 2F-2I**). The PET/CT imaging revealed that [^68^Ga]GR exhibited improved tumor uptake at each time points post-injection, whereas [^68^Ga]GD, [^68^Ga]GE and [^68^Ga]R showed comparable tumor uptake with [^68^Ga]GS. Furthermore, based on the superior binding ability of GR, we designed conjugated covalent warhead on GR, yielding Ralk, Rcl, and Rsf. Nevertheless, the covalently modified GR did not elevate the tumor uptake. It is worth to note that [^68^Ga]Ralk, [^68^Ga]Rcl, and [^68^Ga]Rsf showed reduced tumor uptake than [^68^Ga]GR (**Figure 2F-2G**). The covalent linking may restrain the binding of peptides to another binding site, which resulting in the reduced integrin α5β1 binding. Taken together, PET/CT imaging in integrin α5β1-overexpression tumor-bearing mice demonstrates that [^68^Ga]GR, with mutation of serine to arginine in linker sequence, exhibited facilitate tumor uptake and tumor retention time compared to [^68^Ga]GS, which could be considered as candidate for the further intraoperative fluorescent-guided surgery and TRT.

**Figure 2.**
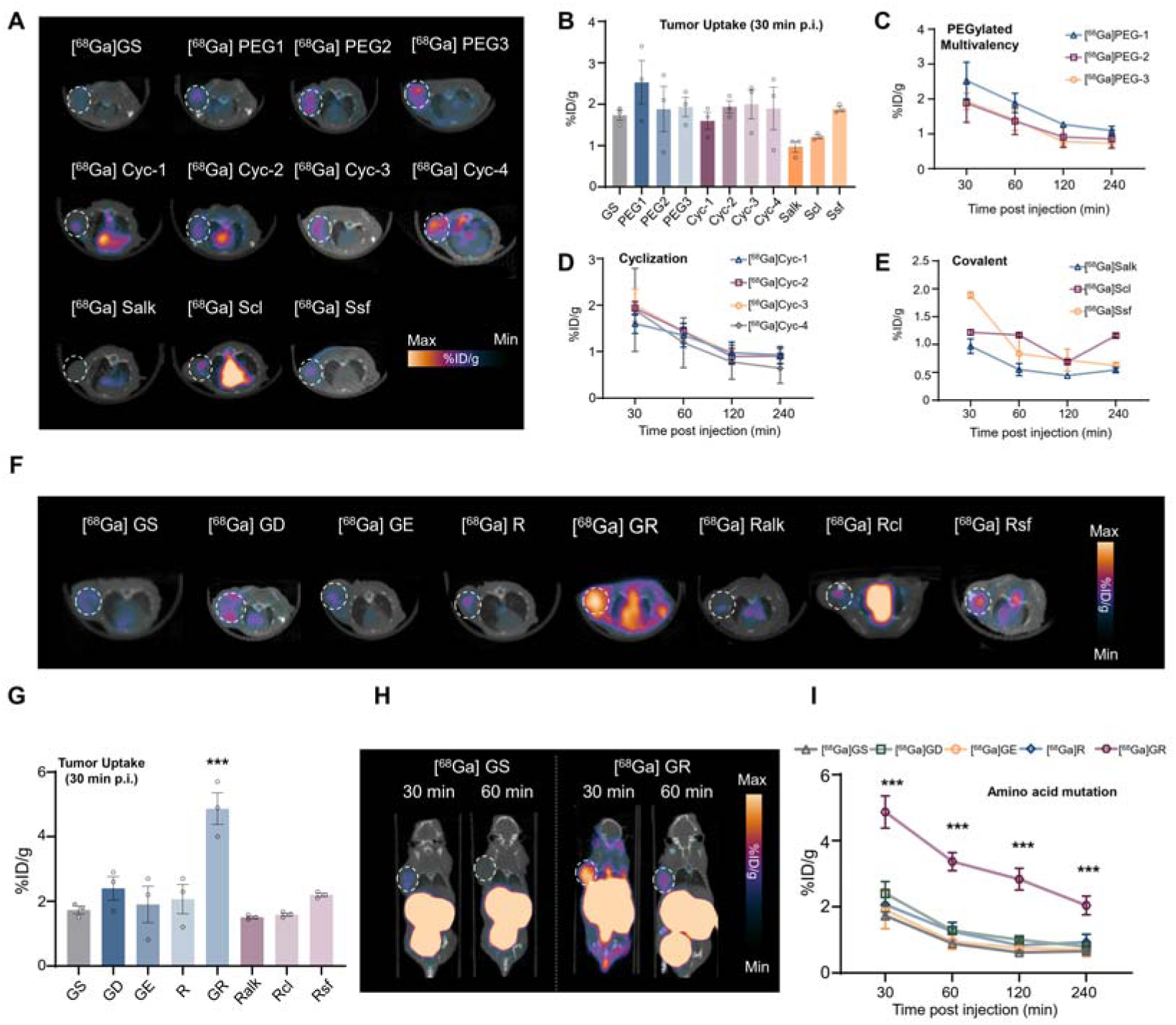
PET/CT imaging of ^68^Ga-labeling peptides in U87MG tumor-bearing mice. **(A)** Series PET/CT images of ^68^Ga-labeling peptides (cyclization, multivalency, and covalent of GS) in U87MG tumor-bearing mice at 30 min post injection. A total of 3.7 MBq of each radiotracer was intravenously injected, and n = 3 mice for each group. The white dotted lines depict the outlines of tumors. **(B)** Tumor uptake of each radiotracer in (**A**) 30 min post-injection. **(C-E)** Time-dependent tumor uptake of each radiotracer in (**A**) within 240 min post-injection. **(F)** PET/CT images of ^68^Ga-labeling peptides (amino acid mutation and covalent of GR) in U87MG tumor-bearing mice at 30 min post injection. A total of 3.7 MBq of each radiotracer was intravenously injected, and n = 3 mice for each group. The white dotted lines depict the outlines of tumors. **(G)** Tumor uptake of each radiotracer in (**F**) 30 min post-injection. **(H)** PET/CT images of [^68^Ga]GS and [^68^Ga]GR in U87MG tumor-bearing mice 30 and 60 min post-injection. **(I)** Time-dependent tumor uptake of each radiotracer regarding amino acid mutation within 240 min post-injection. All results are expressed as means ± SEM, as indicated in at least three independent experiments. A multiple t-test was used when two groups were compared. The symbol “*” represents differences compared with the [^68^Ga]GS. *** *p* < 0.001.

Additionally, *ex vivo* biodistribution studies and dynamic PET/CT imaging of [^68^Ga]GS and [^68^Ga]GR in the U87MG tumor-bearing mice were performed. The results were consistent with the static PET/CT imaging. The uptake of [^68^Ga]GR in the tumor significantly higher than that observed in [^68^Ga]GS (**Figure S39**), which reached 11.8 %ID/g at 5 min post-injection in *ex vivo* biodistribution study. The normal mice *ex vivo* biodistribution and dynamic analysis demonstrate distinct pharmacokinetics between [^68^Ga]GS and [^68^Ga]GR. [^68^Ga]GR exhibited increased liver uptake than [^68^Ga]GS, which may be caused by the improved lipophilic of [^68^Ga]GR, consistent with the PET/CT imaging in tumor-bearing mice (**Figure S40-S43**).

To further investigate the tumor retention time of GR, ^64^Cu with longer half-life (12.7 h) was labeled and yielding [^64^Cu]GS and [^64^Cu]GR. PET/CT imaging revealed that the tumor uptake of [^64^Cu]GR was increasing 24 hours post-injection. On the contrary, [^64^Cu]GS exhibited weak tumor uptake, which was nearly to detected after 4 h of injection. These results were consistent with [^68^Ga]GS and [^68^Ga]GR and further demonstrated the superior tumor uptake and integrin α5β1 binding of GR (**Figure S44**).

### GR could bind integrin α5β1 overexpressed human glioblastoma tissues

Pan-cancer analysis of integrin α5β1 expression based on the Cancer Genome Atlas Program (TCGA) database data revealed that integrin α5β1 expression was universally present in human glioblastoma multiforme (glioblastoma), head and neck squamous cell carcinoma (HNSC), pancreatic adenocarcinoma (PAAD). The comparative analysis of 163 glioblastoma samples and 207 normal tissue samples demonstrated that the expression of integrin α5β1 in glioblastoma tissues was significantly higher than in normal tissues (**Figure 3A**). We collected six tumor samples from glioblastoma patients for histological and expression analysis (**Figure 3B-3E**). WB and IHC results revealed substantial integrin α5β1 expression on human glioblastoma tissues, and varying expression level was founded (**Figure 3C-3E**). Patient 1 and patient 2 exhibited highest expression, whereas patient 3 and patient 6 has weak integrin α5β1 on the tumor cells. Patient 4 and patient 5 showed medium expression level. The integrin α5β1 level on glioblastoma cells was associated with the cancer grade.

**Figure 3.**
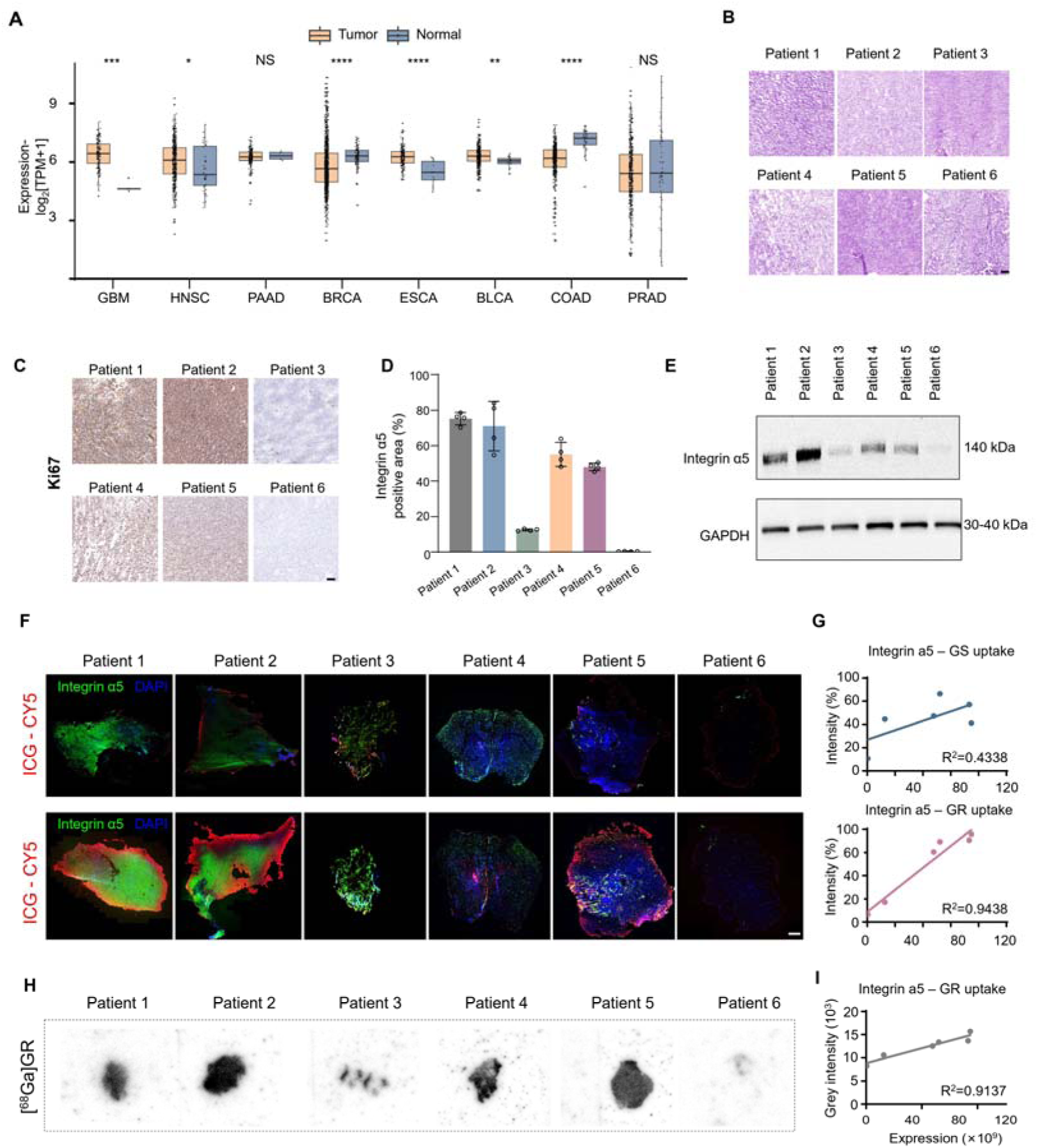
*Ex vivo* evaluation of GS and GR in glioblastoma patient tumors. **(A)** The medium expression of ITGA5 in human cancer tissues or normal tissues. Data were collected from The Cancer Genome Atlas Program (TCGA). GBM, glioblastoma multiforme. HNSC, head and neck squamous cell carcinoma. PAAD, pancreatic adenocarcinoma. BRCA, breast invasive carcinoma. ESCA, esophageal carcinoma. BLCA, bladder urothelial carcinoma. COAD, colon adenocarcinoma. PRAD, prostate adenocarcinoma. **(B)** H&E staining of six glioblastoma patient tumors. Scale bar, 100 μm. **(C-D)** Integrin α5 immunohistochemical images (**C**) and quantification (**D**) of six glioblastoma patient tumors. Scale bar, 100 μm. **(E)** Western blot analysis of integrin α5 expression in six glioblastoma patient tumors. **(F)** Immunofluorescence images of brain tissue sections from six glioblastoma patient tumors treated with Cy5-GS and ICG-GR. Green indicates integrin α5, Red for GS or GR, and blue for nucleus. Scale bar, 1 mm. **(G)** Correlation analysis of integrin α5 expression and GS (bottom) and GR (below) uptake in six glioblastoma patient tumors. **(H)** Autoradiography imaging of six glioblastoma patient tumors when incubated with [68Ga]GR. **(I)** Correlation analysis of integrin α5 expression and GR (below) uptake in six glioblastoma patient tumors. All results are expressed as means ± SEM, as indicated in at least three independent experiments. A multiple t-test was used when two groups were compared. The symbol “*” represents differences compared with the normal tissue. NS, non-significant, * p < 0.05, ** p < 0.01. **** p < 0.0001.

*Ex vivo* staining of the six human tumor tissue sections by both ICG-GR and [^68^Ga]GR was performed to evaluate the binding of GR to human glioblastoma. For fluorescence imaging, we also staining the integrin α5 to determine the specificity of GR binding (**Figure 3F**). ICG-GS exhibited poor binding to human glioblastoma tissues. Moreover, the binding of ICG-GS showed weak correlation with integrin α5, indicating the insufficient targeting and selectivity of GS (**Figure 3F-3G**). In contrast with GS, ICG-GR demonstrated evident binding to human glioblastoma tissues. The binding of GR showed strong positive correlation with integrin α5 (**Figure 3F-3G**). Patient 1 and patient 2 with high integrin α5 level exhibited robust ICG-GR signals, which was overlapped with integrin α5 protein. However, we found that several integrin α5 positive cells were not labeled with ICG-GR, which may be caused by the heterogeneity expression pattern of integrin α5 in each individual. To further determine the binding of GR, autoradiography was performed to investigate the binding of [^68^Ga]GR to human glioblastoma tissues (**Figure 3H-3I**). The results showed that the uptake of [^68^Ga]GR was high in glioblastoma patient tumor tissues, which was also proportional to the expression of integrin α5β1, further indicating that GR holds superior binging affinity and specificity to human glioblastoma integrin α5β1. Therefore, these data demonstrate the potential of GR on clinical glioblastoma diagnosis and treatment, suggesting the clinical value of GR-based intraoperative navigation and postoperative TRT combination therapy.

### GR shows enhanced brain penetration in both healthy and orthotopic glioblastoma mice

Sufficient ability to cross the BBB enable efficient orthotopic glioblastoma imaging and α5β1-targeted intraoperative florescent-guided surgery. Transwell assays was first performed to investigate the BBB cross of [^68^Ga]GS, [^68^Ga]GR, [^68^Ga]Cyc-3, [^68^Ga]Cyc-4, [^68^Ga]Scl, and [^68^Ga]Rcl (**Figure 4A-4C**), which exhibited improved tumor uptake or membrane penetration (**Figure 2 and Figure S34**). After added each radiotracer to the insert for 4 h, the radioactivity of upper and lower culture medium were measured. [^68^Ga]GR, [^68^Ga]Cyc-3, and [^68^Ga]Cyc-4 showed reduced radioactivity of upper culture medium, which was higher in lower medium (**Figure 4B**). Higher cellular uptake of [^68^Ga]GR in U87MG cells that cultured at the lower layer demonstrate the enhanced cell binding and internalization of GR (**Figure 4C**). Although [^68^Ga]Cyc-3 and [^68^Ga]Cyc-4 exhibited improved ability to across the cerebral microvascular endothelial cells and human brain microvascular pericytes (HBVPs), a major part of BBB, the weak α5β1 binding limited the cell uptake by glioblastoma cells (**Figure 4C**). The uptake in BBB organoids also confirmed the superior BBB penetration and glioblastoma binding ability of [^68^Ga]GR (**Figure 4D**).

**Figure 4.**
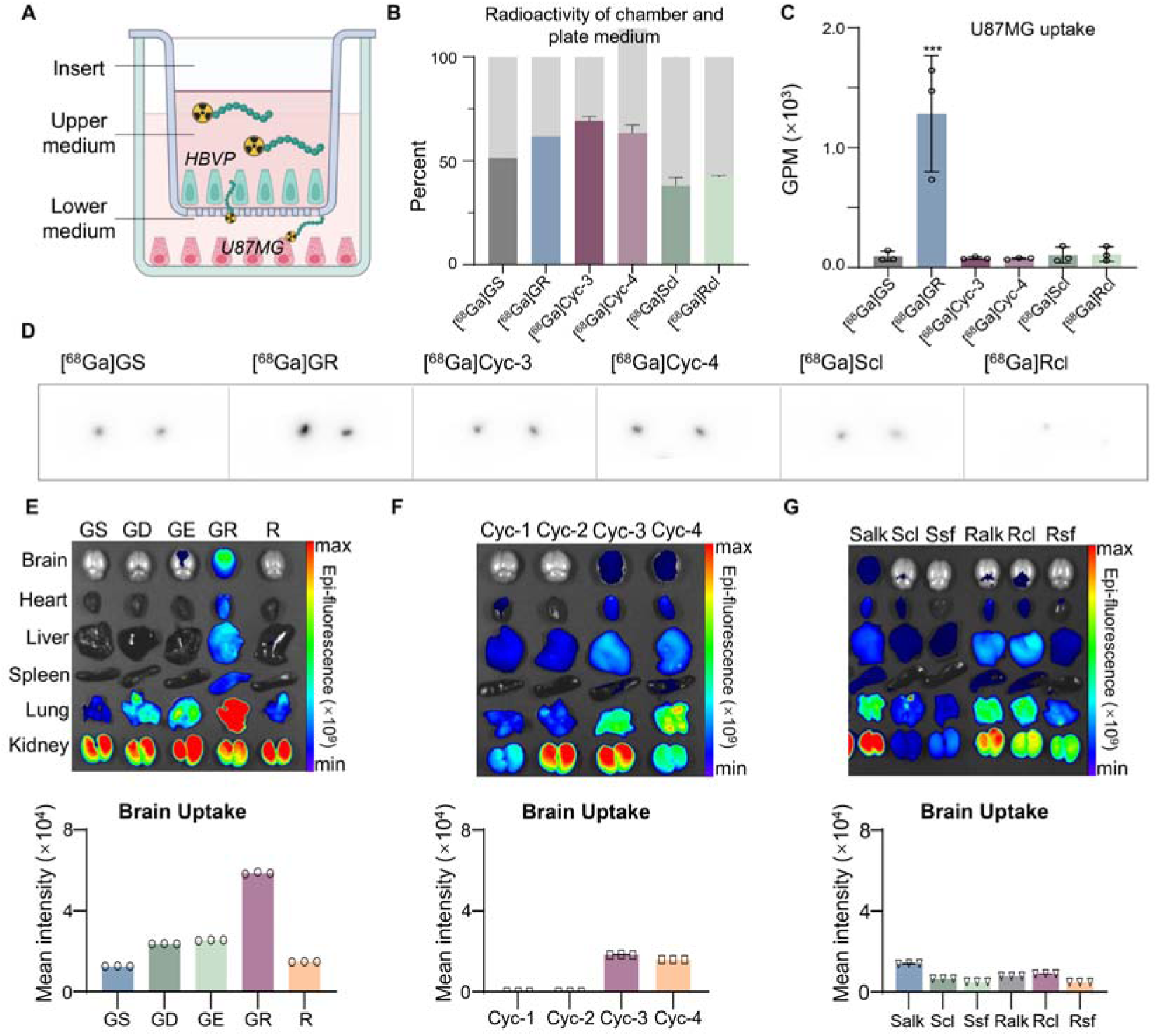
Evaluation of BBB permeability of each peptide. **(A)** Scheme of transwell assay for evaluation of the BBB permeability of 68Ga-labeled peptides. **(B)** The qualification of radioactivity in the bottom and below of transwell for each radiotracer. **(C)** The U87MG cell uptake of each radiotracer in the transwell assay. **(D)** Evaluation of the BBB permeability of 68Ga-labeled peptides using BBB organoids. Autoradiography was performed to measure the radioactivity of each organoid. **(E)** In vivo fluorescence imaging of Cy5-labeled peptides in normal mice. Representative fluorescence images of normal mice acquired at 1h after intravenous injections with Cy5-conjugated peptides (bottom). Each mouse is administered at a dose of 5 mg/kg. n = 3 for each group. Quantification of fluorescence intensity brains corresponding to imaging. All results are expressed as means ± SEM, as indicated in at least three independent experiments. A multiple t-test was used when two groups were compared. The symbol “*” represents differences compared with the [68Ga]GS. *** p < 0.001.

We further investigate the *in vivo* brain accumulation of each peptide after systemic administration in healthy mice (**Figure 4E and 4G**). The highest brain accumulation of Cy5-GR was observed, which was significantly higher than Cy5-GS. Despite, Cy5-Cyc-3 and Cy5-Cyc-4 also showed improved tumor signals compared to Cy5-GS, which was consistent to the *in vitro* evaluations. Therefore, Cy5-GR demonstrates the superior capacity to cross the BBB, which is also improved for Cy5-Cyc-3 and Cy5-Cyc-4.

Due to the different BBB penetration between brains of healthy and cancer mice, we further investigate the capabilities of the Cy5-GR for imaging orthotopic glioblastoma (**Figure 5**). To generate orthotopic glioblastoma tumor-bearing mice, the U87MG cells with Luc reporter were orthotopically injected into the brain of nude-mice. The staining of integrin α5 (tumor cells), GFAP (astrocytes), and NEUN (neurons) confirmed the successful establish of orthotopic glioblastoma tumor-bearing mice (**Figure 5A**). The *in vivo* imaging of Cy5-GS and Cy5-GR demonstrated that Cy5-GR exhibited higher orthotopic tumor uptake, suggesting better BBB permeability and integrin α5β1 binding (**Figure 5B and 5C**). Quantitative analysis revealed that at 25 min post-injection, the fluorescence intensity at the glioblastoma tumor in Cy5-GR was significantly higher than that in Cy5-GS (**Figure 5D**). To evaluate the uptake of each peptide in cell level, the brains of orthotopic glioblastoma tumor-bearing mice treated with Cy5-GS and Cy5-GR were collected and then sectioned for fluorescence imaging (**Figure 5E-5I**). IHC staining confirmed the strong integrin α5β1 expressed tumor tissues in each brain (**Figure 5E**). The tumor uptake of Cy5-GR was higher than Cy5-GS, which was overlapped with integrin α5β. Furthermore, no significant fluorescence signal was observed in normal brain tissues (**Figure 5F-5I**).

**Figure 5.**
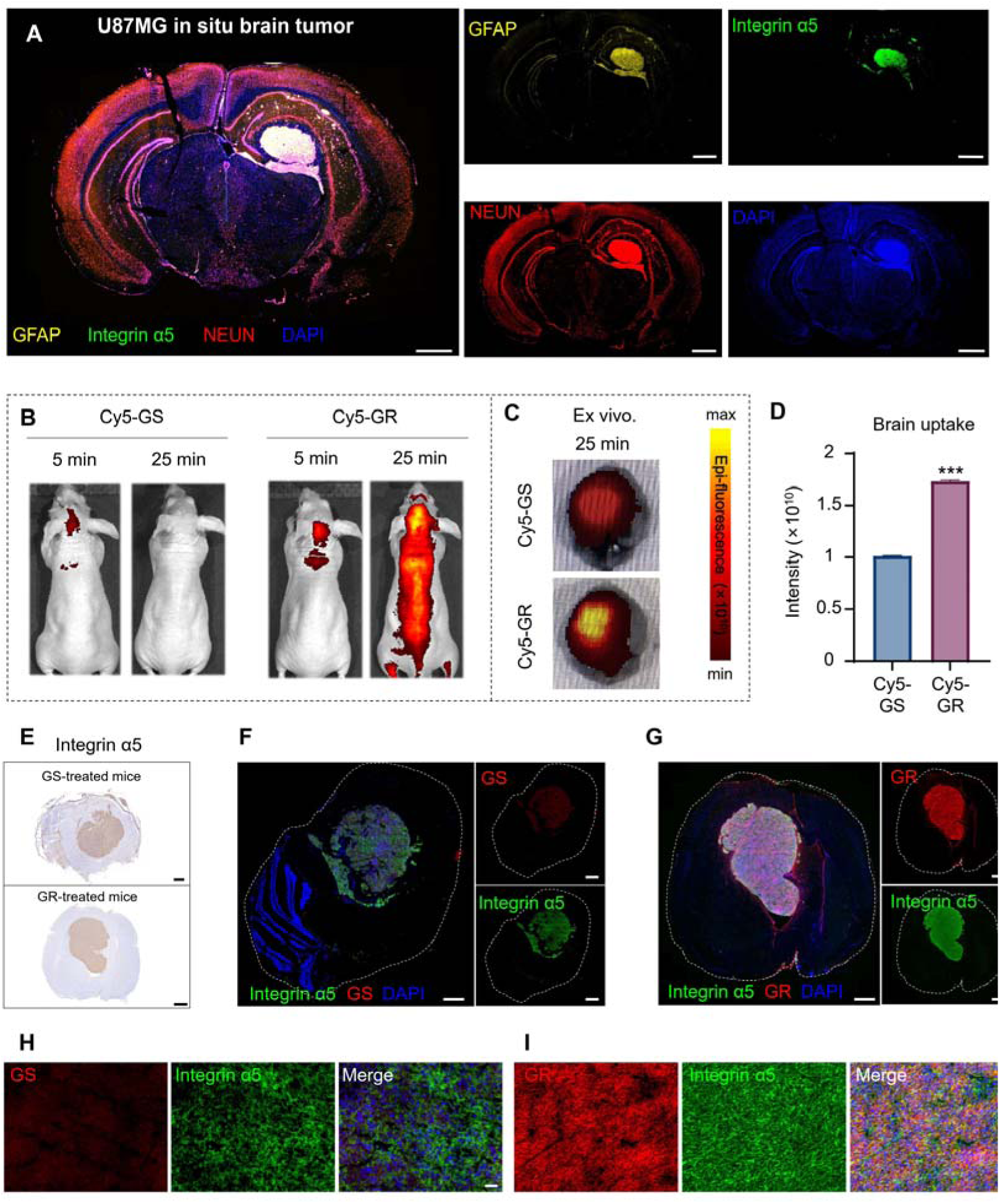
*In vivo* evaluation of BBB permeability of GS and GR in orthotopic glioblastoma tumor-bearing mice. **(A)** Immunofluorescence images show the expression of integrin α5β1, GFAP, and NEUN of U87MG tumors in orthotopic glioblastoma tumor-bearing mice. Green indicates integrin α5, Red for NEUN, yellow for GFAP, and blue for nucleus. Scale bar, 1 mm. **(B-C)** Representative fluorescence imaging of mice bearing in orthotopic U87MG tumors (**B**) and ex vivo brains (**C**). Cy5-GS and Cy5-GR were intravenously injected with a dose of 5 mg/kg. **(D)** Quantification of fluorescence intensity in tumors corresponding to (**C**). **(E)** Immunohistochemical analysis of integrin α5β1 expression in brain tissue sections from orthotopic glioblastoma tumor-bearing mice. Scale bar, 1 mm. **(F-G)** Immunofluorescence images of brain tissue sections from orthotopic glioblastoma tumor-bearing mice treated with Cy5-GS (**F**) and Cy5-GR (**G**). Green indicates integrin α5, Red for GS or GR, and blue for nucleus. Scale bar, 1 mm. **(H-I)** Magnific imaging of brain tissue sections from orthotopic glioblastoma tumor-bearing mice treated with GS-Cy5. Scale bar, 50 μm. All results are expressed as means ± SEM, as indicated in at least three independent experiments. A multiple t-test was used when two groups were compared. The symbol “*” represents differences compared with the Cy5-GS. *** p < 0.001.

Together these results demonstrated that GR is a potential candidate possessing not only ability to cross the BBB in both normal mice and orthotopic brain cancer mice, but strongly and stable binding to the glioblastoma specific biomarker integrin α5β1.

### ICG-GR supports maximal surgical resection in both subcutaneous and orthotopic glioblastoma mice

Based on its remarkable potential in glioblastoma targeting and BBB permeability, GR was utilized to develop the targeted glioblastoma intraoperative navigation modality (**Figure 6A**). GR was conjugated with near-infrared (NIR) fluorescent probe ICG, an FDA-approved probe for clinical cancer intraoperative navigation. We first investigated the *in vivo* performance of ICG-GR in healthy mice, which mainly accumulated in liver, kidney, spleen, and lung, consistent with the results observed in Cy5-GR. ICG-GR also showed evident uptake in the normal brains, demonstrating ICG conjugation did not impact its ability to cross BBB (**Figure 6B**). We administered ICG-GR intravenously to U87MG-bearing subcutaneous mice and performed multiple rounds of fluorescence imaging, showing evident tumor uptake (**Figure 6C**). The results indicated that the highest tumor-to-background fluorescence intensity ratio was achieved at 30 min post-injection, indicating the suitable time window for subsequent intraoperative fluorescent imaging surgery.

**Figure 6.**
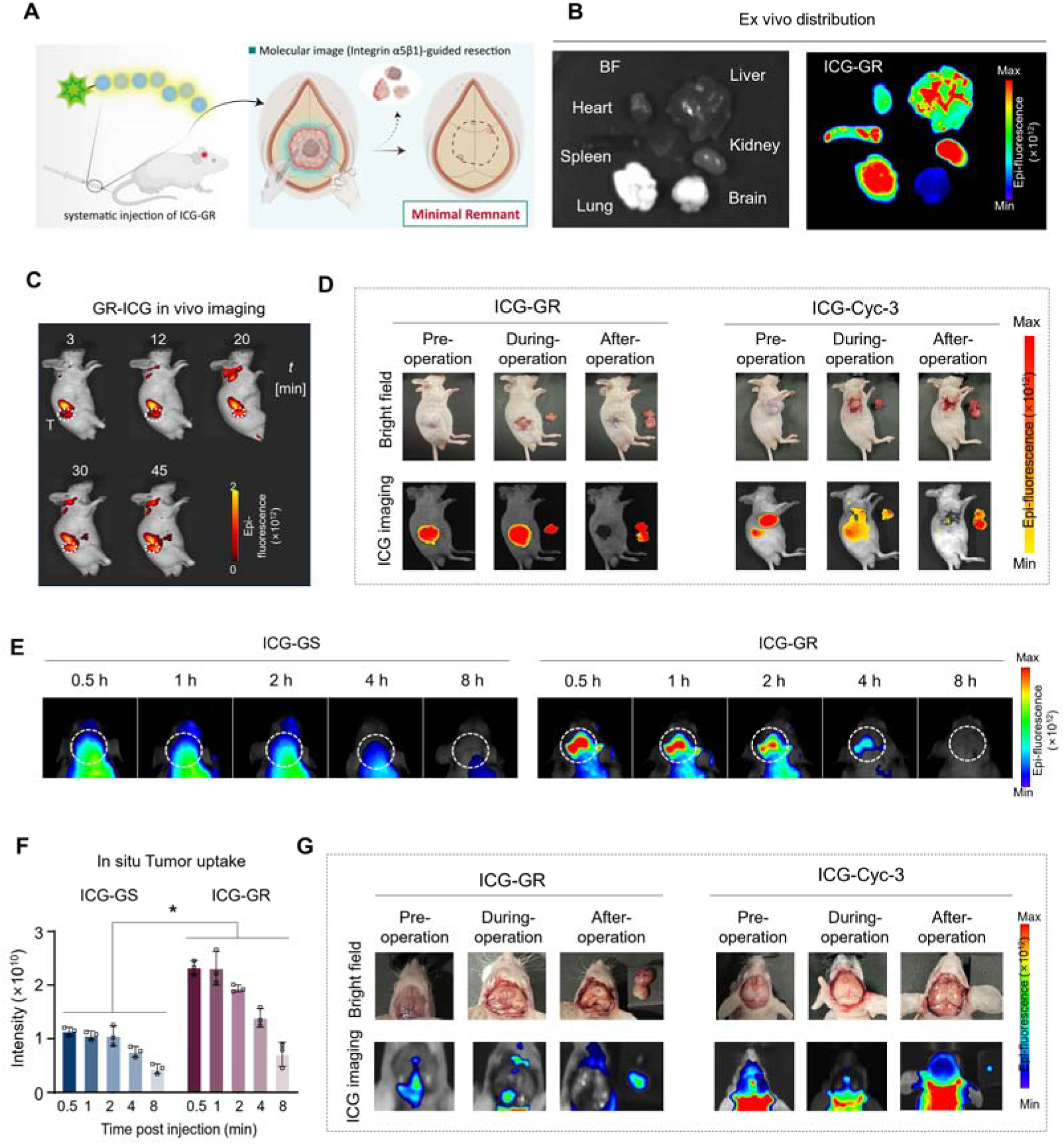
The intraoperative navigation of orthotopic U87MG brain tumor through ICG-labeled peptides. **(A)** Workflow for investigating the application of ICG-labeled peptides for intraoperative navigation. Peptides conjugated ICG were used to guide the resection of subcutaneous tumors. **(B)** Bright field and fluorescence images of the biodistribution of ICG-GR in healthy mice. **(C)** Representative fluorescence imaging of ICG-GR in subcutaneous U87MG tumor-bearing mice acquired at different time points after injection. **(D)** ICG-guided surgery in subcutaneous U87MG tumor-bearing mice at 30 min after injection of ICG-GR and ICG-Cyc-3. **(E-F)** Representative fluorescence imaging (**E**) and tumor qualification (**F**) of ICG-GR in orthotopic U87MG tumor-bearing mice acquired at different time points after injection. **(G)** ICG-guided surgery in orthotopic U87MG tumor-bearing mice at 30 min after injection of ICG-GR and ICG-Cyc-3. All results are expressed as means ± SEM, as indicated in at least three independent experiments. A multiple t-test was used when two groups were compared. The symbol “*” represents differences compared with the ICG-GS. * p < 0.05.

Then subcutaneous tumor was resected under the ICG-GR-guided intraoperative navigation. We performed *in vivo* NIR imaging pre-operation, intra-operation, and after-operation, which revealed that the tumor was completely removed in both subcutaneous (**Figure 6D**). Because of the enhanced ability to across the BBB, ICG-Cyc-3 was also synthesized for evaluating its potential in intraoperative navigation. The results demonstrate that ICG-Cyc-3 could also imaging the tumor border and achieve the maximum tumor resection (**Figure 6D**).

We further confirmed that ICG-GR exhibited efficient tumor uptake in orthotopic glioblastoma tumor-bearing mice, which is higher than ICG-GS (**Figure 6E-6F**). Similarity, the fluorescent signals and tumor-to-background ratio of ICG-GR reaches highest at 0.5 h post-injection, suggesting the suitable time window for orthotopic glioblastoma tumor-bearing mice. We further performed the ICG-GR-guided surgical resection in orthotopic glioblastoma tumor-bearing mice. The tumor was removed based on the ICG-GR fluorescence guiding, which enabled the completely tumor resection and no tumor uptake signal was observed after operation (**Figure 6G**). Consistently, ICG-Cyc-3 guided intraoperative navigation also allowed the maximum tumor removal (**Figure 6G**). Collectively, this study developed an integrin α5β1-targeted intraoperative navigation for maximal and safe removal of glioblastoma in the brains, and without causing adverse effect after surgery.

### ICG-GR guided resection followed [^177^Lu]GR intracavity TRT inhibits glioblastoma growth and recurrence

Radiopharmaceuticals labeled with therapeutic radionuclide result in direct and robust cell-killing. To achieve the maximal elimination of glioblastoma tumor tissues and reduce the possibility of tumor recurrence, we develop a strategy combined with integrin α5β1-targeted intraoperative navigation and intracavity TRT, which was achieved by the effective targeting peptide-GR. In this approach, ICG-GR was first intravenously injected, following by fluorescence imaging guided tumor resection 30 min post-injection. Immediately after surgery, [^177^Lu]GR was administrated in the tumor regions through intracavity injection, the tumor growth was observed within 24 days of treatment (**Figure 7A**). U87MG tumor-bearing mice were randomly assigned to four groups and treated according to different therapeutic regimens. Mice in “Group 1 (G1)” underwent tumor resection without ICG guidance and without [^177^Lu]GR administration. Mice enrolled in “Group 2 (G2)” underwent ICG-GR guided-tumor resection and without [^177^Lu]GR administration. Group 3 (G3) received an intracavity administration of [^177^Lu]GR after unguided tumor resection. Group 4 (G4) received an intracavity administration of [^177^Lu]GR following ICG-GR guided tumor resection (**Figure 7A**).

**Figure 7.**
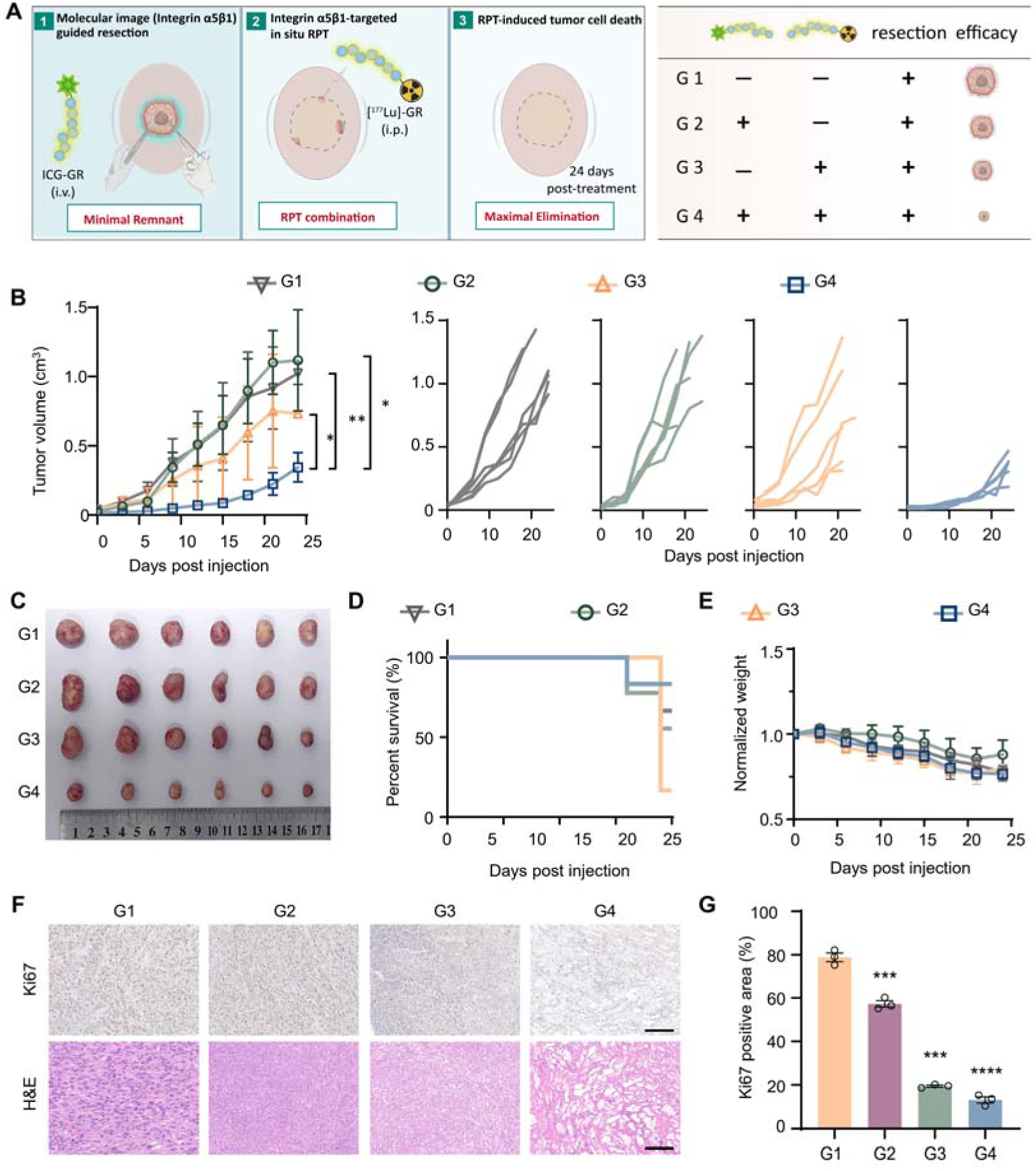
ICG-GR-guided surgical resection combined with intracavity [^177^Lu]GR-based TRT is a safe and effective treatment modality for glioblastoma. **(A)** Treatment scheme of ICG-GR-guided surgical resection combined with intracavity [^177^Lu]GR-based TRT. Mice are randomly divided into 4 groups, and each group is treated with a different protocol. The mice in G1 had their tumors removed in conventional way and the tumors of mice in G2 were removed under ICG guidance. Mice in G3 and G4 received conventional tumor resection ICG-guided tumor resection, respectively, followed by a single dose of subcutaneous injection after tumor resection. The tumor size and body weight were measured every three days. n = 6 mice per group. **(B)** Tumor volume curves for the groups or an individual mouse. ICG-GR-guided surgery combined with intracavity [^177^Lu]GR-based TRT showed the lowest tumor growth. **(C)** Photographs of tumors at the end of treatment 24 days post-injection. **(D)** Survival curves for 24 days after treatment. **(E)** Normalized body weights, which decreased slightly in all therapeutic groups. **(F)** H&E staining and Ki67 immunohistochemical images of tumors in four groups. Scale bar, 100 μm. g, Qualification of Ki67 positive signals according to immunohistochemical images. All results are expressed as means ± SEM, as indicated in at least three independent experiments. A multiple t-test was used when two groups were compared. The symbol “*” represents differences compared with mice in G1. *** p < 0.001. **** p < 0.0001.

Therapeutic efficacy was monitored for 24 days post-treatment (**Figure 7B-7C**). Mice in G1 and G2 exhibited rapid tumor growth, suggesting that tumor resection alone is insufficient to inhibit the cancer recurrence. Although ICG-GR guided tumor resection allows wider range of tumor resection, residual tumor cells is difficult to avoid. The high malignance and proliferation of residual tumor cells resulted in uncontrollable tumor growth. Therefore, ICG-GR guided tumor resection exhibited little advantages compared to unguided tumor resection when applied alone, which further demonstrated the necessity of TRT combination for maximal elimination of residual tumor cells. Tumor-bearing mice received intracavity administration of [^177^Lu]GR after tumor resection without ICG-GR guidance (G3) exhibited delayed tumor growth compared to G1 and G2, suggesting that [^177^Lu]GR could result in efficient tumor cell death. Importantly, the tumor volume in G4 was markedly reduced, compared to G1, G2, and G3. The significant reduced tumor growth of G4 when compared with G3 demonstrates that ICG-guided surgery achieving more tumor resection than blinded resection. Additionally, this finding further confirmed the critical role of postoperative [^177^Lu]GR intracavity administration in maximal elimination the residual tumor cells and inhibit tumor growth. The survival rate also indicated the therapeutic efficacy of the combined modality, with mice in G4 having the most survival individuals after 24 days of treatment (**Figure 7D**). No evident body weight loss was observed during the treatment and there is no significant difference between each group (**Figure 7E**). Additionally, less or no injury or abnormality was founded in the major organs at the end of treatment, suggesting that this is a safe modality and without causing unexpected adverse effect (**Figure S45**). The Ki67 expression was significantly decreased in tumors from mice in G3 and G4. H&E staining demonstrated the more necrotic area, which was consistent with the tumor growth data (**Figure 7F-7G**).

Together, integrin α5β1-targeted (GR) intraoperative navigation combined with postoperative intracavity TRT can significantly inhibit tumor growth and reduce the possibility of tumor recurrence, which is achieved by the minimal remnant of tumors and maximal elimination of residual tumor cells. Furthermore, this glioblastoma treatment strategy also serves as a safe modality because avoiding the systemic administration of RPT, which thoroughly prevents the major organ injury caused by blood circulation of radiopharmaceuticals. Therefore, this modality not only improved the therapeutic efficacy of RPT, but fundamentally avoid its *in vivo* toxicity, considered as a novel direction of next-generation RPT development, which is achieved by precising intraoperative navigation combination.

## Discussion

In this study, precise imaging of glioblastoma for maximal safe surgical resection, and robust elimination of residual tumor cells requires the development of tumor-targeted NIR fluorescence probes and radiopharmaceuticals. Favorable targeting binders with strong binding affinity and favorable BBB penetration enable efficient preoperative diagnosis, intraoperative navigation and intracavity TRT when labeled with ^68^Ga, ICG, or ^177^Lu. Because of its optimal binding pattern and specific recognition of integrin α5β1, GS are selected as the lead compound for targeting binder development. We first systematically modified GS through various strategies including cyclization, multivalency, covalent, and amino acid mutation. Among these derivates, the optimized candidate GR demonstrates superior binding affinity, cell permeability, and BBB penetration. Both ICR-GR and [^68^Ga]GR showed strong tumor accumulation in subcutaneous and orthotopic glioblastoma tissues. Notably, we harvested tumor tissues from glioblastoma patients with various integrin α5β1 expression and investigated the binding of GR in humans. GR exhibited robust binding to human glioblastoma, which showed strong correlation to integrin α5β1 levels. Therefore, we demonstrated the efficiency of GR-based PET/CT/NIR probes for precise and rapid diagnosis of human glioblastoma, endowing it values in clinical application.

Substitution of serine with arginine in linker sequence (GR) increases its positive charge, enhancing cell permeability, which is essential for BBB penetration and tumor accumulation. Moreover, the alteration in linker sequence did not disrupt the interaction of binding sequence to α5 and β1 subunits, which further mimic the distance between FN9 and FN10 domain of fibronectin ^36,37^. In contrast, cyclization, multivalency, and covalent modifications resulted in suboptimal *in vivo* performance, likely due to the unique binding pattern of GS to integrin α5β1, which restrict by the two binding sites of each subunit. PEGylated multivalency that introduced a large group result in steric hindrance and impairs integrin α5β1 binding. Similarity, the covalent linking to cysteine restricts the flexibility of targeting peptides, preventing proper subunit interaction. Cyclic peptides, especially Cyc-3 and Cyc-4, exhibit enhanced cell permeability and BBB penetration, suggesting the forming of helix structure. However, the cyclization of each recognition sequence of GS alters its secondary structure, leading to reduced tumor targeting. Therefore, this study reported the optimal strategy for simultaneously optimization both the BBB and tumor accumulation of GS, offering a potential platform for amphiphilic peptide modification in brain cancer.

The development of targeted fluorescence agents for precising tumor resection is the major focus in intraoperative navigation. A Trop2-targeted monoclonal antibody conjugated ICG agent enables rapid and accurate tumor margin assessment and improves relapse-free survival after intraoperative navigation resection^38^. Targeted fluorescent probes have also been explored for epithelial ovarian cancer, including CD24-tarageted AF750, folate receptor alpha-targeted FITC, and GnRHa-targeted ICG ^39^. Regarding imaging-guided glioblastoma resection, several studies have demonstrated improved tumor imaging capacity using targeted ligand conjugated nanoparticles or proteins to recognize tumor cells, nevertheless, their limited BBB penetration hampered the clinical utility^40–42^. Therefore, the glioblastoma specific biomarker should be applied for improve the target-to-background ratio and shorten the surgery window. To address this, we selected integrin α5β1 as a glioblastoma-specific biomarker. The integrin α5β1–targeted peptide GR conjugated with ICG enabled precise tumor imaging in both subcutaneous and orthotopic U87MG tumor-bearing mice. Importantly, ICG-GR–guided surgery achieved maximal resection with preservation of critical brain structures, a key requirement for improving outcomes in brain tumor treatment.

To improve therapeutic benefit, we combined ICG-GR–guided resection with postoperative intracavity [^177^Lu]GR administration. Our findings demonstrate that this approach achieved efficient eradication of residual tumor cells and significantly reduced recurrence. The combination of TRT and ICG-guided surgery offers several significant advantages over traditional treatment modalities. 1) Maximal tumor cell elimination. The application of ICG-guided resection enables accurate tumor intraoperative localization for maximal and safe resection. Coupled with the targeted intracavity delivery of [^177^Lu]GR, this approach significantly enhances the elimination of residual tumor cells, reducing the risk of recurrence. 2) Reduce systemic toxicity. In this combination therapy, radiopharmaceutical was applied through directly delivery to the tumor site intracavity, which minimizes systemic exposure of radiolabeled agents, thereby reducing the associated adverse effects and improving the safety of TRT. 3) Superior therapeutic option for glioblastoma. The major challenge facing glioblastoma is the high malignancy and difficulties in completely tumor removal, which result in high recurrence and poor prognosis. Multiple studies have reported the combination of intraoperative navigation tumor resection with immunotherapy, photodynamic therapy, and photothermal therapy^43,44^. However, the insufficient capacity in cell-killing may limited their clinical application. The remarkable advantages of RPT endow them a superior option for glioblastoma treatment. Currently, [¹□□Lu]DOTATATE, which has been approved by the U.S. FDA for TRT of neuroendocrine tumors, is now being developed for the treatment of glioblastoma and has entered a phase II clinical trial (NCT05109728). Additionally, a variety of other radiopharmaceuticals are also under development for the diagnosis and treatment of glioblastoma (NCT05739942, NCT05450744) ^45,46^, highlighting the promising potential of radiopharmaceuticals in the treatment of glioblastoma.

In conclusion, this study developed an integrin α5β1 selective peptide GR for precising and specific binding of glioblastoma. We demonstrated the efficacy of GR-based PET/CT/NIR probe for preoperative diagnosis and intraoperative imaging of glioblastoma. Notably, we reported a combined therapy involving ICG-GR-based fluorescence-guided surgical resection followed by intracavity [^177^Lu]GR-based TRT, which achieves maximal tumor resection and efficient elimination of residual tumor cells, thereby potential in reducing the recurrence and promoting the prognosis of patients suffered with glioblastoma. We report the feasibility of the novel combination therapy in treatment of brain tumors, which face the high-demanded needs for reduce the risk of recurrence. Furthermore, this approach also holds great potential for application in a variety of tumor types, marking the beginning of a new path in cancer treatment.

## RESOURCE AVAILABILITY

### Lead contact

Further information and requests for resources and reagents should be directed to and will be fulfilled by the lead contact, Kuan Hu.

### Materials availability

This study did not generate new unique reagents.

### Data and code availability

- All data needed to support the conclusions are presented in the paper or the supplemental information. Additional data related to this paper may be requested from the authors.
- This study does not report any custom code.
- Any additional information required to reanalyze the data reported in this paper is available from the lead contact upon request.

## ACKNOWLEDGMENTS

This study was supported by the National Natural Science Foundation of China (Nos. 82372002, 82502399, and 22507148), the Nonprofit Central Research Institute Fund of the Chinese Academy of Medical Sciences (No. 2022-RC350-04), the CAMS Innovation Fund for Medical Sciences (Nos. 2024-12M-ZH-009, 2021-I2M-3-001, 2023-I2M-2-006, 2021-I2M-1-026, 2025-I2M-XHJC-026, and 2025-I2M-XHXX-098), and the Beijing Nova Program and Beijing Nova Program Interdisciplinary Cooperation Project to K. H. This work was also supported by the Beijing Natural Science Foundation (Nos. L234044, L248087, L246051, and 7252206), Supported by the Fundamental Research Funds for the Central Universities, Peking Union Medical College (Nos. 3332025193, 3332025067, 3332025068, and 3332025153), the China Postdoctoral Science Foundation (No. 2025M773592, 2025M781897), the Postdoctoral Fellowship Program of CPSF (GZB20250842), the China National Nuclear Corporation Young Talent Program, and Medical + X Innovation Team of the Discipline Construction Enhancement Project, the Second Affiliated Hospital of Soochow University (XKTJ-TD202410), and the Beijing Municipal Health Commission Research Ward Excellence Clinical Research Program (BRWEP2024W032040202).

## AUTHOR CONTRIBUTIONS

K.H. and R.W. conceptualized the study. S.Q.Z., J.T.S. and Y.M.L. designed the methodology. J.T.S., X.K.S., D.F.W., and W.H.L. synthesized and characterized the integrin α5β1-targeting peptides. S.Q.Z., J.T.S., Y.M.L., J.W., C.L., Z.Z. and H.Y.H. performed radiolabeling, PET/CT imaging, intraoperative navigation, and therapeutic experiments. S.Q.Z., J.T.S., Y.M.L., and C.L., performed data analysis. S.Q.Z., J.T.S., and Y.M.L. wrote the original draft with contributions from L.Y.X, M.J.Y., and F.W., K.H., and R.W. review and editing the manuscript.

## METHOD DETAILS

### Reagents and equipment

All chemical reagents for peptides synthesis were purchased from GL Biochem (Shanghai, China), Macklin (Shanghai, China), Bide Pharmatech (Shanghai, China), and Beijing InnoChem Science & Technology (Beijing, China). All chemical reagents and solvents were used as received without further purification. Dulbecco’s modified Eagle’s medium (DEME medium, GIBCO, 11872093, NY, USA), penicillin-streptomycin (GIBCO, 15240062, NY, USA), and Dulbecco’s phosphate buffered saline (DPBS, GIBCO, C14190500BT, NY, USA) were purchased from Gibco and used as received. Fetal bovine serum (FBS, SE100-011, New Zealand) was purchased from VisTech.

All peptides were purified by a semi-preparative High Performance Liquid Chromatography (HPLC, SPD-16, Shimadzu, Japan) with the C18 reversed-phase column (4.6 × 250 mm, 5 μm, Waters, USA). The molecular weight of all peptides was confirmed by ultra-performance liquid chromatography-mass spectrometry (UPLC-MS, Waters 2998,Waters, USA). All radio-HPLC analyses for the radiotracers were performed using a Shimadzu HPLC system (LC-20AT, Shimadzu, Japan) equipment with FlowCount PRO for radioisotope detection (Eckert & Ziegler, Berlin, Germany). Gallium-68 was obtained at Nanjing First Hospital (Nanjing, China) by a 68Ge/68Ga generator (ITM, Munich, Germany). PET/CT scans were performed using an Inveon Micro PET/CT scanner (Siemens Medical Solutions, Knoxville, Munich, Germany). The radioactivity of biodistribution samples was measured using an automatic gamma counter (WIZARD2 2480, PerkinElmer, MA, USA). Biacore 8K (Cytiva, Uppsala, Sweden) was used for the SPR analysis. Confocal microscopy was conducted using an integrated fluorescence microscopy imaging system (BZ-X810, Keyence, Osaka, Japan). Flow cytometry was analyzed using Agilent NovoCyte Penton Flow Cytometer Systems (NovoCyte Penton Flow Cytometer Systems 5 lasers, Agilent, California, USA). The small animal imaging system was purchased from the Guangzhou Guangyi Biotechnology Co., Ltd. (Guangzhou, China).

### Animal ethics and human ethics

The experiment was approved by the Institutional Animal Care and Use Committee (IACUC) of the Institute of Materia Medica, Chinese Academy of Medical Sciences & Peking Union Medical College (IMM-S-25-0079). All animals received human care, and the experiments were conducted according to the recommendations of the Committee for the Care and Use of Laboratory Animals.

Glioblastoma tumor tissues used to analyze the expression of integrin α5β1 and evaluation the tumor accumulation of integrin α5β1-targeted peptides was obtained from Beijing Tiantan Hospital. The Ethics Committee of Beijing Tiantan Hospital approved this study, and informed consent was obtained from all participants (KY-2024-340-03).

### Cell lines and animal models

Human glioblastoma cell line U87MG and human glioblastoma cell line transferred with luciferase gene luc2 (U87MG-Luc2) were obtained from Meisen Life Science & Technology Co. (Zhejiang, China). All the cell lines were cultured in DMEM containing high glucose (GIBCO, Carlsbad, CA, USA), which was supplemented with 10% FBS and 1% penicillin-streptomycin. The cells were cultured in a humidified atmosphere of 5% CO_2_ at 37 □. The medium was changed every day and passages were performed every 2 days.

The Balb/c Nude mice (5 weeks old) and ICR mouse were purchased from GemPharmatech (Beijing, China). For subcutaneous tumor-bearing mice, approximately 1×10^6^ U87MG cells were harvested and suspended in serum-free DMEM at a total volume of 0.1 mL and subcutaneously injected into the right flank. Two weeks after inoculation when tumor volumes reached 300-400 mm^3^, tumor-bearing mice were used for PET/CT imaging, fluorescence imaging, in vivo biodistribution study and ICG-guided tumor resection. Mice were euthanized once tumors reached a volume of 1.5 cm^3^. For orthotopic glioblastoma-bearing mice, a 2□mm diameter pore was generated on the left parietal bone using cranial drill. Then, 1×10^5^ U87MG-Luc cells were intracavity injected into the cerebral cortex approximately 250□μm below the brain surface using a stereotaxic apparatus. The growth of orthotopic glioblastoma was observed by in vivo fluorescence imaging of Luc signals.

### Solid-phase peptide synthesis

Series of integrin α5β1-targeted peptides were manually synthesized using the Fmoc solid-phase peptide synthesis strategy as previously described ^47,48^. Briefly, Fmoc-protected amino acids in sequences were coupled in turn under the presence of HCTU (2-(6-Chloro-1H-benzo[d][1,2,3]triazol-1-yl)-1,1,3,3-tetramethyluronium hexafluorophosphate(V), O804759-500g, Macklin) and DIPEA (N,N-Diisopropylethylamine, N807281-500ml, Macklin) (amino acids/HCTU/DIPEA, n/n/n, 4/4/8) in DMF (n,n-dimethyl-formicaci, N807508-4L, Macklin) solvent for 1 h. After each coupling step, the Fmoc was removed through reaction with 50% morpholine/DMF for 30 min. Following peptide synthesis, a beta-alanine was attached to the N-terminal of peptides to link the bifunctional chelator DOTA (1,4,7,10-tetraazacyclododecane-N,N’,N,N’-tetraacetic acid, BD126860, Bidepharm), ICG, or Cy5 to peptides. Next, the peptide precursors were deprotected and removed from the resin using a mixture of TFA (Trifluoroacetic acid, T818782-500ml, Macklin), TIPS (Triisopropylsilane, T819181-100ml, Macklin), and water (TFA/TIPS/water, v/v/v, 95/2.5/2.5) for 2 h. Once excised, the peptide precursors were purified using the semi-preparative HPLC, and HPLC conditions were as follows: flow rate = 1 mL/min; λ = 220 nm; A = 0.1% TFA/H_2_O; B = 0.1% TFA/acetonitrile; B gradient: 0-20 min, from 0 to 94%. UPLC-MS were used to confirm the successful synthesis of each peptide. Then the purified peptides were lyophilized and stored at −20 □.

### Surface plasmon resonance (SPR) for evaluation the binding affinity of integrin α5β1-targeted peptides

The binding affinity of each peptide was determined by SPR assay. Recombinant human integrin α5β1 (alpha 5 beta 1, HY-P77718, MCE, NJ, USA) was immobilized on a CM5 sensor chip at 25 □ following a standard amine coupling kit. The final immobilization level of integrin α5β1 was 5500-7500 rack unit (RU). Peptides were injected as analytes at various concentrations in HBS-P (BR100368, Cytiva, MA, USA) running buffer (10 mmol/L HEPES buffer, 150 mmol/L NaCl, 0.02% surfactant P20, pH 7.4) containing 5% DMSO (Dimethyl sulfoxide, A68908-500ml, Innochem) to evaluate the binding affinity. The chip platform contact time was 120 s, and the dissociation time was 240 s. The study was conducted by Biacore 8K system (Cytiva, MA, USA). The data was analyzed using Biacore evaluation software (8K version 1.0), fitted using a 1:1 steady affinity model and plotted using GraphPad Prism 8.0.2.

### Radiochemical synthesis and quality control of integrin α5β1-targeted peptides

For the ^68^Ga and ^177^Lu labeling, 20 μg of each peptide were dissolved in double distilled water (20 μL) and then added to a solution of ^68^GaCl_3_ or ^177^LuCl_3_ (185 MBq) in sodium acetate (0.1 M, pH = 4∼4.5, 100 μL), and reacted at 100 □ for 15 min. For the ^64^Cu labeling, 20 μg of each peptide (GS or GR) and ^64^CuCl_2_ (350-400 MBq, 0.1 mol/L NaOAc, pH = 4.1) were mixed and the reacted for 10 min at 80°C.

Radiochemical yield (RCY) of the labeled radiotracers was analyzed by radio-HPLC using YMC-Triat-C18 column (4.6X150 mm, 5 μm) under the following conditions: flow rate = 1 mL/min; solvent A = 0.1% TFA/H_2_O; solvent B = 0.1% TFA/acetonitrile. B gradient: 0-20 min, from 10% to 90%. The radioactivity was measured with a CRC-25R Dose Calibrator (CAPIN-TEC, USA). For measurement of LogP, 1 mL of saline,1 mL of n-octanol, 0.37MBq radiotracers were added in a 5 mL centrifuge tube, and the PBS and n-octanol were completely layered after standing at room temperature for about 2 h. The 10 μL of upper (n-octanol) and lower (saline) solutions were transferred to a radioimmunotherapy tube, respectively. The radioactivity was measured with an automatic gamma counter (WIZARD^2^ 2480, PerkinElmer Instruments Inc., USA). The lipid-water distribution coefficient (LogP_O/W_) is calculated by the following formula: LogP_O/W_ = Log(CPM _in-octanol_ /CPM _in saline_). Three parallel samples were set up for each radiotracer, and the result was expressed as mean ± SD (n = 3).

### *In vitro* cellular uptake assays

For the cell binding assay, U87MG cells were treated with different concentrations (500, 600, 800, 900, 1000 nM) of [^68^Ga]GS and [^68^Ga]GR. The cells were incubated at 37 □ for 1 h. After incubation, the medium was removed, and the cells were washed with cold PBS three times. Then, 300 μL of 0.2 M NaOH was added, and the cell lysate was collected into 1.5 mL Eppendorf tubes. Radioactivity in each tube was measured with an autogamma counter (PerkinElmer WIZARD2 2480, MA, USA).

### Secondary structure prediction and molecular docking

The secondary structures of the peptides were predicted by Alphafold3 and visualized by PyMOL 2.5. For molecular docking, the 3D structures were initially set up with ChemDraw 3D 21.0.0 using the MM2 force field for energy minimization and then exported in the mol2 format. Subsequently, the structures were processed using AutoDock 1.5.7 and exported in the pdbqt format. The 3D crystal structure of the integrin α5β1 (PDB: 7NWL) was obtained from the PDB database. According to the literature, the active binding site of integrin α5β1 is located at coordinates x = 273.238, y = 282.787, and z = 258.107.^49–51^ This active site was used as the docking target. Molecular docking was performed using AutoDock 1.5.7, and the results were visualized using PyMOL 2.5.

### Immunohistochemistry staining

For immunohistochemistry staining of subcutaneous and orthotopic U87MG tumors, the activity of endoperoxides was inactivated using an H_2_O_2_ solution (3%) after penetration in Triton-X100. Following treatment with 3% BSA for 2 h at room temperature, the sections were incubated with anti-integrin α5 and Ki67 primary monoclonal antibody overnight at 4 □. After being washed with PBS for three times, HRP-labeled secondary antibody (Servicebio, GB23303) was added to the samples for 1 h at room temperature and 3,3-diaminobenzidine (DAB). Horseradish Peroxidase Color Kit (Beyotime, P0202, Shanghai, China) was then used to visualize the expression of integrin a5. Finally, hematoxylin staining was conducted following standard procedures. An optical microscope (OLYMPUS, Japan) was used to observe the stained sections and generate high-quality pictures.

### Western Blot

The U87MG cells, subcutaneous U87MG and patient glioblastoma biopsies were treated with RIPA (Radioimmunoprecipitation) lysis buffer (Beyotime, P0013B, Shanghai, China) for cell lysate collection. After centrifuging at 12,000 rpm for 20 min, the supernatant was collected. The total protein concentration was determined using a BCA Protein Assay Kit (Bicinchoninic Acid Protein Assay Kit, Beyotime, P0009, Shanghai, China). For Western blotting, the cell lysate was electrophoresed on 8% SDS□PAGE by 80 V for 20 min, followed by 200 V for 30 min. Then, the protein was transferred to a polyvinylidene difluoride (PVDF) membrane for 2 h at 4 □. After transferring, 5% skim milk was added to the PVDF membrane for blocking over 3 h in a shaker at room temperature. Primary antibodies against integrin α5 (ab150361, 1:5000; Abcam, Cambridge, UK) added and incubated overnight at 4 □ with agitation. After washing with 1% TBST three times, the PVDF membrane was incubated with secondary antibody for 1 h at room temperature. Finally, the immunoblots were visualized with enhanced chemiluminescence (ECL) solution by imaging system (BG-gdsAUTO 720, boygene, Shanghai, China).

### Flow cytometry

The U87MG cells with a density of 2 × 10^5^ cells per well in 24-well plate were incubated with Cy5-labeled GS, GD, GE, R, GR, Cyc-1, Cyc-2, Cyc-3, Cyc-4, Salk, Scl, Ssf, Ralk, Rcl, and Rsf (20 μM) for 2, 4, 8, and 24 h at 37 □. After been washed with PBS for three times, the cells were collected and analysed using Invitrogen™ Attune™ NxT Flow Cytometer (ThermoFisher, MA, USA). FlowJo software was used to measure the number of Cy5-positive cells and their mean fluorescence intensity.

### Confocal imaging for evaluation the cell permeability of each peptide

Cy5-labeled peptides were initially dissolved in HFIP at a concentration of 1 mg/ml. The solvent was then evaporated under a gentle stream of nitrogen gas. The resulting peptide film was reconstituted in water to obtain a 100 μM stock solution, which was further diluted 10-fold in culture medium to achieve a final concentration of 10 μM. U87MG cells were incubated with Cy5-conjugated peptides. After 4, 12, 18, and 24 hours of incubation, the cells were washed with PBS, followed by staining with DAPI for 10 min. Subsequent PBS washing was performed. Cellular uptake was analyzed using the integrated fluorescence microscopy imaging system with DAPI/Cy5 channels.

### PET/CT imaging

PET/CT scans were performed using an Inveon Micro PET/CT scanner (Siemens Medical Solutions, Knoxville, Munich, Germany). The 3.7 MBq of radiotracers dissolved in 100 μL saline (0.9%) were intravenously injected into each mouse (normal mouse and U87MG tumor-bearing mouse, n = 3) and anesthetized with 1.5% isoflurane. The dynamic PET/CT scans were started immediately post-injection and lasted for 1 h. The static PET/CT images were acquired at 0.5 h, 1 h, 2 h, and 4 h post-injection. Images were reconstructed by a three-dimensional ordered subsets expectation maximum (3D OSEM) algorithm and converted to %ID/g images. The CT data from the PET/CT examination were reconstructed in the transverse plane as 0.1 mm thick sections. The following parameters were used for imaging: 80 kV, 100 mA, and 0.32 s per rotation. The radioactivity was decay-corrected for the injection time and expressed as the percent of the total injection dose per gram (%ID/g) at the Inveon Research Workstation. To compare the radioactive signals in the tumors and other tissues, regions of interest (ROIs) were drawn.

### *Ex vivo* biodistribution in normal and U87MG-bearing mice

Ex vivo biodistribution of [^68^Ga]GS and [^68^Ga]GR were evaluated in normal mice and subcutaneous U87MG-bearing mice (n = 3). Each mouse was injected intravenously with about 1.85 MBq of each radiotracer in 0.1 mL of saline and sacrificed at 5 min, 30 min, 1 h, 2 h, and 4 h post-injection. The main organs, including blood, heart, liver, spleen, lung, kidney, intestine, pancreas, muscle, and tumor (for U87MG-bearing mice group only), were collected and weighed. The radioactivity of all organs was measured using an automatic gamma counter (WIZARD2 2480, PerkinElmer, MA, USA) and expressed as a percentage of the %ID/g.

### Immunofluorescence staining

For immunofluorescence staining, optimal cutting temperature compound (OCT)-embedded tumor tissues from orthotopic tumor-bearing mice or glioblastoma patients were sliced into 15-μm-thick sections and stored at −80 □ for future use. The tumor sections or U87MG cells were treated with 0.1% Triton X-100 and then blocked in 3% bovine serum albumin (BSA) at room temperature for 2 h. Then, the sections were incubated with primary monoclonal antibody overnight at 4 □. The primary antibody used was listed as below: GFAP (16825-1-AP, 1:200; Proteintech, Wuhan, China), NEUN (66836-1-lg, 1:200; Proteintech, Wuhan, China), anti-integrin α5 (ab150361, 1:400; Abcam, Cambridge, UK), anti-integrin β1 (sc-9970, 1:500; Santa Cruze Biotechnology, Texas, USA). After being washed with PBS for three times, secondary antibody was added to the samples for 1 h at room temperature, which was listed as blow: goat anti-rabbit IgG H&L alexa fluor@568 (ab175471, 1:1000; Abcam, Cambridge, UK), and goat anti-mouse IgG H&L alexa fluor@488 (ab150113, 1:500; Abcam, Cambridge, UK), goat anti-rabbit IgG H&L alexa fluor@647, ab150083, 1:1000; Abcam, Cambridge, UK). For the analysis of peptide binding in patient glioblastoma tissues, ICG-GS or ICG-GR (10 μM) was then incubated with sections for 1 h at room temperature in dark. DAPI was used to label the cell nucleus, and the stained tissues were observed under a confocal fluorescence microscope. For the analysis of peptide binding in orthotopic U87MG tumors, Cy5-GS and Cy5-GR were i.v. injected. Mouse were sacrificed and subjected to dissection to harvest brain 30 min post-injection. Immunofluorescence staining of integrin α5 was performed as described above.

### Autoradiography

Frozen sections of six glioblastoma patient tumors were incubated with [^68^Ga]GR (2 μCi/ml each). After 2 hours incubation, samples were then exposed to phosphor imaging plates for 1 h, and ARG signals were subsequently acquired using a phosphor screen scanner.

### BBB organoids culture

Sterile 1% agarose (w/v) was prepared by adding 0.1 mg of molecular biology grade agarose into 10 mL of DPBS in a flask, and boiled with microwave until completely dissolved. The melted agarose solution was then dispensed into each well of a 96-well plate (50 μL per well) while it was still hot using a multi-channel pipette, and allowed to solidify (∼15 min). Approximately cerebral microvascular endothelial cells (hCMEC/D3 cell line), HBVPs, and U87MG resuspended in EBM-2 working medium (10^3^ cells for each cell type, supplemented with 2% human serum) were seeded onto the agarose gel in each well of the 96-well plate in a 1:1 ratio (final volume = 200 μL). Cells were placed in a humidified incubator at 37 □ with 5% CO_2_ for 48 h to allow for the assembly of multicellular BBB organoids.

### *In vitro* autoradiography of BBB organoids

BBB organoids were established for 48 h as described before. Then, 100 μL of EBM-2 medium containing 50 μCi/mL peptides labelled with ^68^Ga ([^68^Ga]GS, [^68^Ga]GR, [^68^Ga]Cyc-3, [^68^Ga]Cyc-4, [^68^Ga]Scl, and [^68^Ga]Rcl, n=3) was added to each well. It was placed in a humidified incubator at 37 □ with 5% CO_2_ for 2 hours. After incubation, each well was washed with DPBS for three times, dried in air, and then exposed to imaging plates for 120 min with accuracy of 25 um and voltage of 1000 mV. The radioactivity in each well was quantified using the ImageJ software and obtained in terms of photo stimulated luminescence per unit area (photostimulated luminescence/mm^2^).

### Transwell co-culture model

Both the apical and basal sides of 24-well transwell inserts were coated with rat-tail collagen (5 μg/mL in DPBS) overnight at 4 □. Each transwell insert was inverted, and 5 × 10^5^ HBVP cells were seeded on the basal side of the insert. The inserts were encased in an inverted plate and allowed to adhere for 2 h at 37 □ with 5% CO_2_. Inserts were then removed and placed within a 24-well plate containing 300 μL EGM-2 medium, ensuring that the HBVP cells on the basal side of the insert was properly submerged in the medium. 5.0 × 10^7^ hCMEC/D3 cells were resuspended in 200 μL EGM-2 medium and seeded on the apical side of the insert. In a separate 24-well plate, 5.0 × 10^4^ U87MG were seeded in DMEM (supplemented with 10% FBS and 1% P/S). All cells were allowed to adhere overnight at 37 □ with 5% CO_2_. Next day, U87MG cells in the 24-well plate, cells on both apical and basal sides of the transwells were washed twice with EGM-2 medium. The inserts (containing hCMEC/d3 on the apical side, and HBVP on the basal side) were then transferred into the 24-well plate containing the U87MG cells using a sterile forcep. The final volume of the upper EGM-2 medium was 500 μL and 200 μL on the lower part of the insert. The transwell co-culture model was placed in a humidified incubator at 37□ with 5% CO_2_. The medium was replaced with fresh EGM-2 medium every 2 days.

### Evaluation of peptides crossing the BBB in transwell model

The hCMEC/D3 medium in the basal compartment was replaced with 200 μL EGM-2 medium containing 20 μCi peptides labelled with ^68^Ga([^68^Ga]-GR/GS/Cyc-3/Cyc-4/Rcl/Scl) for 4 h. The medium in the basal compartment and the apical side was collected. U87MG cells seeded in the 24-well plate were washed for three times with DPBS, released by 0.25% trypsin and then collected as well. The radioactivity was measured using an automatic gamma counter (WIZARD2 2480, PerkinElmer, MA, USA) and expressed as a percentage of the dosage per well.

### *In vivo* fluorescence imaging

Cy5-conjugated peptides (GS, GD, GE, GR, R, Cyc-1, Cyc-2, Cyc-3, Cyc-4, Salk, Scl, Ssf, Ralk, Rcl and Rsf) were administered to healthy mice. After 1 h, mice were sacrificed and subjected to dissection to harvest organs (heart, liver, spleen, lung, kidney, brain), small animal imaging system performed imaging.

U87MG-bearing subcutaneous and orthotopic mice with a tumor volume of about 300 mm^3^ were intravenously injected with Cy5-conjugated peptides with a dose of 5 mg/kg. A small animal *in vivo* imaging system (LASER6000, Bio-io, Guangzhou, China) was used to perform imaging post-injection. To compare the fluorescence signals in the tumors and other tissues, regions of interest (ROIs) were drawn.

ICG-GS and ICG-GR were intravenously injected with a dose of 5 mg/kg in both U87MG-bearing subcutaneous and orthotopic mice. The excitation light was provided by an 808-nm diode laser. NIR-II fluorescence images were collected also using small animal *in vivo* imaging system (LASER6000, Bio-io, Guangzhou, China). ICG-GR was administered to healthy mice. Mice were sacrificed and subjected to dissection to harvest organs (heart, liver, spleen, lung, kidney, and brain). Small animal imaging was performed to observe the accumulation of each peptide to major organs.

### ICG fluorescence imaging -guided tumor resection in subcutaneous and orthotopic tumor-bearing mice

We conducted ICG-guided tumor resection experiments in both U87MG-bearing subcutaneous and orthotopic mice. The mice were performed imaging-guided surgery under 1.5 % isoflurane anesthesia. ICG labeled peptides (5 mg/kg) were injected into the mice via tail vein, and imaging-guided tumor resection was performed 30 min post-injection. Tumors were visualized with NIR-II fluorescence and the tumor with positive signals were removed until there are no fluorescence signals in local lesions. The surgical operations were similar for subcutaneous and orthotopic tumor-bearing mice.

### ICG-GR-guided surgical resection combined with postoperative intracavity TRT

U87MG-bearing mice with a tumor volume of 300 mm^3^ were randomly divided into four groups. The mice in Group 1 (G1) had their tumors removed without ICG-GR-guided and the tumors of mice in Group 2 (G2) were removed under ICG-GR imaging-guidance. Mice in Group 3 (G3) received a single dose of subcutaneous injection of [^177^Lu]GR after tumor resection under visible white light. Mice in Group 4 (G4) received ICG-GR imaging-guided tumor resection, followed by an in situ injection of [^177^Lu]GR. The tumor size and body weight were measured every three days after surgery. The treatment experiment lasted for 24 days. At the end of the experiment, we collected tumor tissues to record the tumor growth of each group, and took tissues for H&E and IHC staining.

### H&E staining

Preparation of paraffin sections from patient tissues and U87MG tumor-bearing mice. Paraffin-embedded tissue sections were dewaxed in xylene and rehydrated through a graded ethanol series. The sections were stained with hematoxylin for 5-10 min, followed by a brief water rinse. After bluing with running water, the sections were rinsed again with water. The sections were then dehydrated in 95% ethanol for 1 min and stained in eosin solution for 10-30 s. Then, the sections were dehydrated through a graded ethanol series and cleared in xylene. Finally, the sections were mounted with neutral balsam, and the tissue structure was observed under a light microscope.

### Statistical analysis

Statistical analyses were performed with GraphPad Prism 9.0. All quantitative data were expressed as mean ± SD, as indicated in at least three independent experiments. An unpaired two-tailed Student’s t-tests were used to measure the significant differences. Statistical significance was defined at *p ≤ 0.05, **p ≤ 0.01, ***p ≤ 0.001, and ****p ≤ 0.0001.

