## Supplemental Figure 1 - 45 and Supplemental Table 1 for "Development of Integrin α5β1-targeted PET/NIR imaging probes for glioblastoma intraoperative navigation and intracavity targeted radionuclide therapy"

**Affiliations:**

DOTA-GS

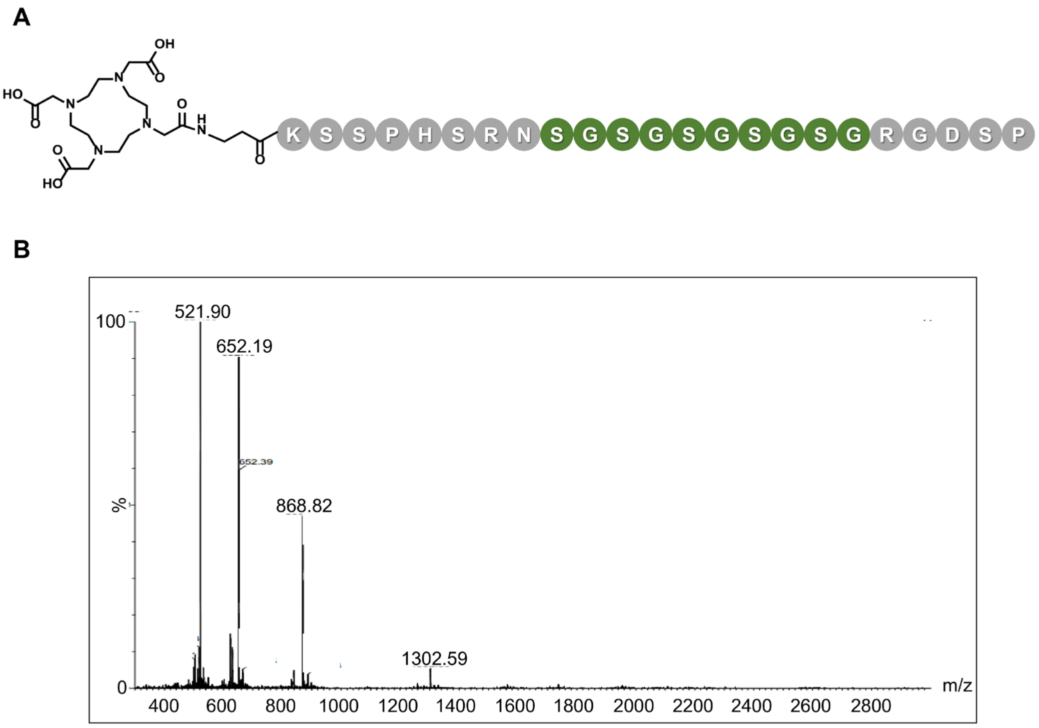


**Figure S1.** MS analysis of the DOTA-GS. The purity of DOTA-GS was greater than 90%.

DOTA-GD

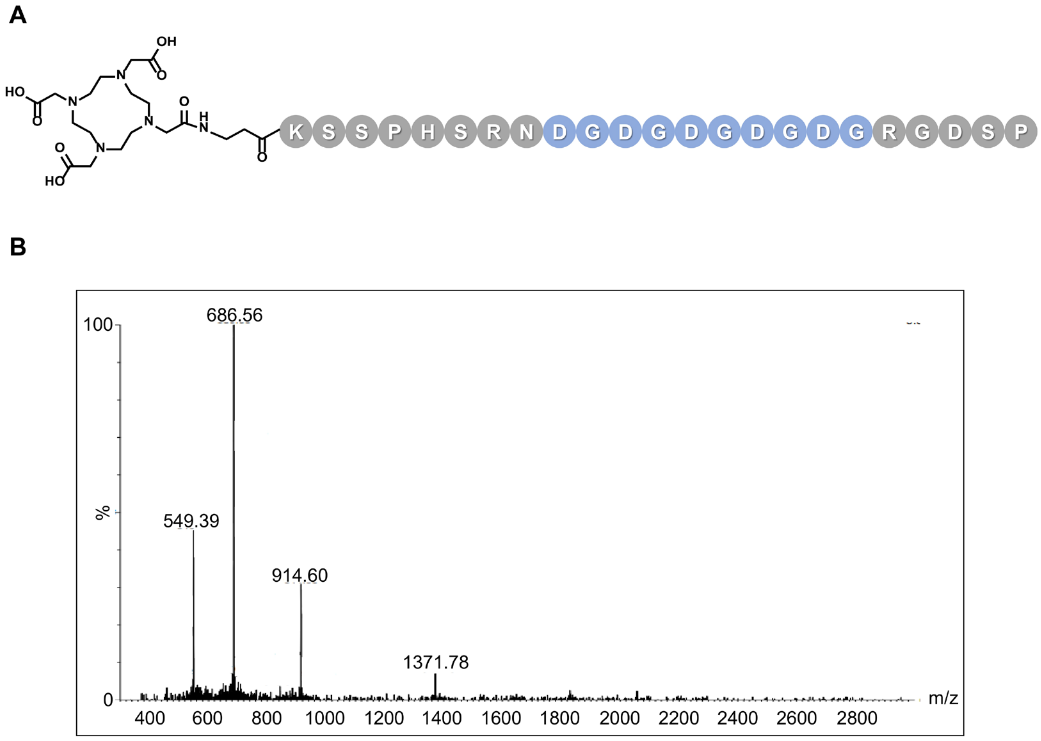


**Figure S2.** MS analysis of the DOTA-GD. The purity of DOTA-GD was greater than 90%.

DOTA-GE

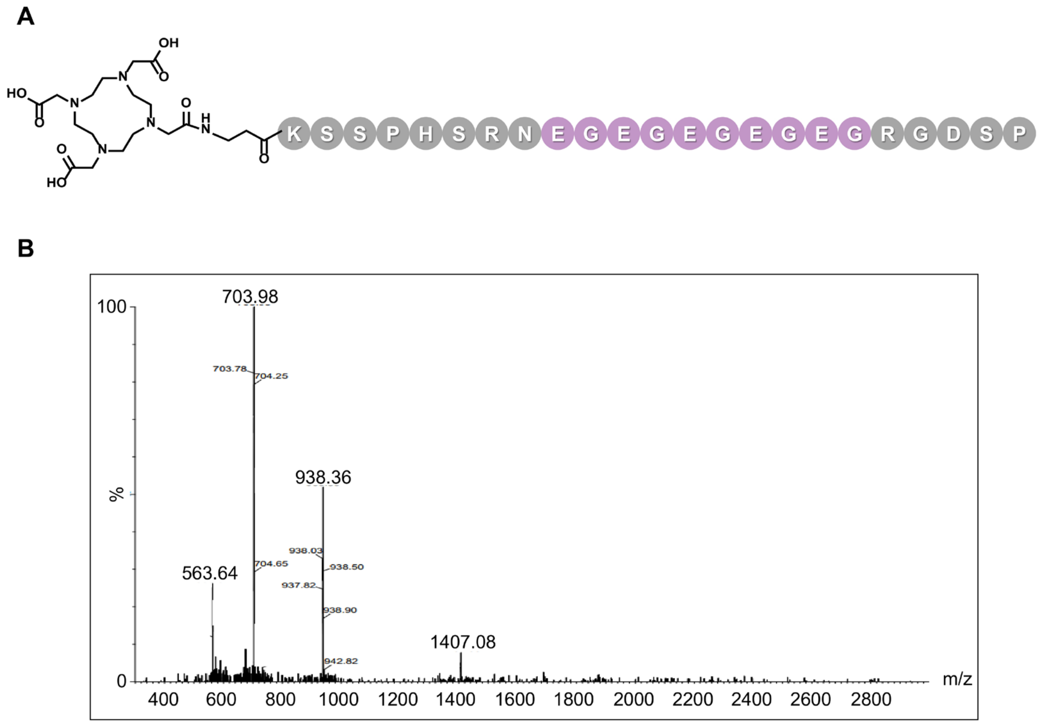


**Figure S3.** MS analysis of the DOTA-GE. The purity of DOTA-GE was greater than 90%.

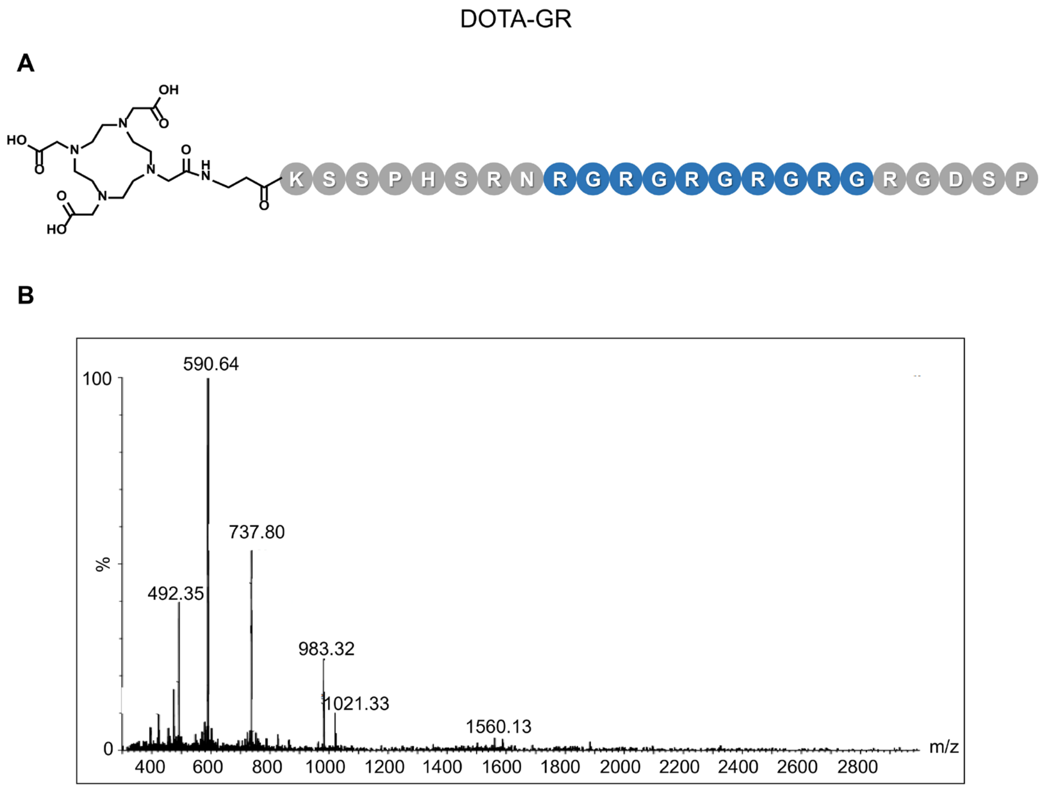


**Figure S4.** MS analysis of the DOTA-GR. The purity of DOTA-GR was greater than 90%.

DOTA-R

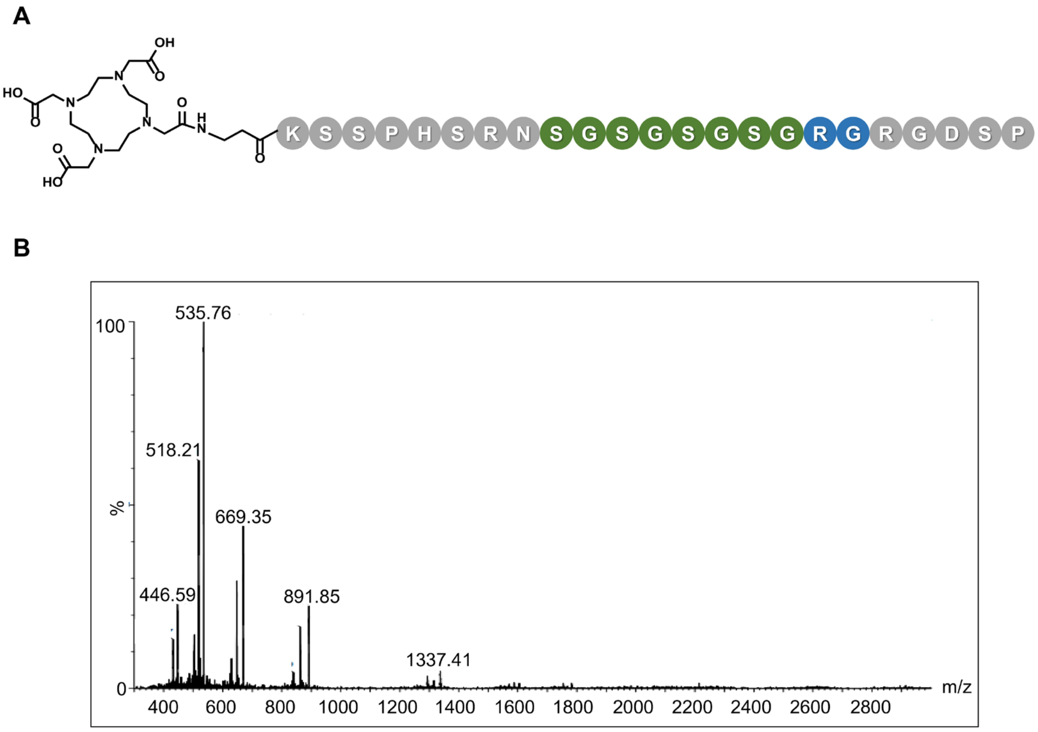


**Figure S5.** MS analysis of the DOTA-R. The purity of DOTA-R was greater than 90%.

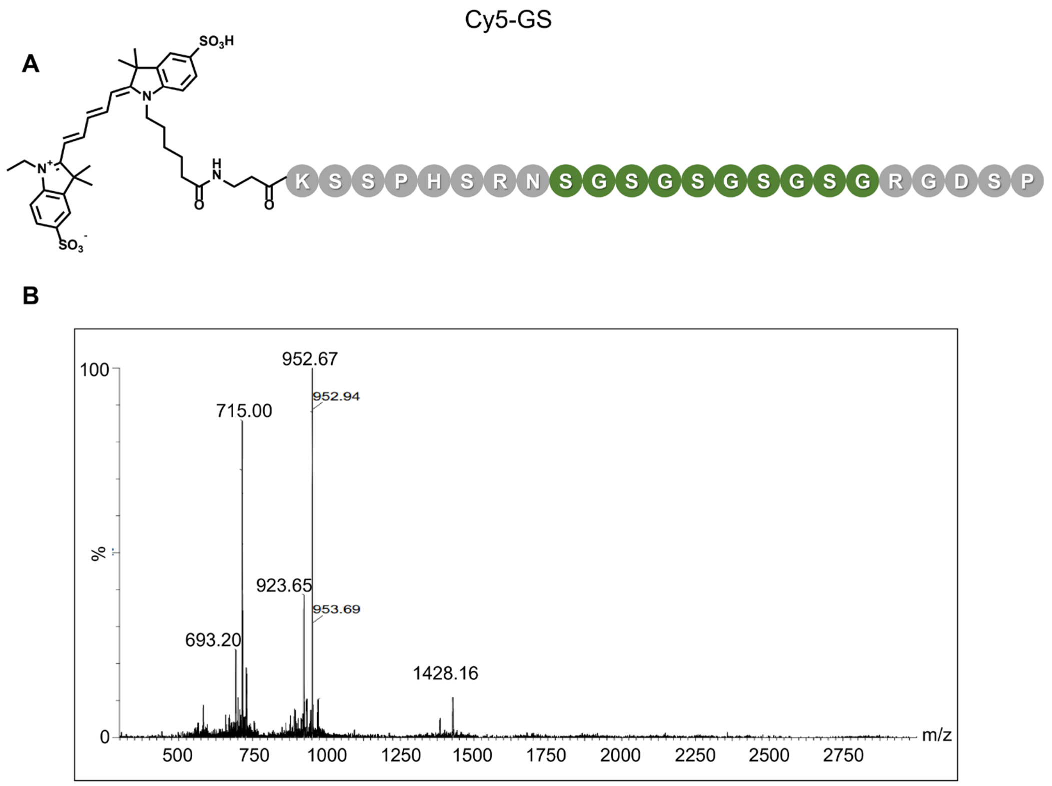


**Figure S6.** MS analysis of the Cy5-GS. The purity of Cy5-GS was greater than 90%.

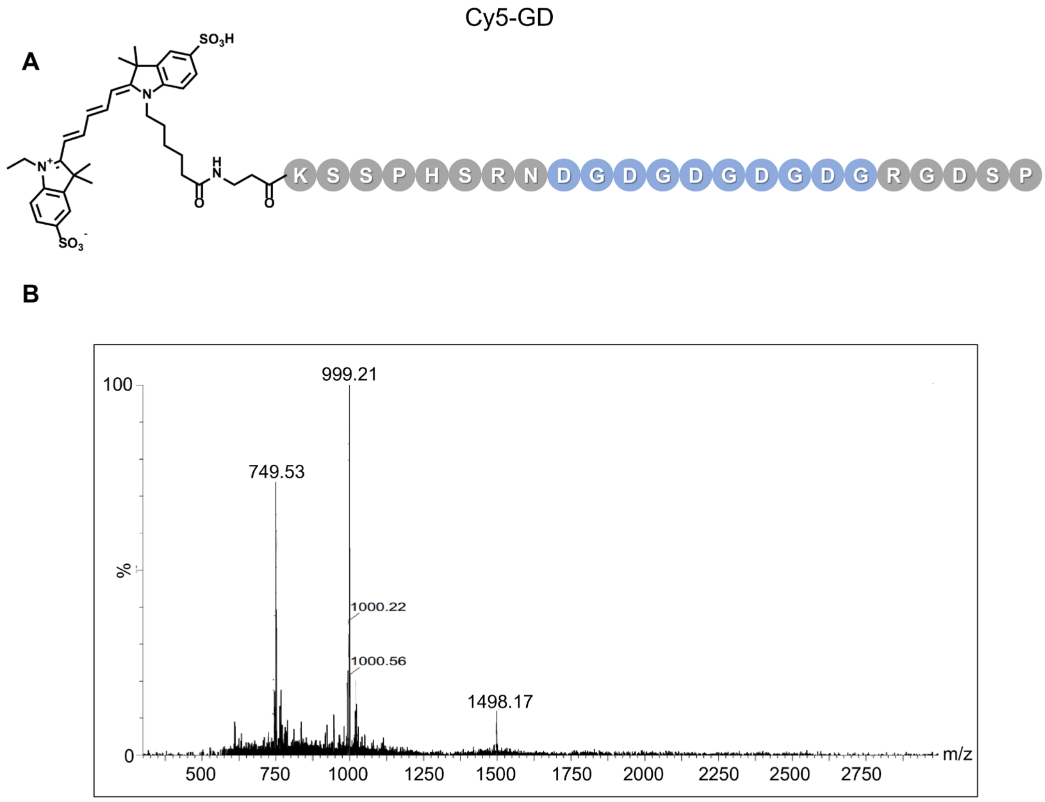


**Figure S7.** MS analysis of the Cy5-GD. The purity of Cy5-GD was greater than 90%.

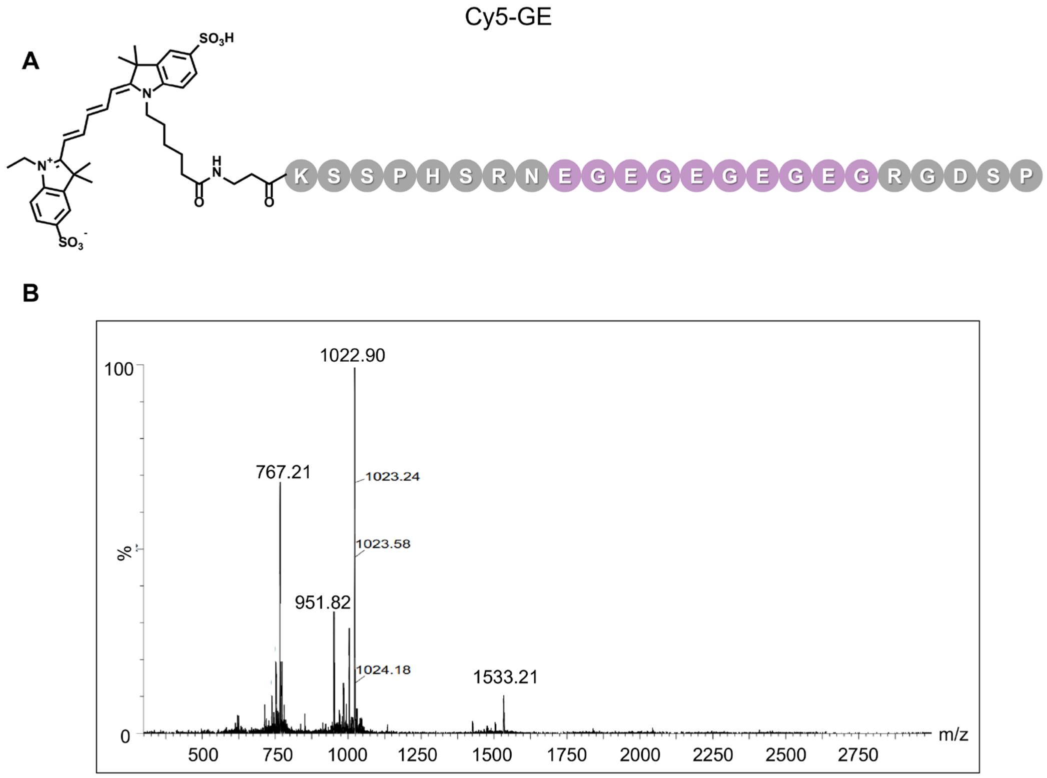


**Figure S8.** MS analysis of the Cy5-GE. The purity of Cy5-GE was greater than 90%.

A

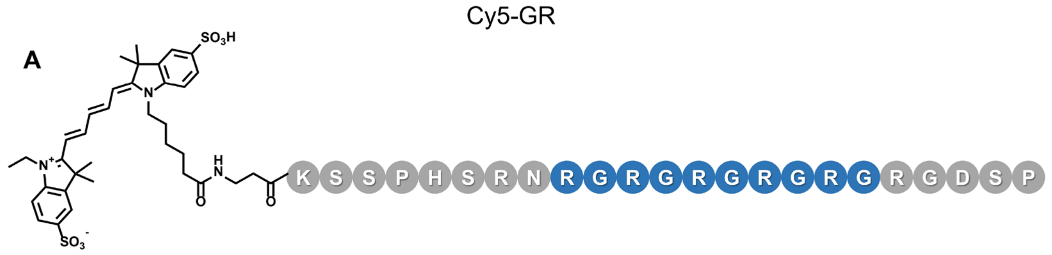


B

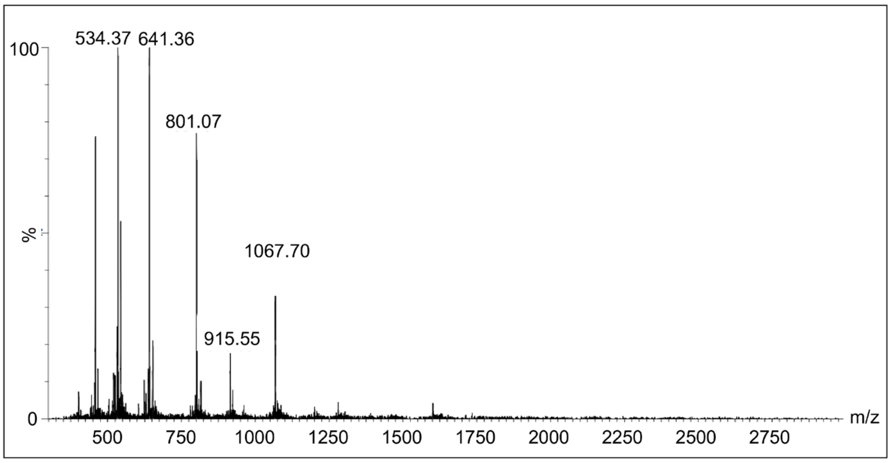


**Figure S9.** MS analysis of the Cy5-GR. The purity of Cy5-GR was greater than 90%.

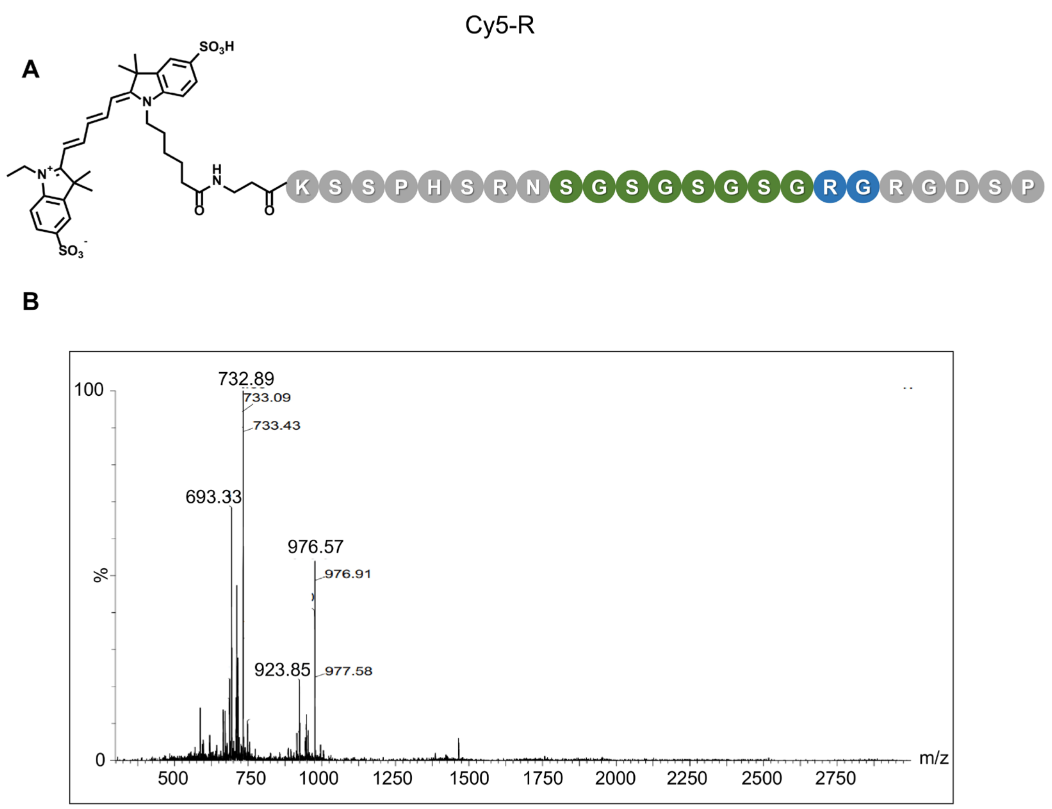


**Figure S10.** MS analysis of the Cy5-R. The purity of Cy5-R was greater than 90%.

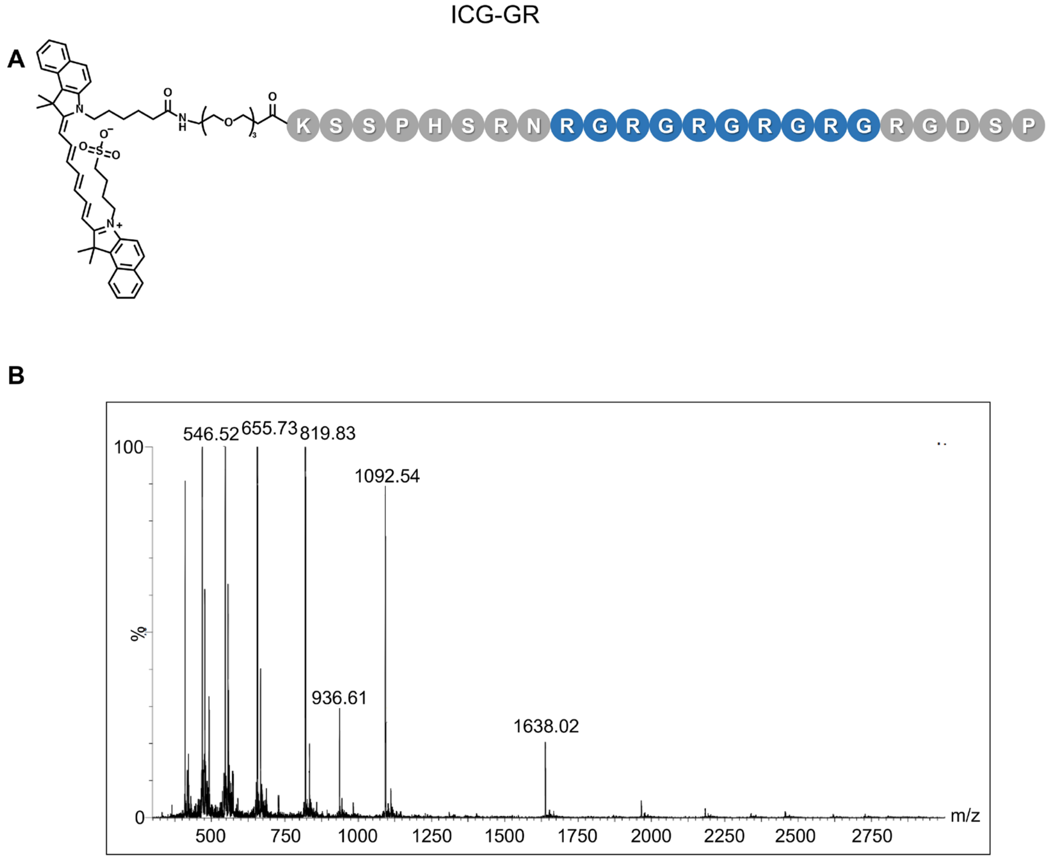


**Figure S11.** MS analysis of the ICG-GR. The purity of ICG-GR was greater than 90%.

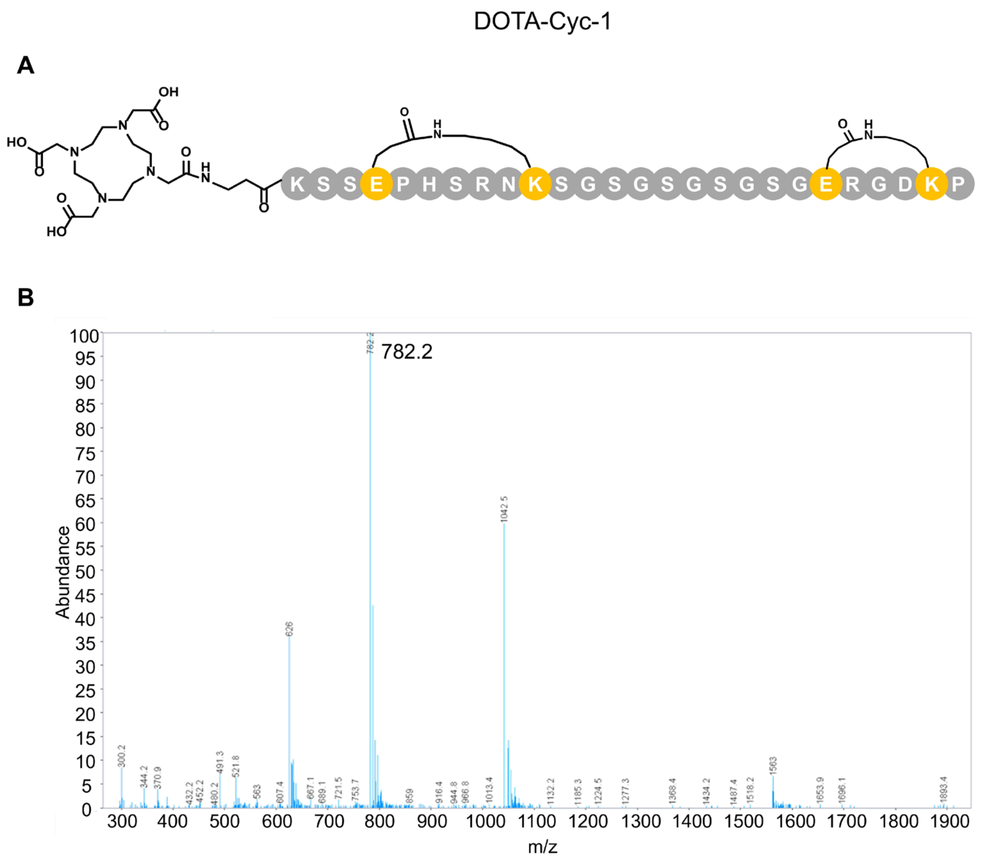


**Figure S12.** MS analysis of the DOTA-Cyc-1. The purity of DOTA-Cyc-1 was greater than 90%.

DOTA-Cyc-2

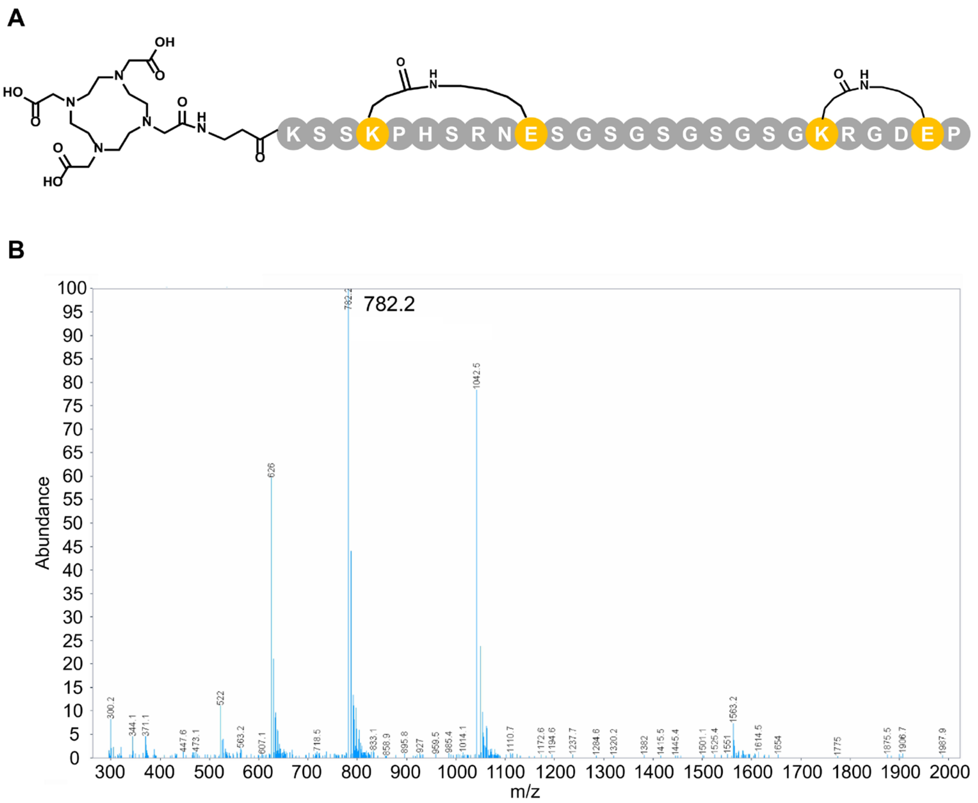


**Figure S13.** MS analysis of the DOTA-Cyc-2. The purity of DOTA-Cyc-2 was greater than 90%.

DOTA-Cyc-3

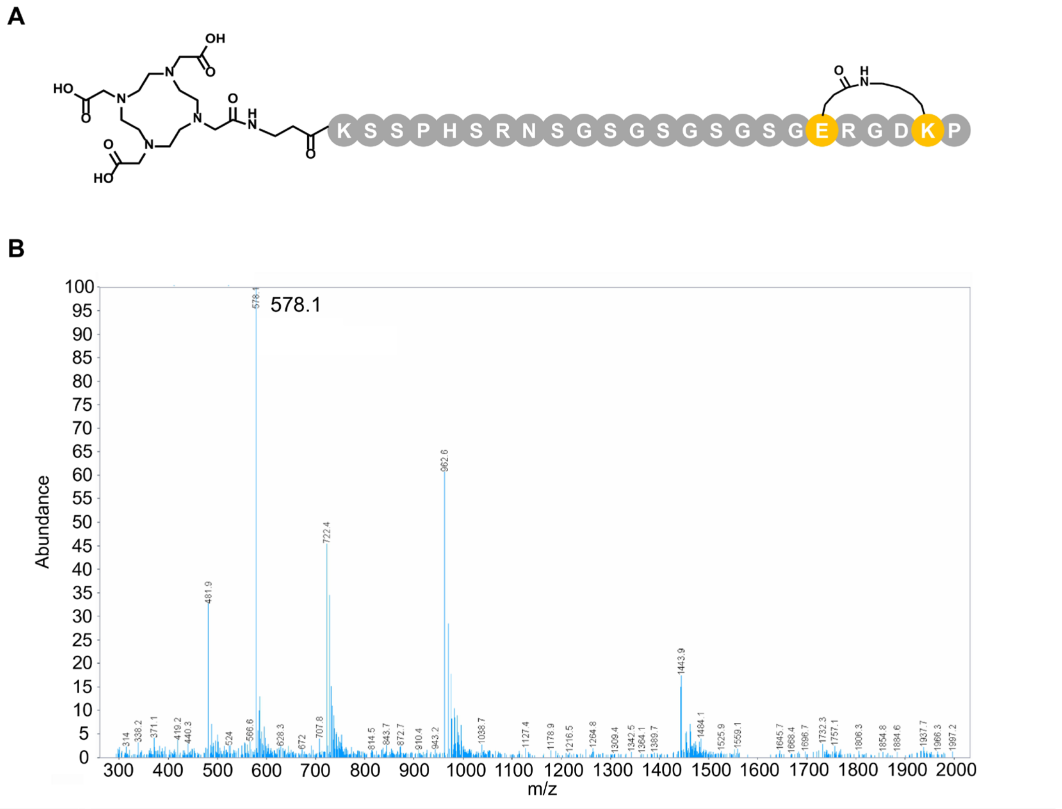


**Figure S14.** MS analysis of the DOTA-Cyc-3. The purity of DOTA-Cyc-3 was greater than 90%.

DOTA-Cyc-4

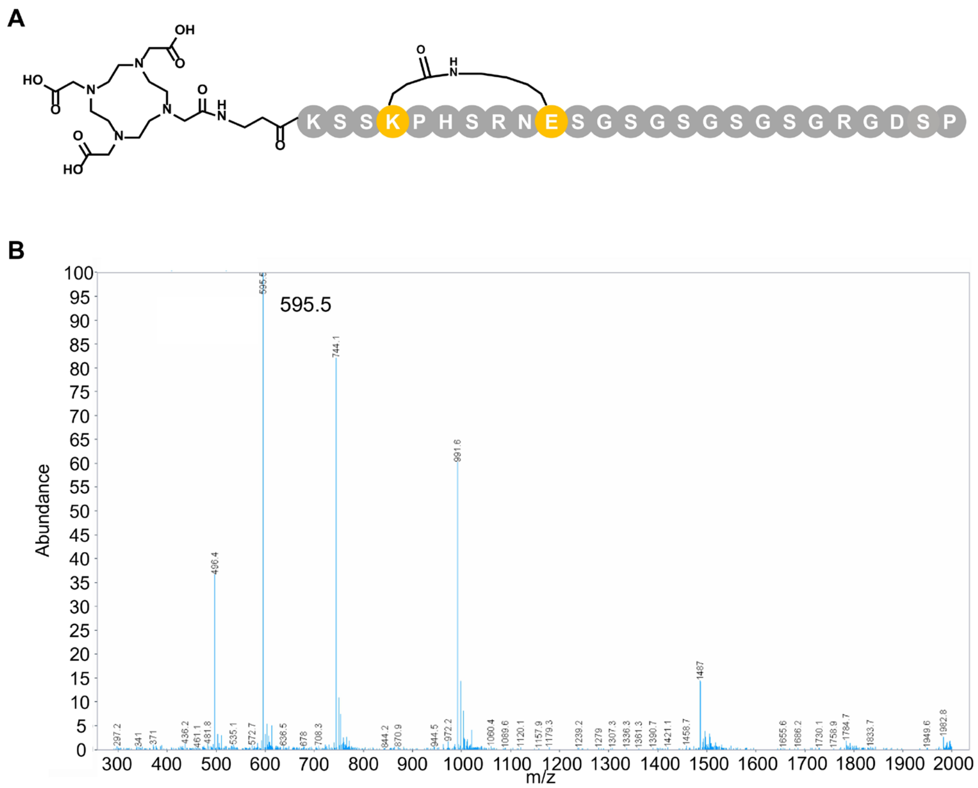


**Figure S15.** MS analysis of the DOTA-Cyc-4. The purity of DOTA-Cyc-4 was greater than 90%.

Cy5-Cyc-1

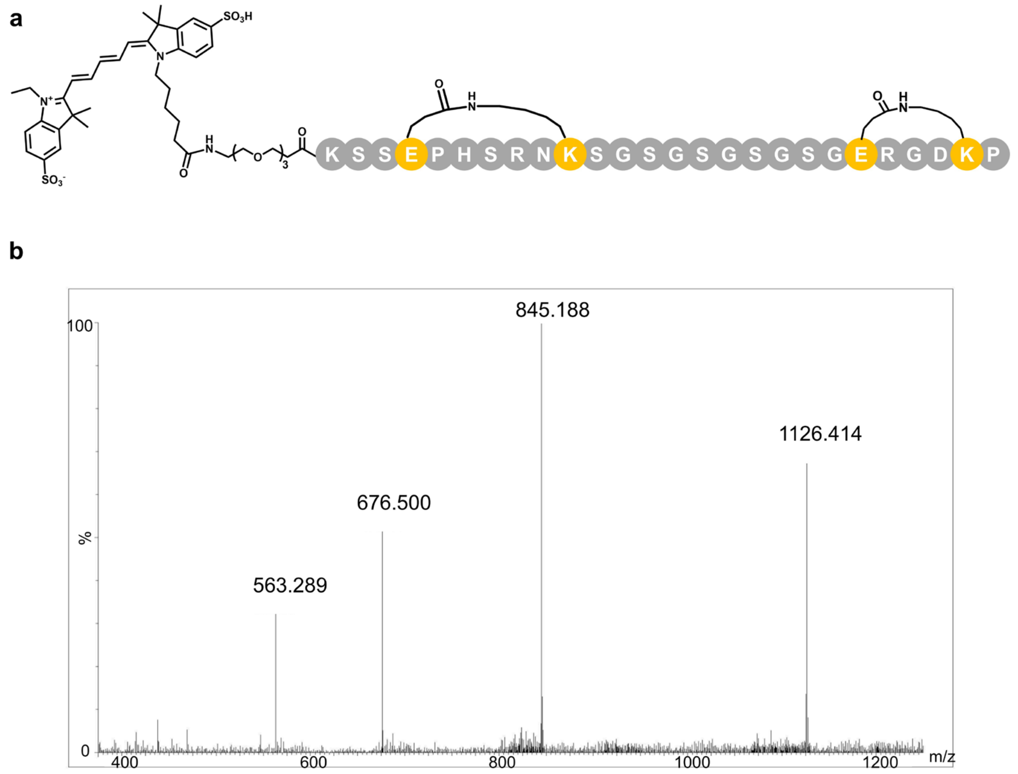


**Figure S16.** MS analysis of the Cy5-Cyc-1. The purity of Cy5-Cyc-1 was greater than 90%.

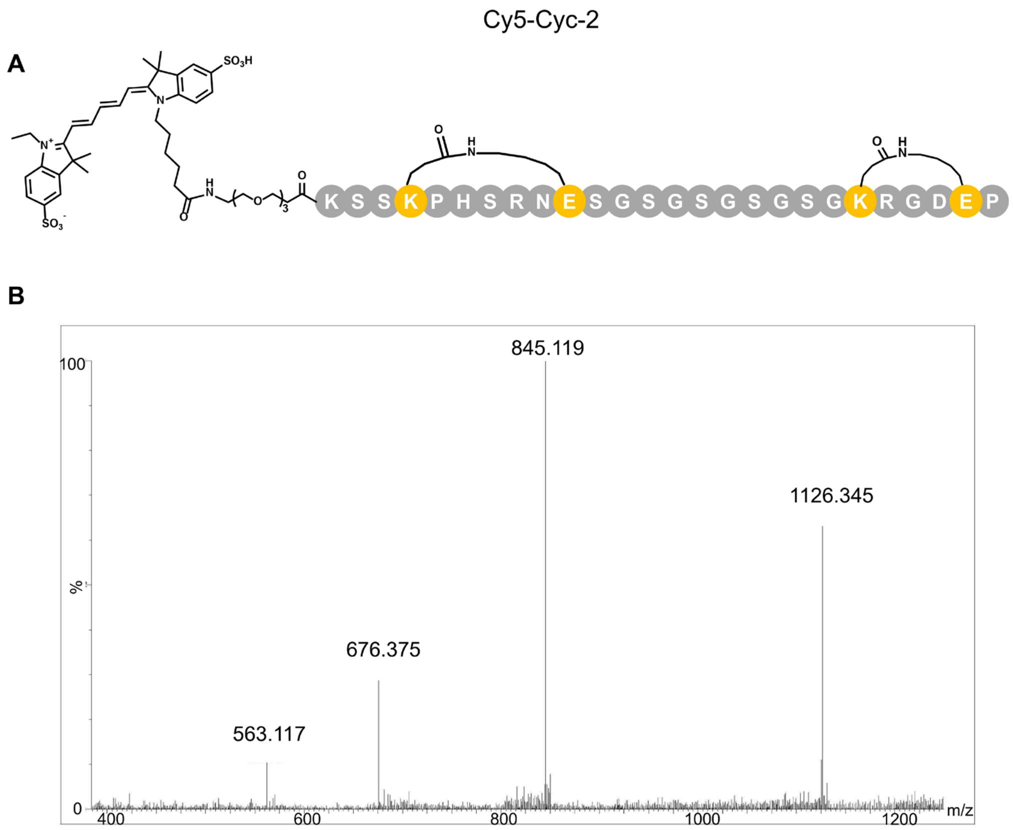


**Figure S17.** MS analysis of the Cy5-Cyc-2. The purity of Cy5-Cyc-2 was greater than 90%.

Cy5-Cyc-3

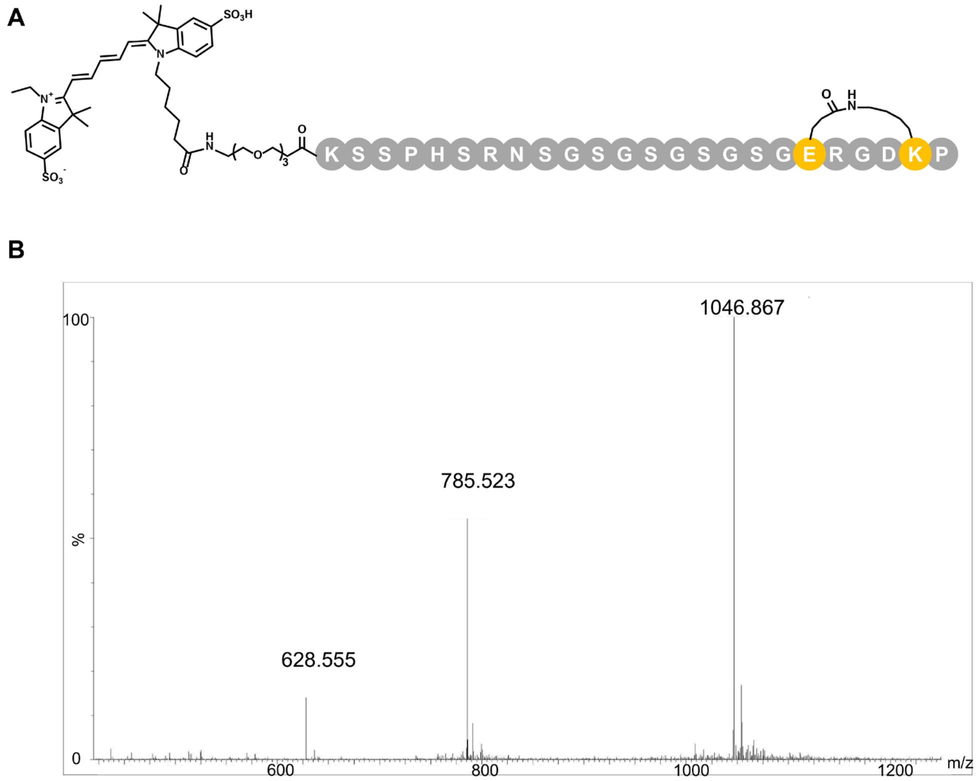


**Figure S18.** MS analysis of the Cy5-Cyc-3. The purity of Cy5-Cyc-3 was greater than 90%.

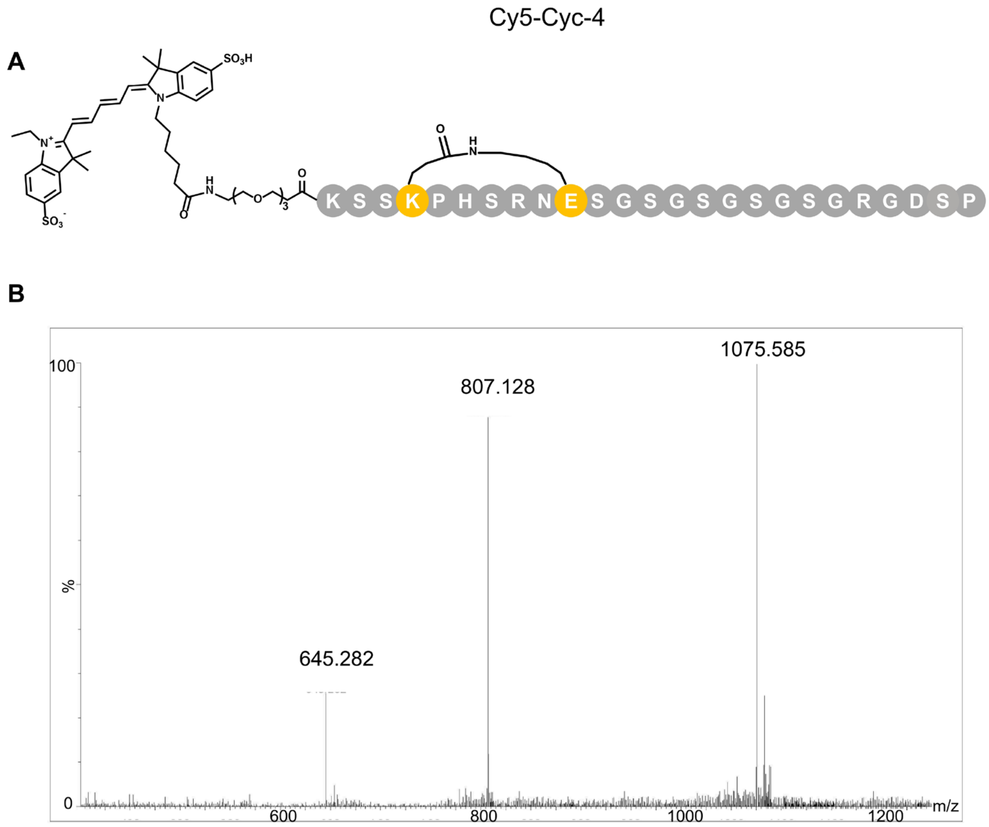


**Figure S19.** MS analysis of the Cy5-Cyc-4. The purity of Cy5-Cyc-4 was greater than 90%.

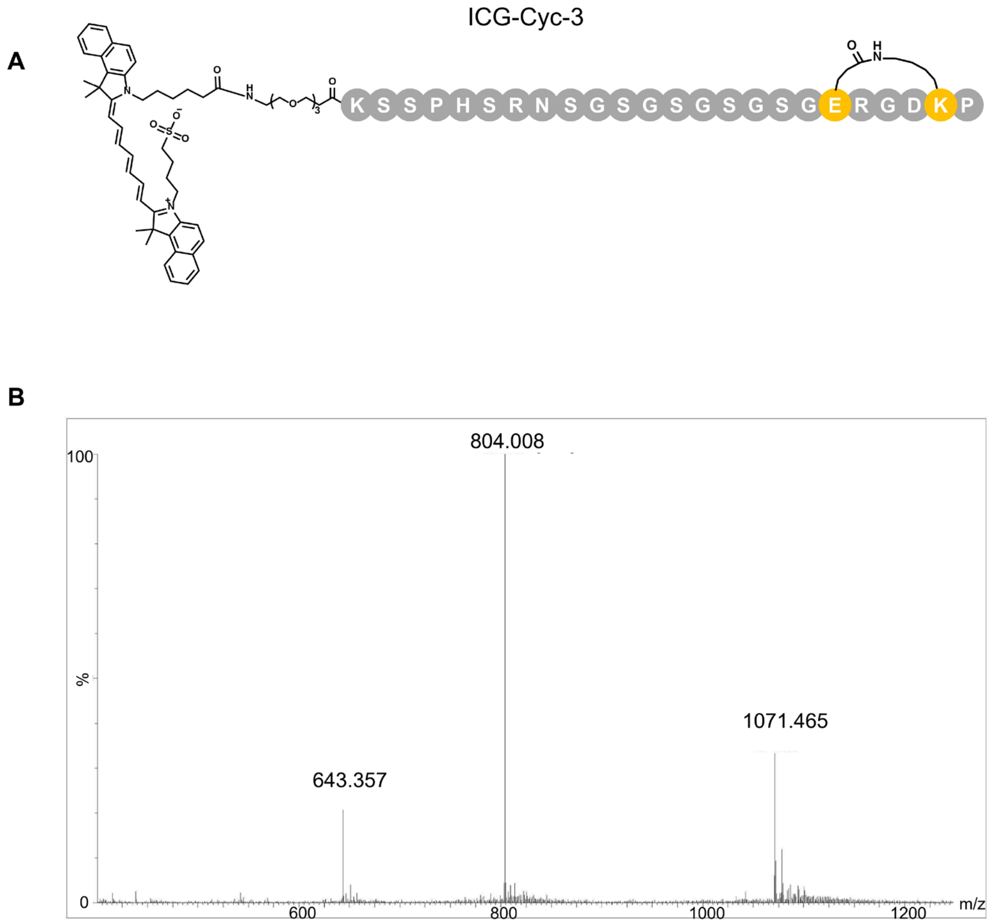


**Figure S20.** MS analysis of the ICG-Cyc-3. The purity of ICG-Cyc-3 was greater than 90%.

DOTA-PEG-1

**a**

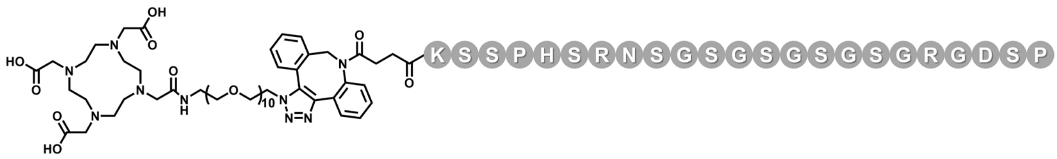


**b**

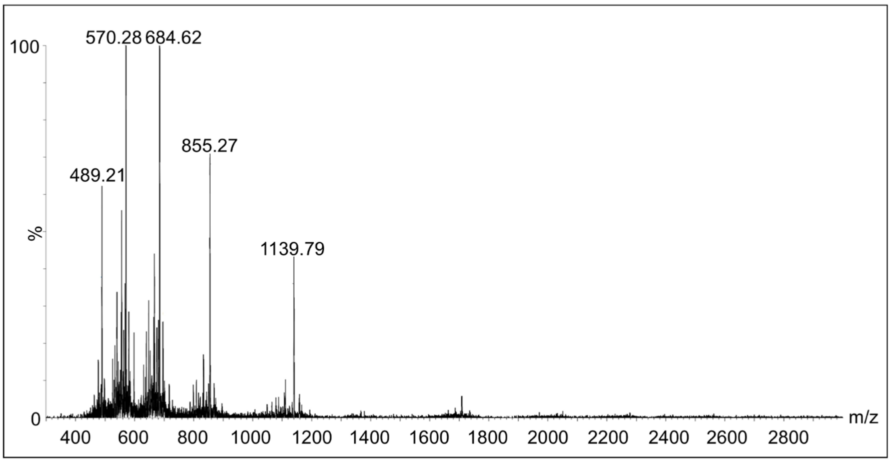


**Figure S21.** MS analysis of the DOTA-PEG-1. The purity of DOTA-PEG-1 was greater than 90%.

DOTA-PEG-2

A

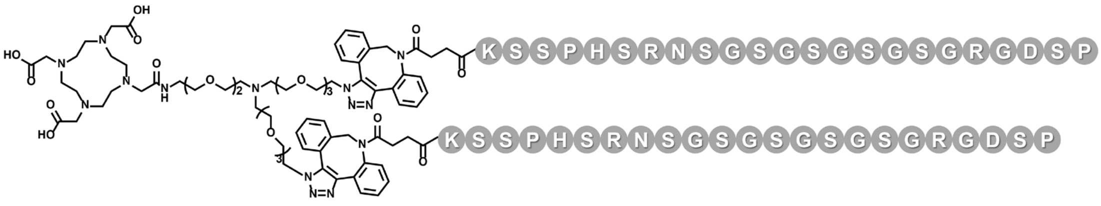


B

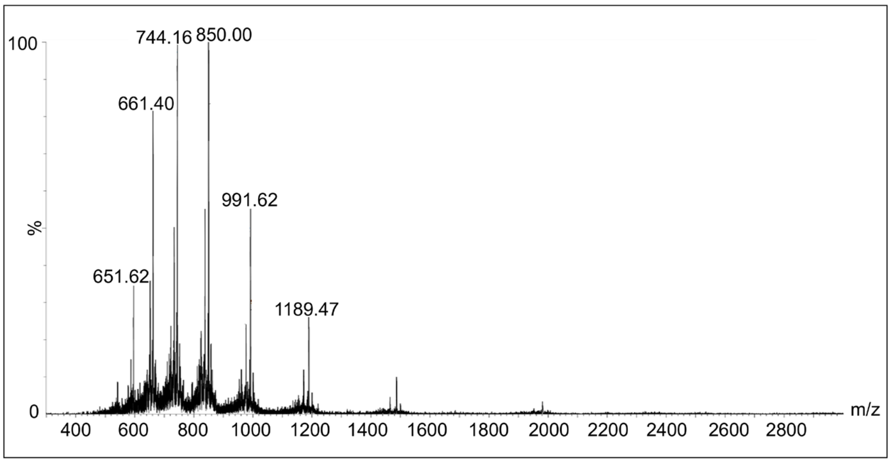


**Figure S22.** MS analysis of the DOTA-PEG-2. The purity of DOTA-PEG-2 was greater than 90%.

DOTA-PEG-3

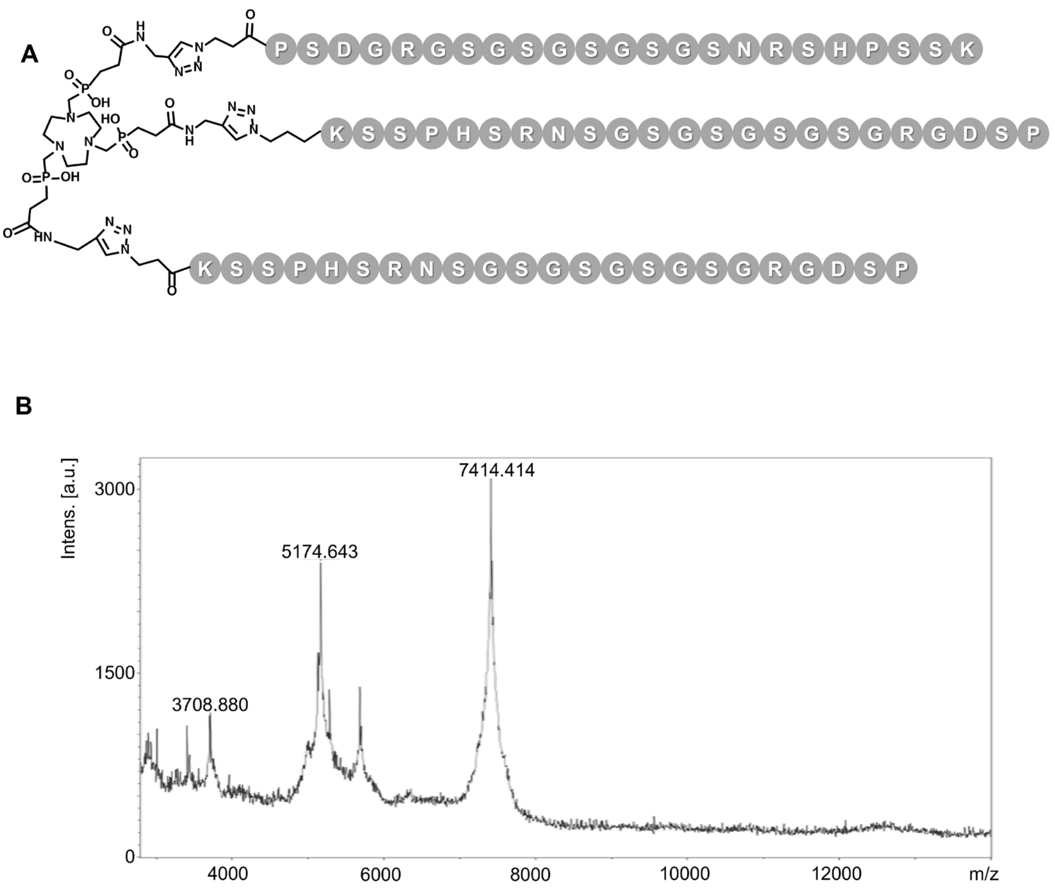


**Figure S23.** MS analysis of the DOTA-PEG-3. The purity of DOTA-PEG-3 was greater than 90%.

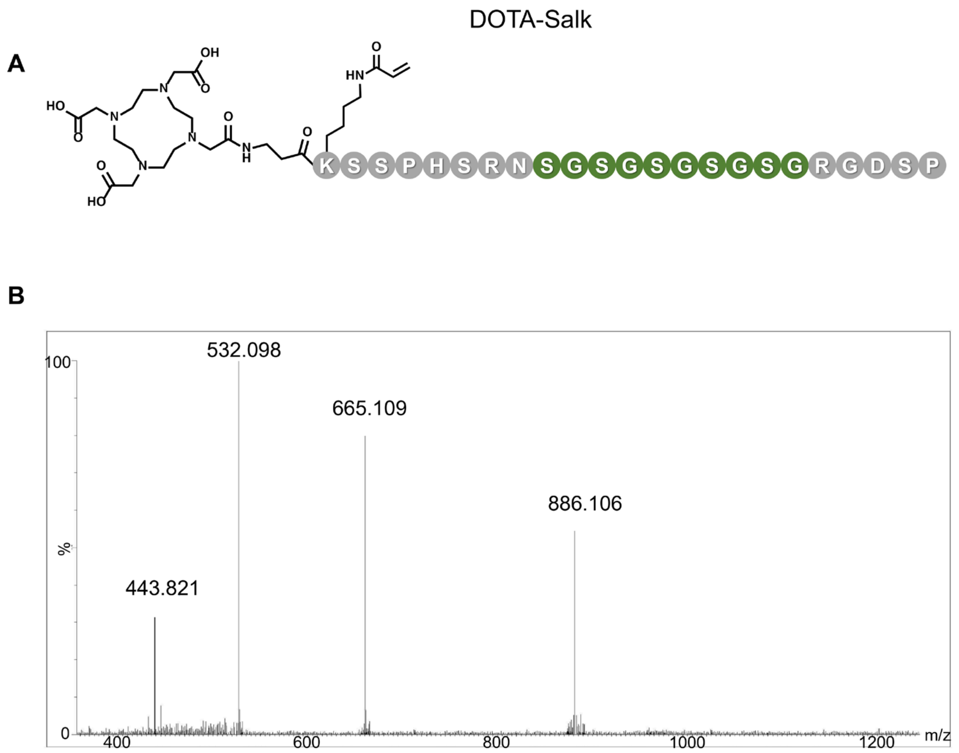


**Figure S24.** MS analysis of the DOTA-Salk. The purity of DOTA-Salk was greater than 90%.

A

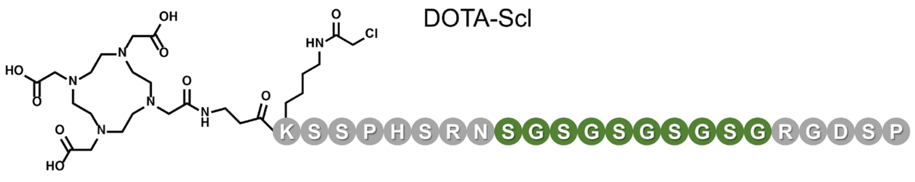


B

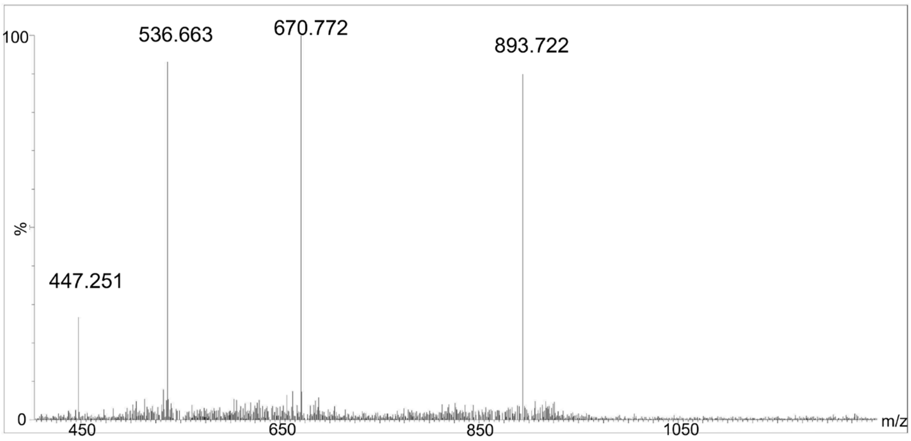


**Figure S25.** MS analysis of the DOTA-Scl. The purity of DOTA-Scl was greater than 90%.

DOTA-Ssf

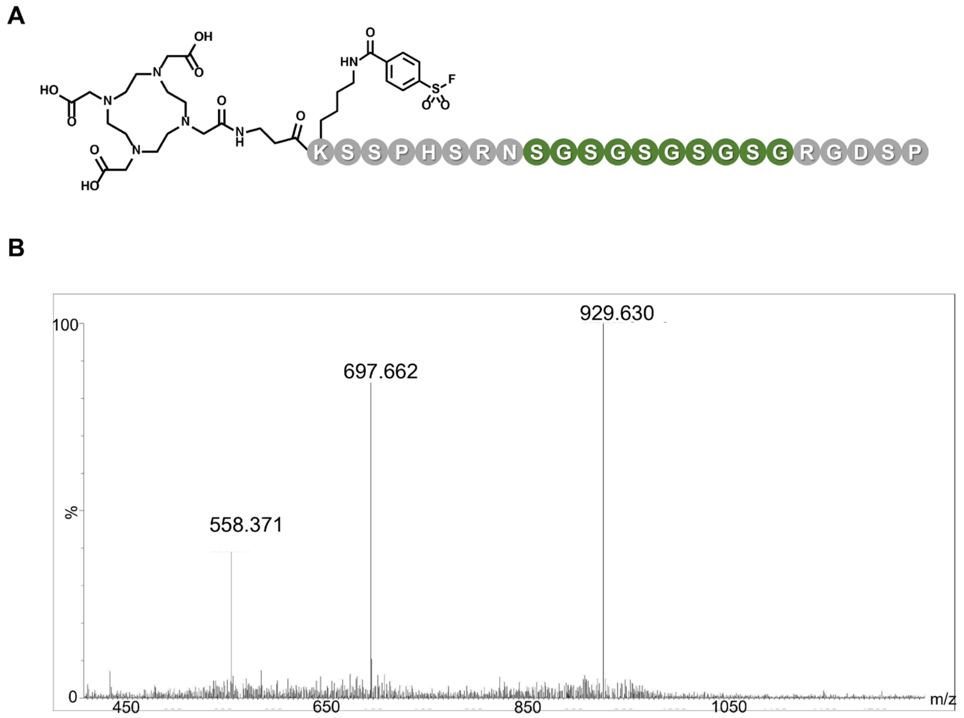


**Figure S26.** MS analysis of the DOTA-Ssf. The purity of DOTA-Ssf was greater than 90%.

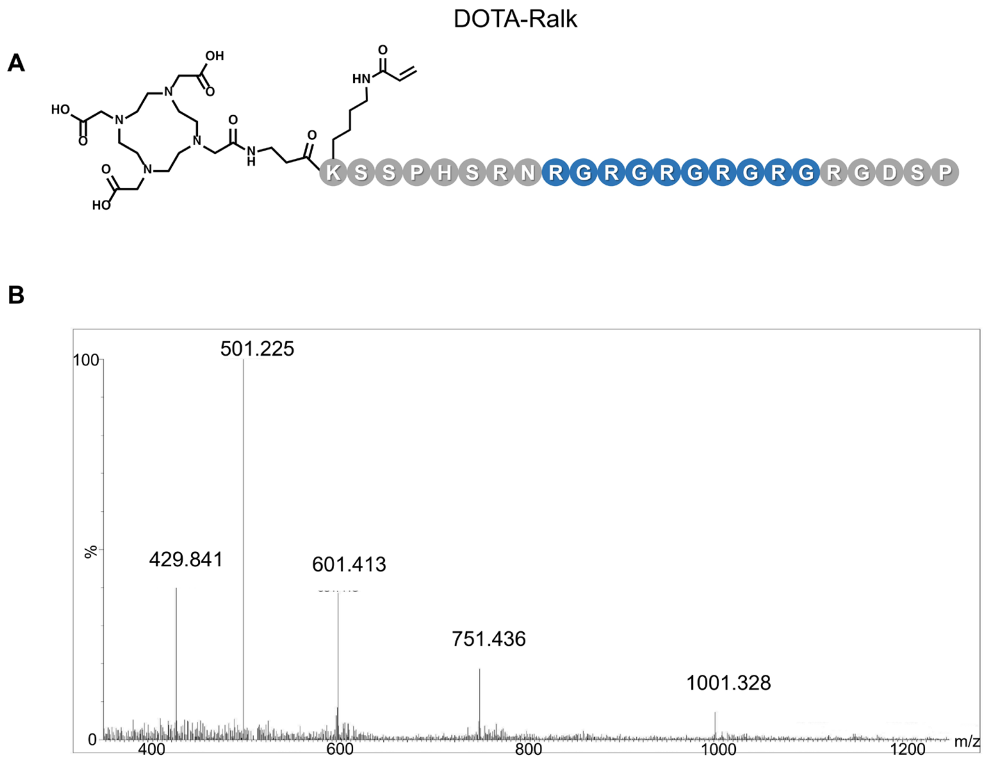


**Figure S27.** MS analysis of the DOTA-Ralk. The purity of DOTA-Ralk was greater than 90%.

DOTA-Rcl

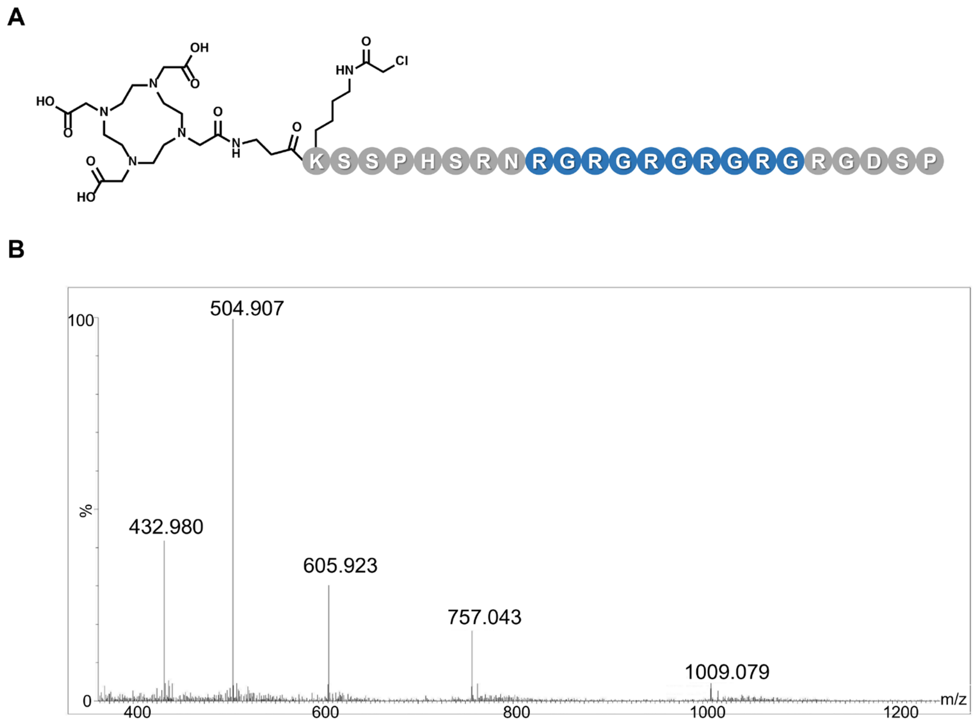


**Figure S28.** MS analysis of the DOTA-Rcl. The purity of DOTA-Rcl was greater than 90%.

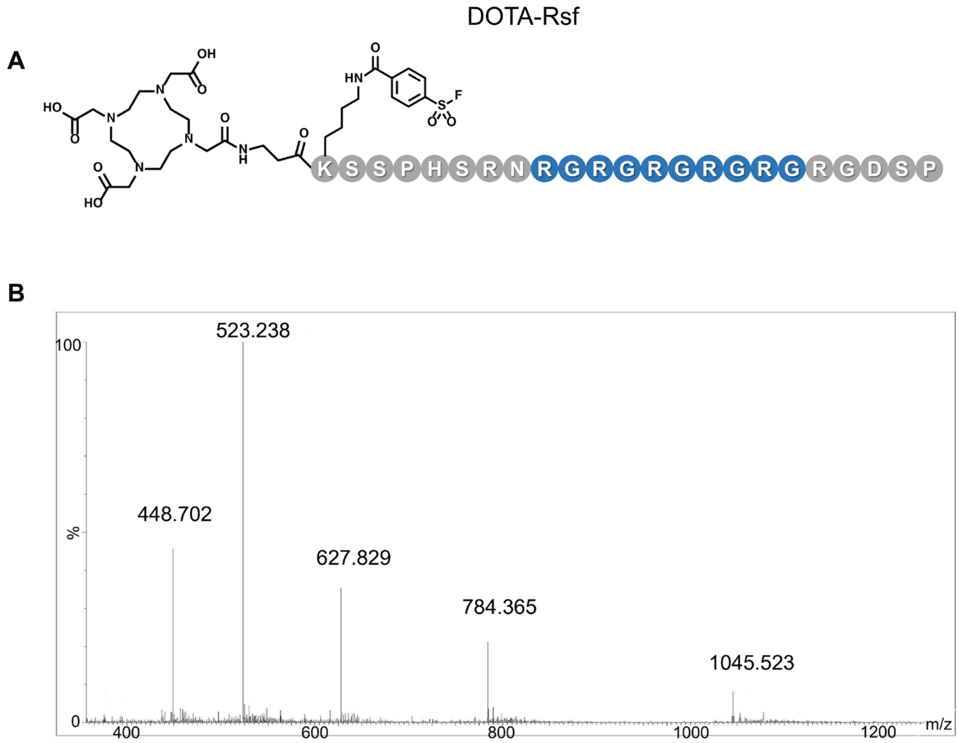


**Figure S29.** MS analysis of the DOTA-Rsf. The purity of DOTA-Rsf was greater than 90%.

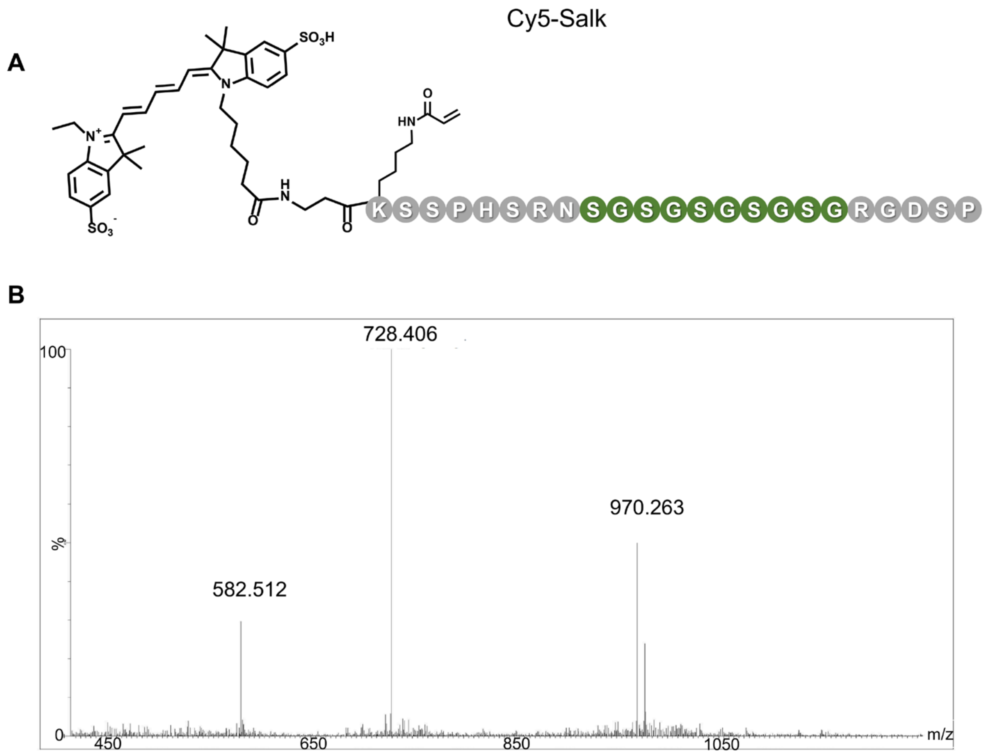


**Figure S30.** MS analysis of the Cy5-Salk. The purity of Cy5-Salk was greater than 90%.

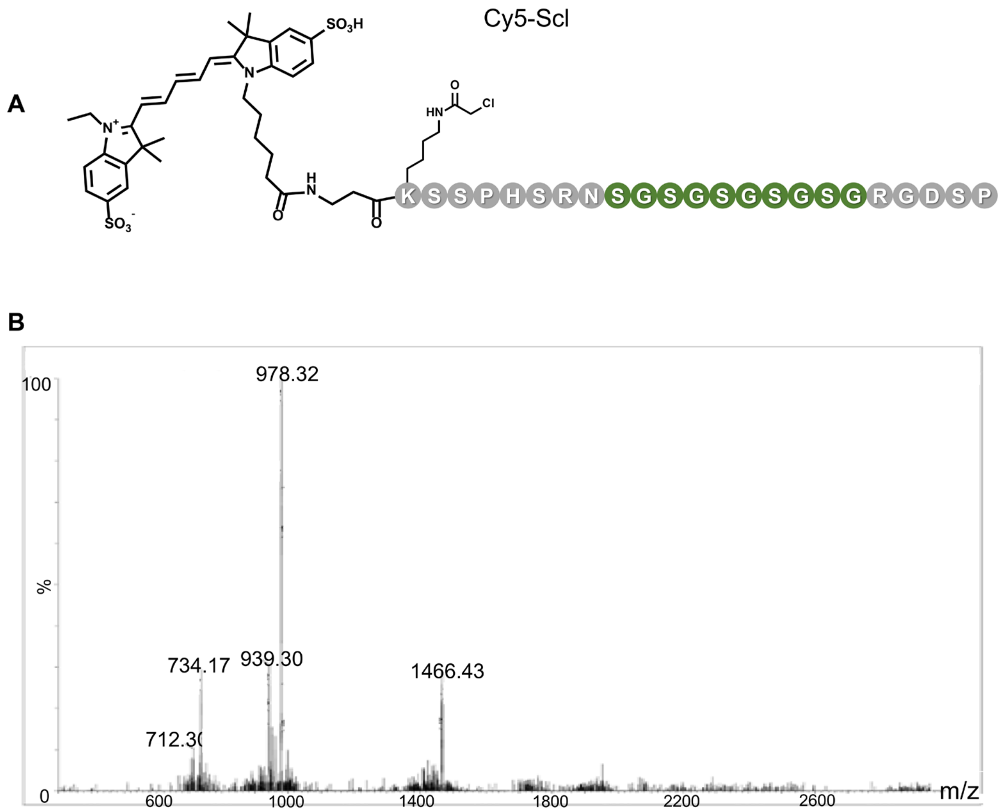


**Figure S31.** MS analysis of the Cy5-Scl. The purity of Cy5-Scl was greater than 90%.

A

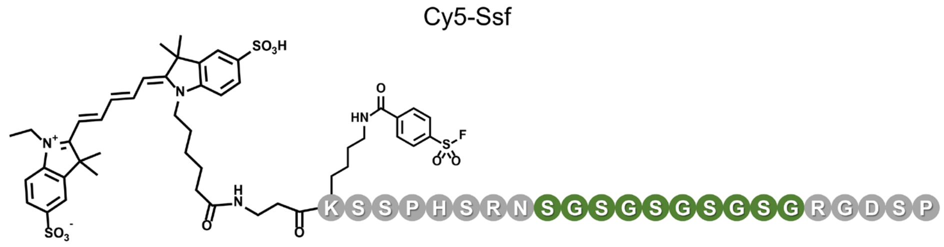


B

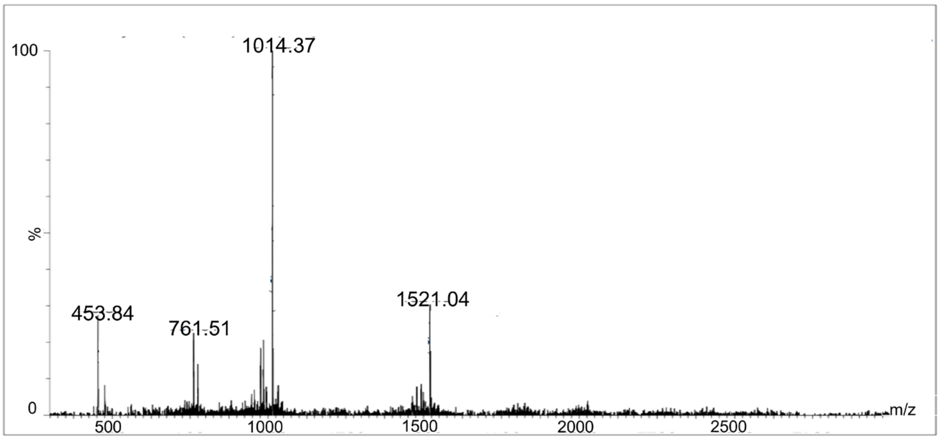


**Figure S32.** MS analysis of the Cy5-Ssf. The purity of Cy5-Ssf was greater than 90%.

Cy5-Ralk

A

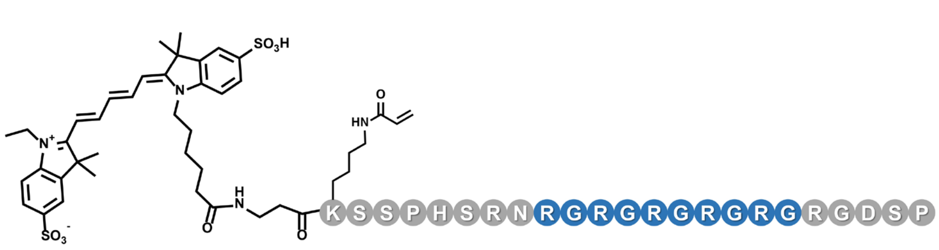


B

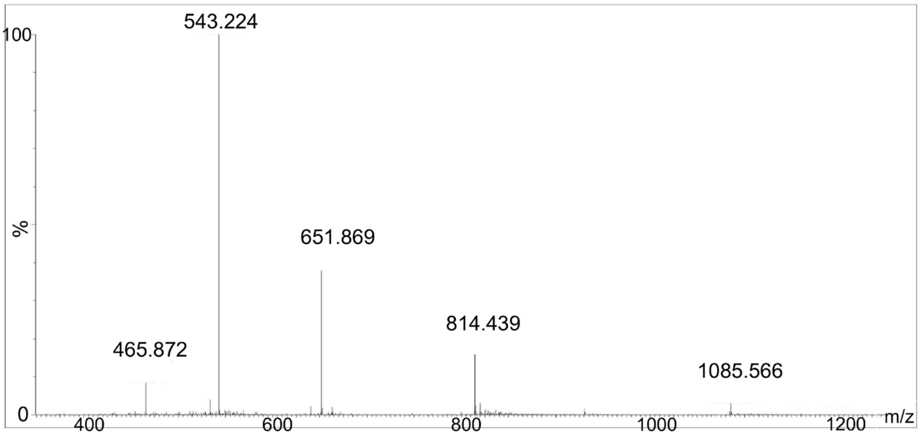


**Figure S33.** MS analysis of the Cy5-Ralk. The purity of Cy5-Ralk was greater than 90%.

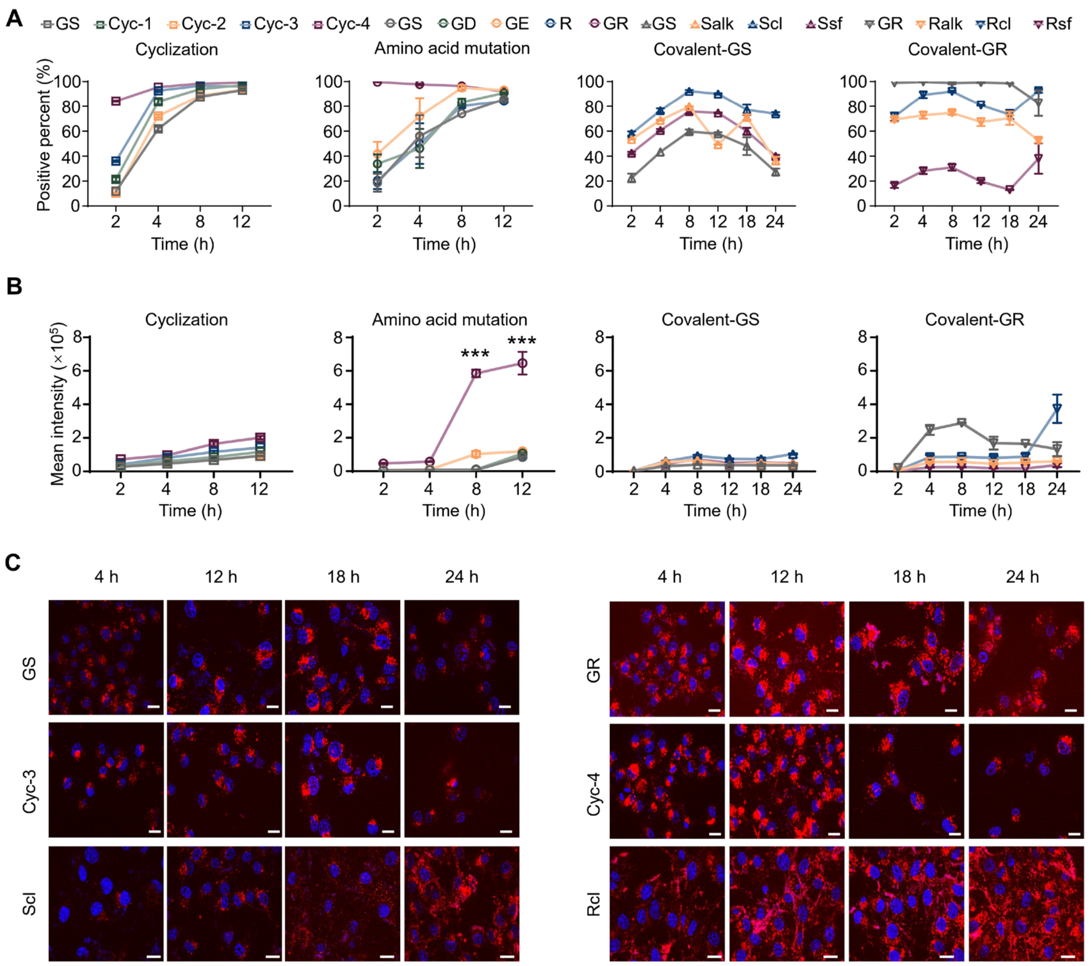


**Figure S34. In vitro evaluation of cell penetration of Cy5-labeled peptides.** (**A-B**) In vitro binding of peptides with integrin α5β1 in U87MG cell lines by flow cytometry. The positive percent (**A**) and mean fluorescence (**B**) after treated with cyclization, amino acid mutation, and covalent modification peptides for 2, 4, 8, 12, and 24 h. (**C**) The confocal imaging of Cy5-GS, Cy5-GR, Cy5-Cyc-3, Cy5-Cyc-4, Cy5-Scl, and Cy5-Rcl after added in U87MG cells for 4, 12, 18, and 24 h. Scale bar, 10 μm. All results are expressed as means ± SEM, as indicated in at least three independent experiments. A multiple t-test was used when two groups were compared. The symbol "*" represents differences compared with the Cy5-GS. *** $p < 0.001$.

Confocal imaging of Cy5-labeled peptides in U87MG cells.

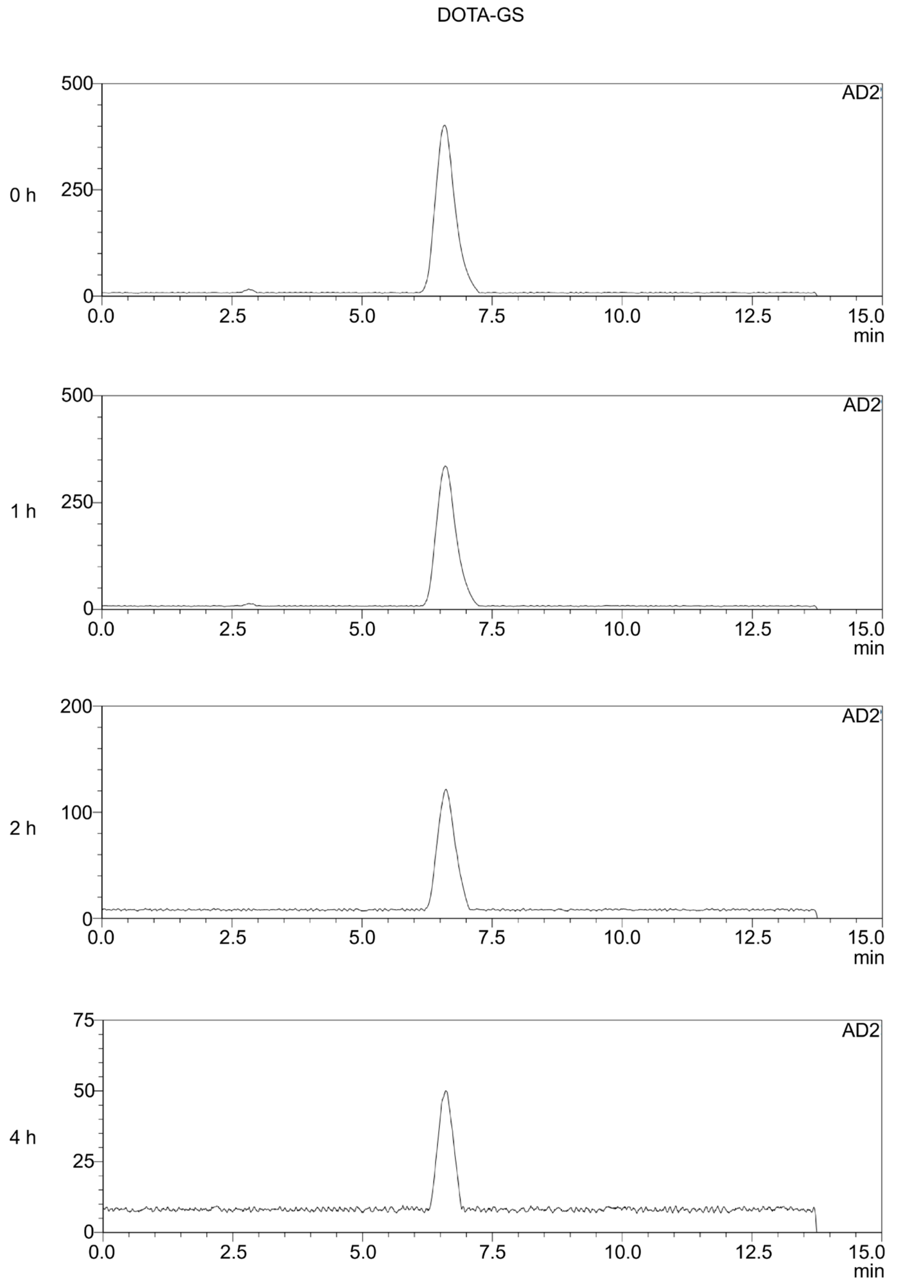


**Figure S35.** *In vitro* Stability of [$^{68}$Ga]GS in saline.

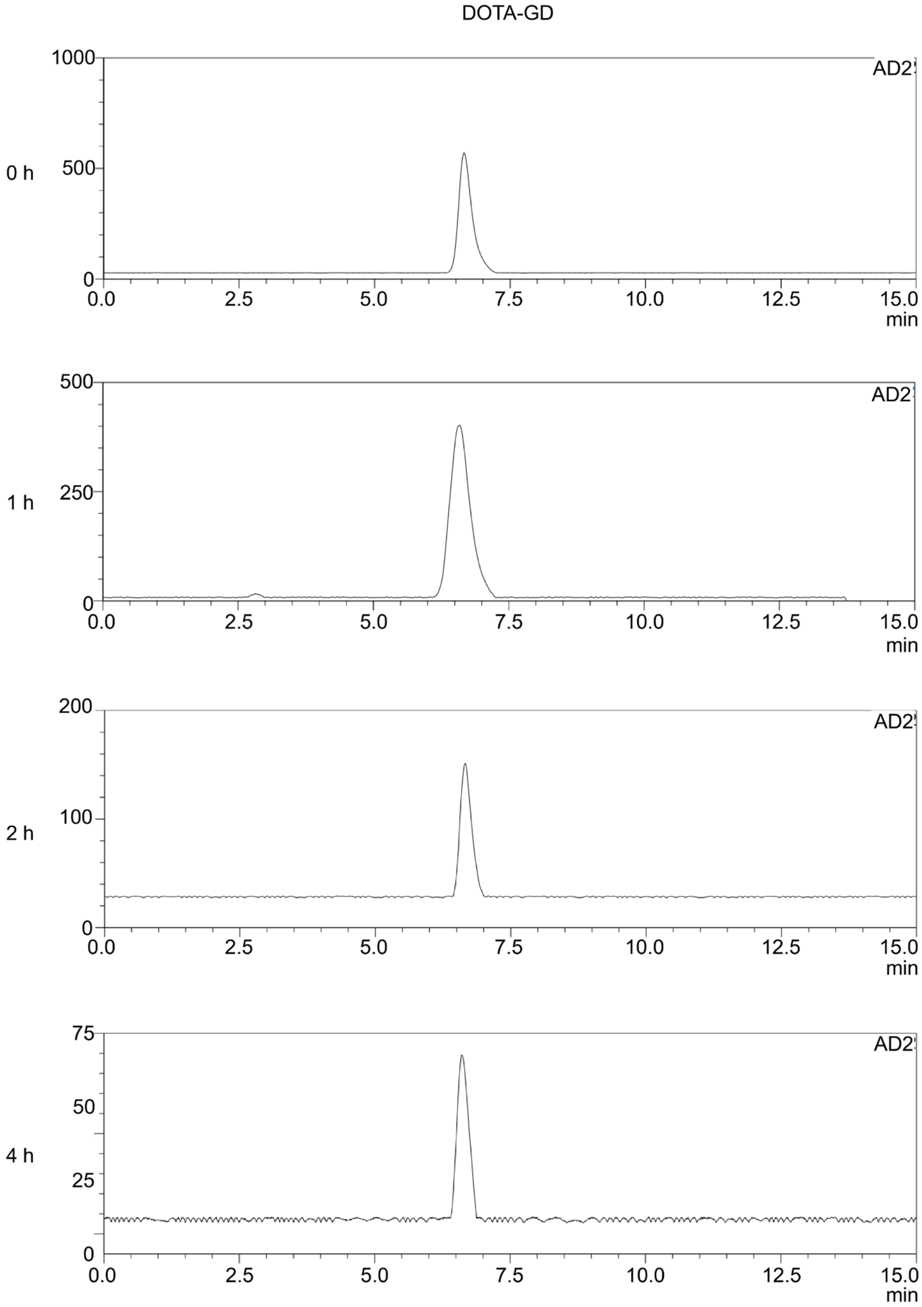


**Figure S36.** *In vitro* Stability of [$^{68}$Ga]GD in saline.

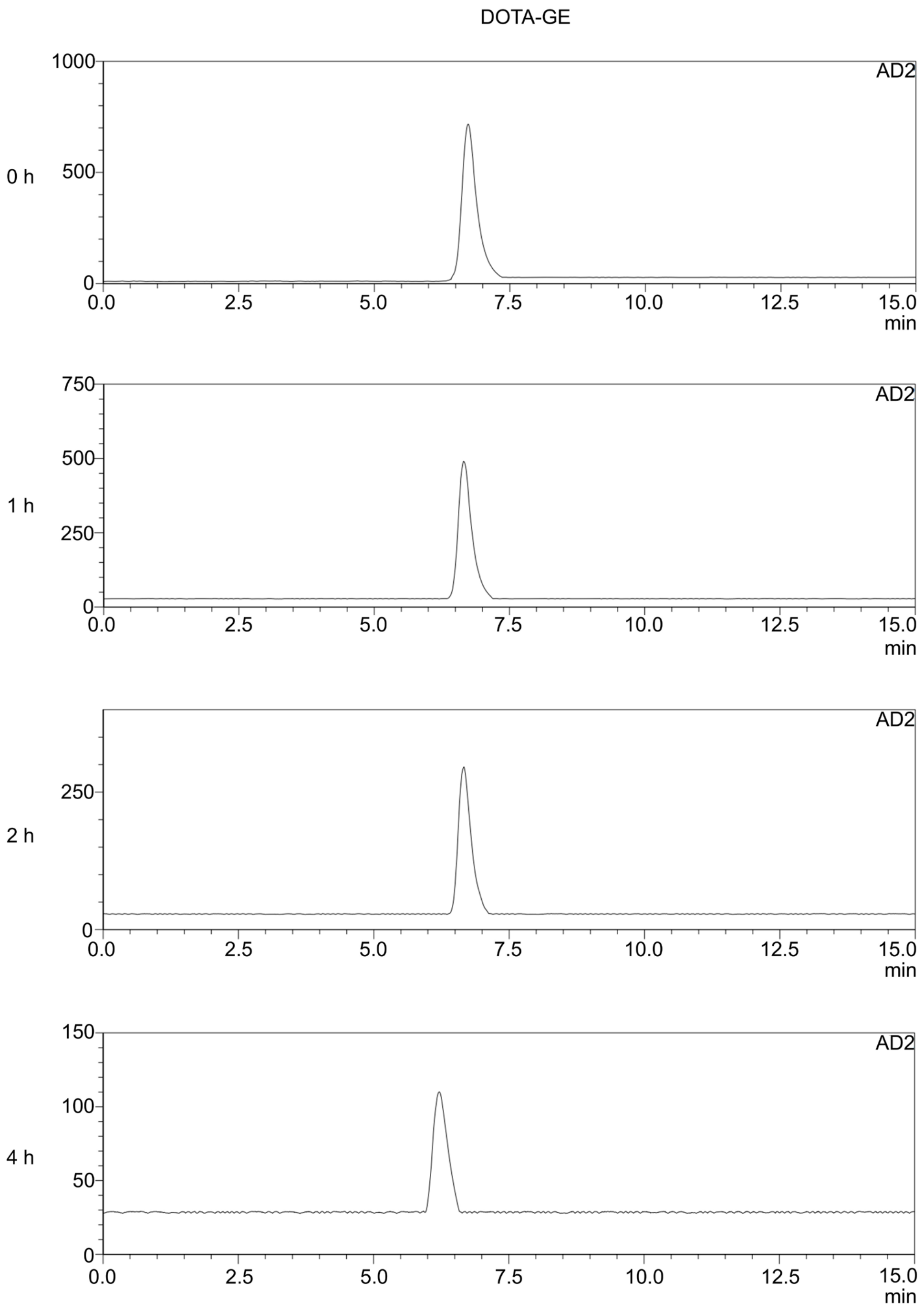


**Figure S37.** *In vitro* Stability of [$^{68}$Ga]GE in saline.

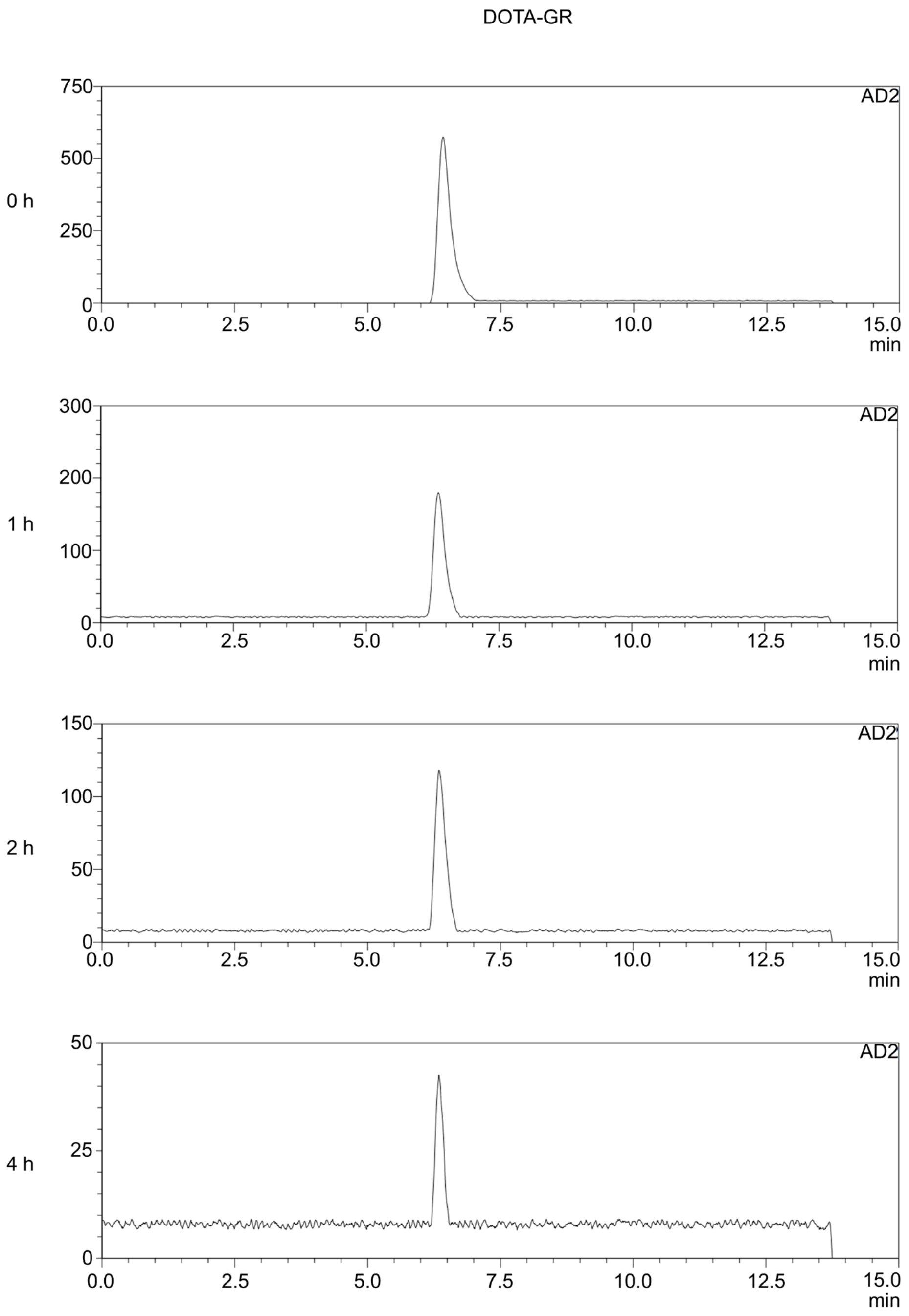


**Figure S38.** *In vitro* Stability of [$^{68}$Ga]GR in saline.

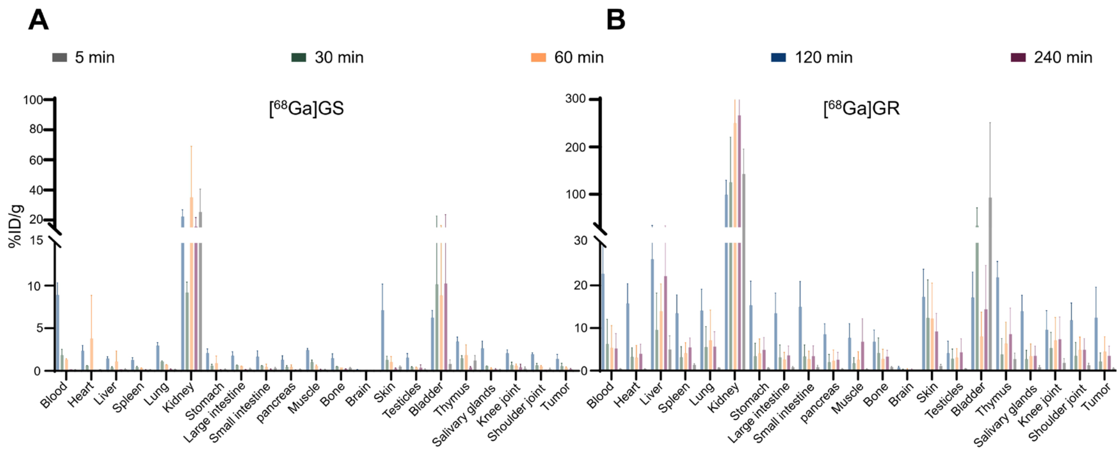


**Figure S39.** *Ex vivo* biodistribution study of [$^{68}$Ga]GR and [68Ga]GS in normal mice. **(A-B)** *Ex vivo* biodistribution analysis of [$^{68}$Ga]GS (a) and [$^{68}$Ga]GR (b) in normal mice at 5, 30, 60, 120, and 240 min p.i. (n = 3). Each mouse was injected with 1.85 MBq radiopharmaceutical, and the blood and major organs were collected at each time point. Blood, heart, liver, and kidney uptake of [$^{68}$Ga]GR and [$^{68}$Ga]GS based on ex vivo biodistribution. All results are expressed as means ± SEM, as indicated in at least three independent experiments.

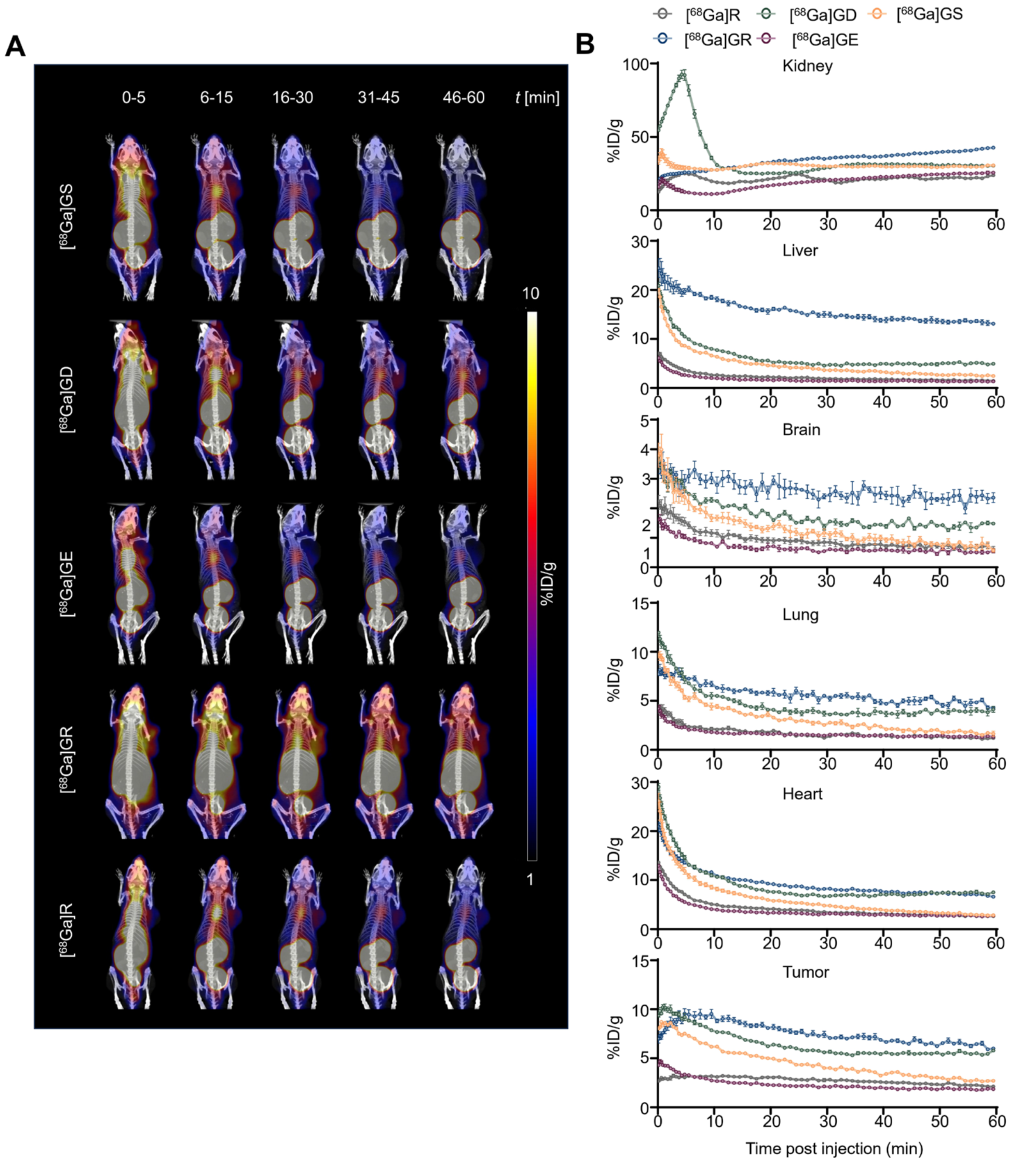


**Figure S40.** Dynamic PET/CT imaging in tumor-bearing mice. Representative dynamic PET/CT images **(A)** and quantitative analysis of uptake in major organs **(B)** in U87MG-bearing mouse for continuous 1 hour after intravenous injection of approximately 0.1 mCi [$^{68}$Ga]GS, [$^{68}$Ga]GD, [$^{68}$Ga]GE, [$^{68}$Ga]GR, and [$^{68}$Ga]R in saline. All results are expressed as means ± SEM, as indicated in at least three independent experiments.

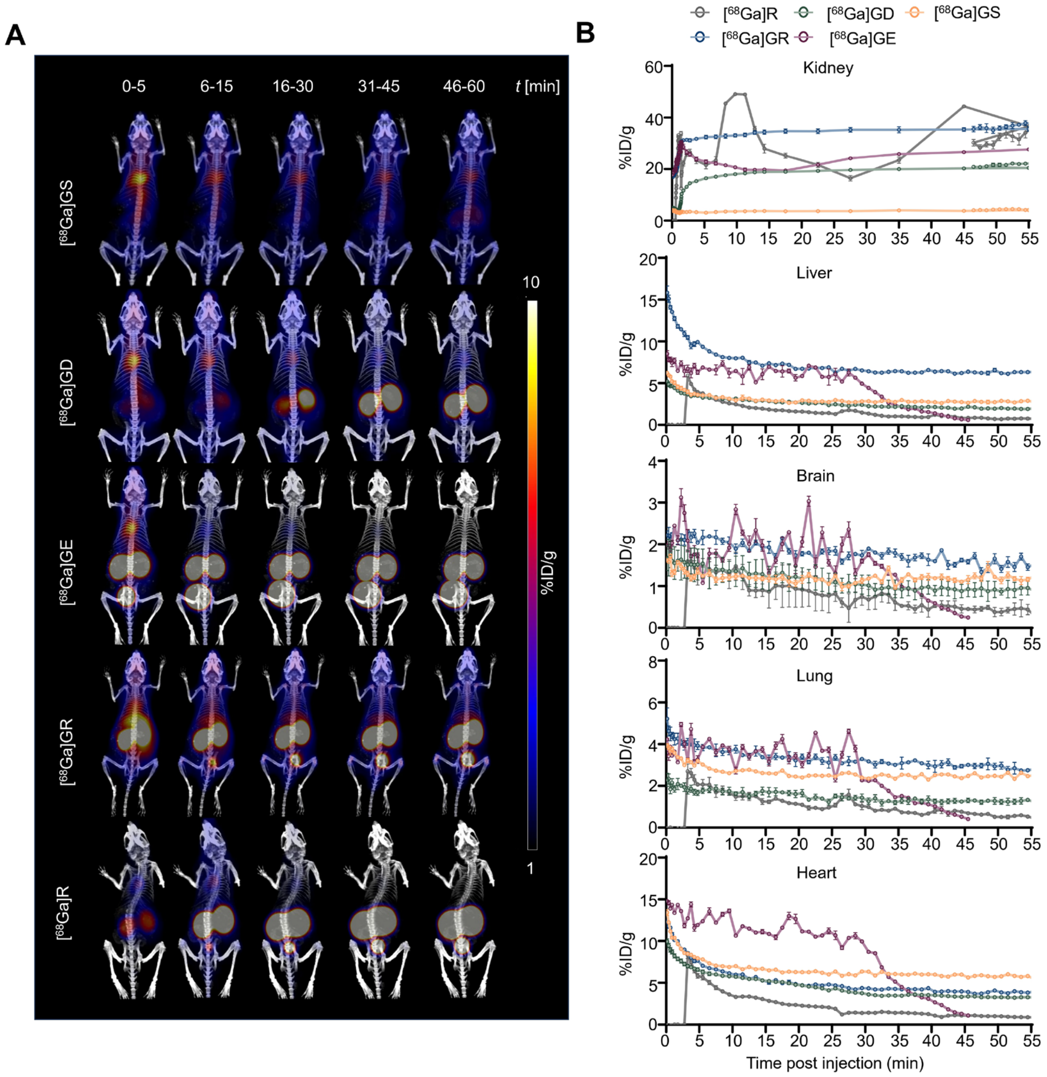


**Figure S41.** Dynamic PET/CT imaging in normal mice. Representative dynamic PET/CT images **(A)** and quantitative analysis of uptake in major organs **(B)** in normal mouse for continuous 1 hour after intravenous injection of approximately 0.1 mCi [$^{68}$Ga]GS, [$^{68}$Ga]GD, [$^{68}$Ga]GE, [$^{68}$Ga]GR, and [$^{68}$Ga]R in saline. All results are expressed as means ± SEM, as indicated in at least three independent experiments.

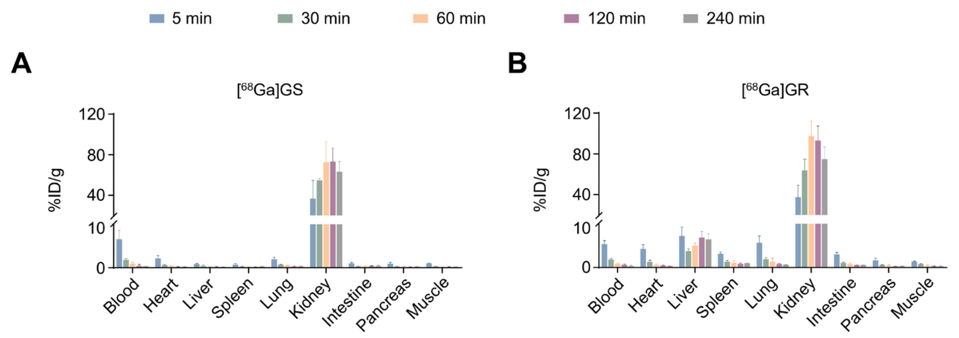


**Figure S42.** *Ex vivo* biodistribution study in normal mice. a-b, *Ex vivo* biodistribution of [$^{68}$Ga]GS **(A)** and [$^{68}$Ga]GR **(B)** in normal mice. A dose of 1.85 MBq was injected intravenously into each mouse. The mice were sacrificed at 5 min, 30 min, 1h, 2h, and 4h after injection (n = 3). All results are expressed as means ± SEM, as indicated in at least three independent experiments.

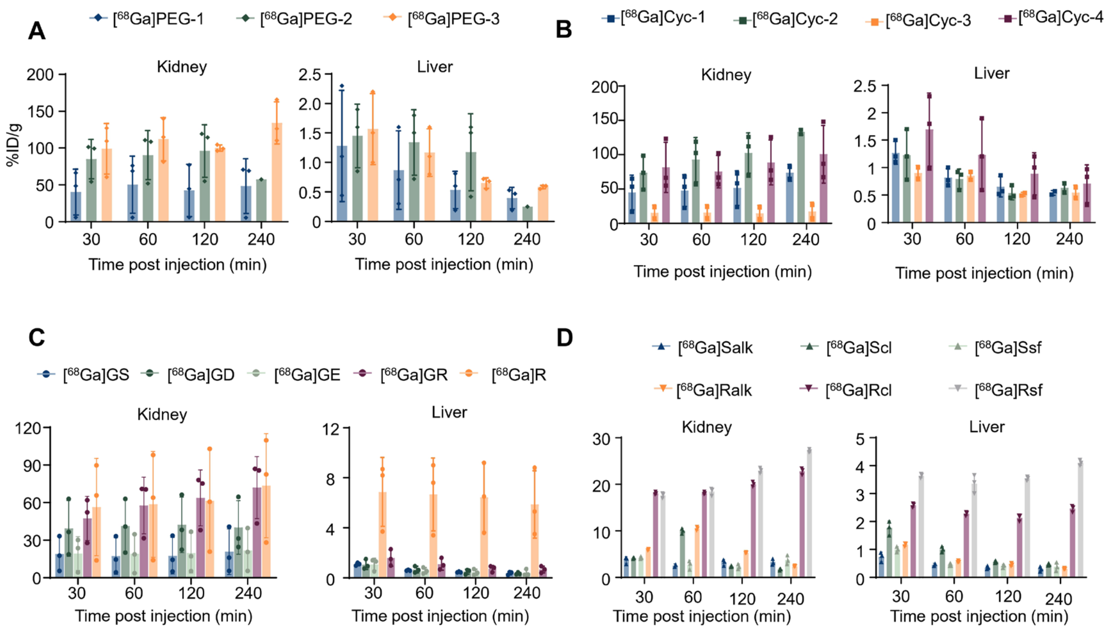


**Figure S43.** Quantification of major organ accumulation based on the PET/CT imaging of $^{68}$Ga-labeling peptides. A dose of 1.85 MBq was injected intravenously into each mouse. All results are expressed as means ± SEM, as indicated in at least three independent experiments.

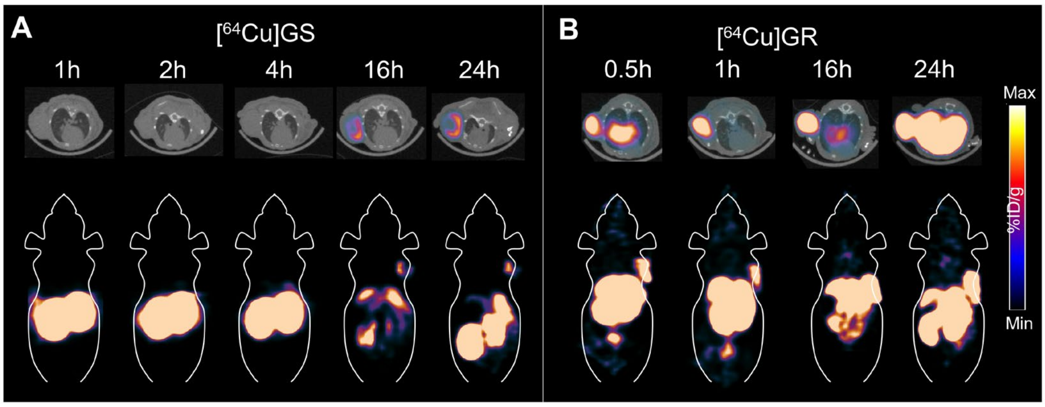


**Figure S44.** PET/CT imaging of [$^{64}$Cu]GS and [$^{64}$Cu]GR in U87MG tumor-bearing mice. A dose of 1.85 MBq was injected intravenously into each mouse.

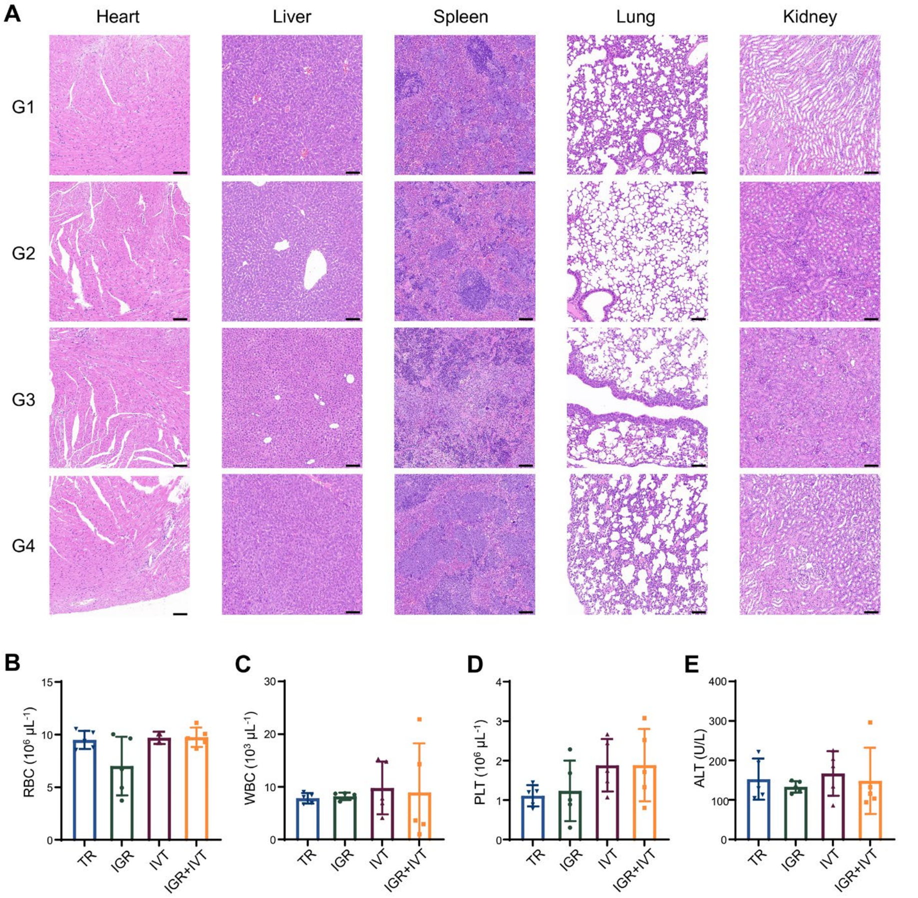


**Figure S45.** Safety analysis. **(A)** G1, G2, G3 and G4 safety evaluation of main organs (heart, liver, spleen, lung and kidney) in U87MG tumor-bearing mice. Scale bar: 100 μM. Hematological analysis: **(B)** RBC (red blood cell), **(C)** WBC (white blood cell), **(D)** PLT (platelet) and **(E)** ALT (alanine aminotransferase). All results are expressed as means ± SEM, as indicated in at least three independent experiments.

**Table S1.** Characteristics of the synthetic peptides in this study.

| *Strategy* | *Identity* | *[a] MW* | *[b] $t_R$ [min]* | *[c] RCY* | *[d] LogP* | *[e] $K_D$ (M)* |
|---|---|---|---|---|---|---|
| | GS | 2601.65 | 6.586 | 90.1% | -2.15 | 2.76E-09 |
| ***Multivalency*** | PEG-1 | 3415.60 | 8.966 | 58.8% | -1.67 | > 1E-06 |
| | PEG-2 | 5942.19 | 8.600 | 57.7% | -1.76 | > 1E-06 |
| | PEG-3 | 7414.34 | 8.733 | 90.43% | -1.73 | > 1E-06 |
| ***Cyclization*** | Cyc-1 | 3125.28 | 7.674 | 70.73% | -1.74 | 3.40E-06 |
| | Cyc-2 | 3125.28 | 7.828 | 71.79% | -1.86 | 1.75E-07 |
| | Cyc-3 | 2886.01 | 7.438 | 69.44% | -1.72 | 7.25E-07 |
| | Cyc-4 | 2972.10 | 7.214 | 77.28% | -2.06 | 1.55E-07 |
| ***Amino acid*** | GD | 2741.70 | 6.646 | 44.90% | -2.09 | 2.58E-09 |
| ***mutation*** | GE | 2811.84 | 6.210 | 66.00% | -2.60 | 2.74E-09 |
| | GR | 2947.21 | 6.413 | 49.30% | -1.49 | 1.44E-09 |
| | R | 2669.26 | 6.661 | 79.30% | -1.96 | 3.28E-09 |
| ***Covalent*** | Salk | 2655.70 | 6.649 | 86.73% | -2.04 | 1.71E-07 |
| | Scl | 2678.13 | 6.681 | 76.40% | -1.98 | 5.53E-08 |
| | Ssf | 2787.81 | 6.714 | 86.72% | -2.10 | 7.66E-08 |
| | Ralk | 3001.25 | 6.248 | 52.62% | -1.54 | 6.35E-07 |
| | Rcl | 3023.68 | 6.381 | 80.80% | -1.49 | 8.40E-07 |
| | Rsf | 3133.36 | 6.332 | 80.40% | -1.52 | 5.44E-07 |

[a] MW: molecular weight. [b] Retention time of all peptides, which was measured by HPLC. The solvent gradient was as follows: solvent A, aqueous solution with 0.1% trifluoroacetic acid (TFA); solvent B, acetonitrile with 0.1% TFA. The data were acquired over 20 min, with acetonitrile concentration increasing from 10% to 100%, and the flow rate was set to 1 mL/min. [c] Radiochemical yield (RCY) of all $^{68}$Ga-labeled peptides. The RCY was determined by radio-HPLC. The solvent gradient was as follows: solvent A, aqueous solution with 0.1% trifluoroacetic acid (TFA); solvent B, acetonitrile with 0.1% TFA. [d] LogP was determined by the 2480 Wizard auto γ-counter, and was measured in the n-butanol and water phases. [e] The $K_D$ of all peptides was measured through SPR using the Biacore. PD-L1 protein (20 μg/mL) was immobilized on the COOH-sensor chip.
